# Multi-output computation by single neuron biophysics in a visual system

**DOI:** 10.1101/2025.02.02.636137

**Authors:** H. Sebastian Seung

## Abstract

As long anticipated (Sandberg and Bostrom 2008; Seung 2012; Szigeti et al. 2014), connectomics is providing a new foundation for brain simulation by replacing theoretical assumptions about network connectivity with solid empirical facts. Connectomics also yields detailed information about neuronal morphology, which is useful for simulating the biophysics of single neurons (Yang et al. 2016; Meier and Borst 2019; T. X. Liu et al. 2022). Here I introduce a formalism for simulating a brain as a network of synapses interacting via an effective resistance matrix. By computing this matrix for fly visual interneurons, I find evidence that some neurons may be true multi-output devices, neither well approximated as “point neurons” (Lappalainen et al. 2024; Shiu et al. 2024), nor as collections of functionally independent compartments (Meier and Borst 2019). Within a linear approximation, such a neuron is instead equivalent to a hierarchy of virtual neurons that spatially pool over multiple length scales. The computational powers of multi-output neurons may support highly sophisticated normalizations in the fly visual system (Seung 2024a).

---

Neural network models are traditionally based on the assumption that a neuron carries a single output signal, a simplification that has been justified in two ways. Textbook accounts tell us that propagation of the action potential produces a stereotyped voltage signal (“spike”) at all output synapses on the axon. Alternatively, if a neuron is electrotonically compact, its transmembrane voltage is uniform across its output synapses even in the absence of spiking. In either case, the multiple output synapses of an axon can be approximated as broadcasting a single output signal, the scalar-valued “activity” of the neuron. This assumption is implicit in most network models in theoretical neuroscience, and recent fly brain simulations are no exception (Lappalainen et al. 2024; Shiu et al. 2024).

It is already known that some fly neurons are neither spiking nor electrotonically compact (Yang et al. 2016; Meier and Borst 2019; Amin et al. 2020; Schenk and Gaudry 2023). The importance of such empirical findings for function remain speculative. Perhaps such neurons are minor or rare exceptions that can largely be ignored when understanding brain function, a viewpoint taken by recent fly brain simulations (Lappalainen et al. 2024; Shiu et al. 2024). Or perhaps true multi-output neurons are common in the fly brain, and possess nontrivial computational powers that require more faithful simulation.

This is the context for the biophysical simulations of single neurons presented here. Based on connectomic data, I compute, analyze, and interpret the effective resistance matrix *R*_*ij*_, which relates the presynaptic voltage V_*i*_ at synapse *i* to the postsynaptic current *I*_*j*_ due to synapse *j*. The linear approximation

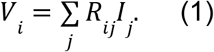

is familiar from passive models of dendrites of vertebrate neurons. Voltages are measured relative to the resting potential defined for zero synaptic currents. Here Eq. (1) is applied to the entirety of a fly neuron’s arbor, which may be a reasonable approximation if the neuron is nonspiking (Yang et al. 2016; Meier and Borst 2019).

Eq. (1) ignores many known aspects of single neuron biophysics, and its applicability is restricted to quasi-steady state phenomena. Despite these and many other limitations discussed later on (Uncertainties and Limitations), I focus on Eq. (1) because it is the basis for a “synapse network” formalism (Mathbox) that could replace all or part of the traditional neural networks used in fly brain simulation (Lappalainen et al. 2024; Shiu et al. 2024). A neuron in a synapse network generically carries multiple distinct output signals, the voltages associated with its output synapses. The synapse network formalism (Eq. 1 and Mathbox) should be viewed as a compromise: complex enough to model multi-output neurons, yet simple enough to be tractable.

If a neuron is electrotonically compact, then *R*_*ij*_ = const for all output synapses *i* and input synapses *j* of the same neuron. In this “point neuron” limit, *V*_*i*_ is the same for all output synapses of neuron *i*. Assuming point neurons, a synapse network reduces to a traditional neural network (Mathbox). Viewed abstractly, synapse networks are computationally equivalent to neural networks. But the two model classes ascribe differing computational powers to a single neuron.

I first show how to compute *R*_*ij*_ from the shapes of neurons that are reconstructed in exquisite detail by connectomics (Zheng et al. 2018; Dorkenwald et al. 2024; P. Schlegel et al. 2024; Matsliah et al. 2024). Second, the effective resistance matrix is analyzed for some fly visual neurons to understand how they deviate from the point neuron approximation. Finally, a hierarchical decomposition of the effective resistance matrix provides a way of interpreting the visual function of these neurons, and also a route to efficient simulation of synapse networks.

## Dm interneurons are potentially multi-output devices

Dm cells of the fly visual system are defined as neurons that are completely confined to the distal medulla, a neuropil of the optic lobe (Fischbach and Dittrich 1989; A. Nern, Pfeiffer, and Rubin 2015; Matsliah et al. 2024). Dm types are amacrine, meaning that output synapses are distributed throughout Dm arbors, rather than concentrated in some axonal compartment (Fig. 1a-c). An amacrine cell must be regarded as having multiple distinct outputs, if there is any violation of the point neuron approximation. If on the other hand, the output synapses were confined to some axonal compartment, they could experience a relatively uniform voltage, even if there were nonuniformity elsewhere. Therefore Dm cells are an appropriate testbed for exploring the implications of multi-output neurons.

**Figure 1.**
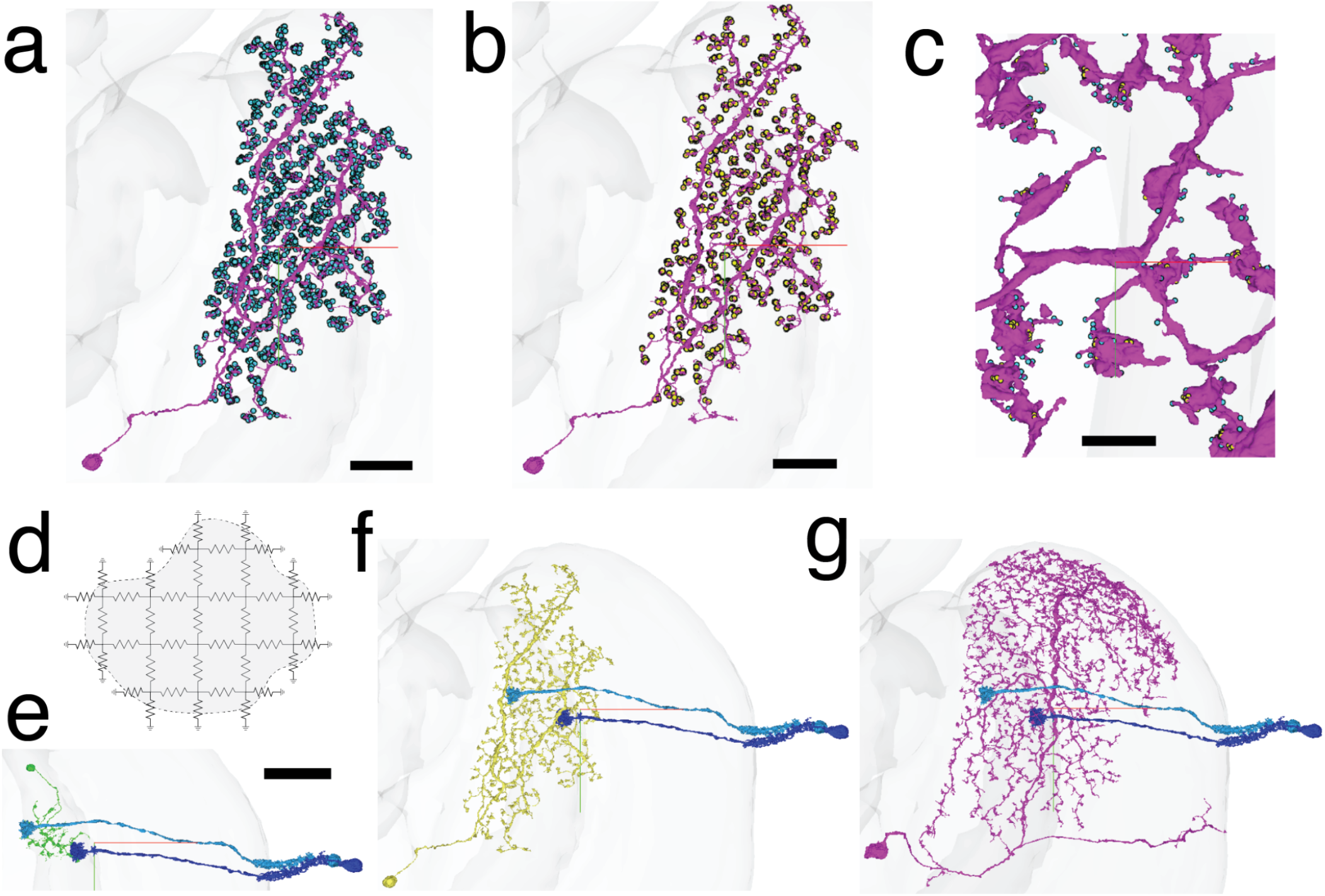
Dm interneurons as multi-output devices simulated by resistive networks. **a, b**, Dm6 neuron (magenta), input (cyan) and output (yellow) synapse locations. Since output synapses are widely distributed, the neuron will be a multi-output device, if it is nonspiking and the transmembrane voltage varies significantly across the neuron. **c**, Zooming in to the same Dm6 neuron shows its complex shape, which may not be so easily captured by traditional cylindrical approximations. Input (cyan) and output (yellow) synapses may be located at thinner or thicker regions of neurites. **d**, A neuron is modeled as a 3D grid of resistors based on its detailed 3D reconstruction from electron microscopic images. A 2D cross section of a 3D model is cartooned. The cross section of a neurite (gray) is bounded by the plasma membrane (dashed). The resistances are determined by the specific resistivity of the cytoplasm and plasma membrane, as well as the lattice constants. **e, f, g**, Dm15 (green), Dm6 (yellow), and Dm17 (magenta) cells. An example pair of L2 cells (light and dark blue) is reciprocally connected with all Dm cells. Scale bars: 20 (**a**,**b**), 3 (**c**), and 30 (**e, f, g**) µm. Crosshairs (red=lateral, green=ventral) are at the same location in **a**-**c**, and **e**-**g**.

## Passive electrical modeling of neurons

Reconstructing a fly brain from electron microscopic images (Zheng et al. 2018) yields synaptic connectivity between neurons, which has been used to constrain the connectivity between model neurons in fly brain simulations (Lappalainen et al. 2024; Shiu et al. 2024). But an electron microscopic reconstruction yields information beyond connectivity, which can also be leveraged by simulations. The detailed 3D shapes of all neurons and the locations of all synapses are determined during the reconstruction, and can be used to model the electrical behavior of a neuron.

Traditionally, this was done by approximating a neuronal arbor as cylinders or tapered cylinders of passive membrane, organized into a branching tree structure (Rall et al. 1992). While some parts of Dm neurons do resemble cylinders, other parts do not (Fig. 1c). The cylindrical approximation can be replaced by the reconstructed shapes of neurons. The resulting passive model is far more complex, but still tractable for contemporary computers.

I downsampled the FlyWire reconstructions of Dm neurons to near isotropic (32×32×40 nm^3^) voxel sizes. These voxels are small enough to accurately represent neuronal geometries. At the same time, a voxel is still large enough to contain hundreds of sodium ions and thousands of potassium ions, so neglecting the discreteness of charge is a reasonable approximation. Also the voxel size is much larger than the Debye length, so the ohmic approximation is reasonable.

I modeled each Dm cell as a 3D grid of resistors between nearest neighbor nodes located at voxel centers (Fig. 1d, Methods). Resistances between neighboring nodes were determined by the resistivity of the cytoplasm and the voxel dimensions. Resistances between nodes and the extracellular space were set by the specific transmembrane resistivity and the voxel dimensions. The extracellular space was assumed to be equipotential.

The currents flowing into the nodes are related to the voltages at the nodes by a linear system of equations involving a sparse matrix of conductances (Methods). By solving this sparse linear system of equations, one can calculate the nodal voltages of the neuron for a given configuration of nodal currents. Equivalently, inverting the conductance matrix yields an effective resistance matrix relating the nodal voltages to the nodal currents. These matrices are large, with dimensionality equal to the number of nodes in a neuron. The simulations to be presented span a range of 2 to 38 million nodes. Although the conductance matrix is sparse, the effective resistance matrix is dense, so it is impractical to compute all of its elements. As it turns out, it is sufficient to compute a much smaller submatrix including only the nodes at which synapses are located, as described below.

## Modeling synapses

While the resistive grid has a large number of nodes, only a small subset function as inputs and outputs. A neuron receives *N*_in_ input synapses and sends *N*_out_ output synapses. The synapse locations are provided by FlyWire (Fig. 1a-c), and are used to designate some nodes as inputs and outputs. An input synapse to a neuron is modeled as injecting current into the node at the location of the input synapse. For limitations of this current source model, see Uncertainties and Limitations. The signal carried by an output synapse is identified with the voltage of the node at the output synapse location (Mathbox). The input-output relationship for the neuron is specified by the *N*_out_ × *N*_in_ matrix *R* in Eq. (1). This can be obtained by repeatedly solving the linear equations for the “impulse response” to injecting current into a single input node.

## Mapping to visual space

The dominant inputs and outputs to a Dm type are generally modular cell types (Seung 2024a). A modular cell type forms a hexagonal lattice (Takemura et al. 2015; Matsliah et al. 2024), with cells that can be placed in one-to-one correspondence with image hexels (hexagonal pixels) sensed by the ommatidia of the compound eye.

To visualize the effective resistance matrix, it is helpful to map synapses to hexels. Let *P*_*I*_ denote the set of output synapses onto the cell of the target hexel type at location *I*, and *Q*_*J*_ the set of input synapses from the cell of the source hexel type at location *J*. Then one can define an averaged form of the effective resistance matrix indexed by locations on the hexagonal lattice,

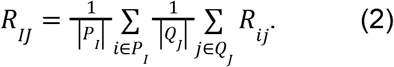

Rows and columns of this matrix will be visualized as images on the hexagonal lattice in subsequent figures.

## Simulations of small, medium, and large Dm cells

The above formalism will be applied to three Dm cells of three different types. The three types (Dm15, Dm6, and Dm17) are all strongly reciprocally connected with L2 cells (Fig. 1e-g), meaning that L2 is both the strongest input and strongest output of all three types (Seung 2024a).

Individual L2 cells are confined to single hexels and do not overlap with each other. Individual Dm cells, on the other hand, extend over multiple hexels, and the cells of a Dm type typically overlap with each other.

Most Dm interneuron types are thought to be inhibitory (Methods). Elsewhere I have argued that they serve the function of spatial normalization (Seung 2024a). Each of these three Dm types enables an L2 cell to compare its activity with the pooled activities of neighboring L2 cells.

Dm cells vary over a large range of sizes. Dm15 is small (Fig. 1e), Dm6 is medium (Fig. 1f), and Dm17 is large (Fig. 1g). Naively, one might expect that each type spatially pools L2 activity in neighborhoods of different sizes, enabling normalization over different length scales. The simulations below will show how this naive picture must be revised if the point neuron approximation breaks down.

As a preliminary, it is helpful to consider the intuitions from cable theory. Injecting current into a cable should result in a voltage that decays exponentially with distance from the injection. The length constant of the decay is 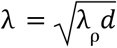, which is the geometric mean of the neurite caliber (diameter) *d* and a length scale 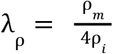 set by the ratio of the specific membrane resistivity *ϱ*_*m*_ and the cytoplasmic resistivity *ϱ*_*i*_. A biological neuron has a more complex shape than a cable, but this formula encapsulates the intuition that violations of the point neuron approximation become likely when neurites with small caliber extend over long distances.

I will use two parameter sets drawn from the literature (Methods). Gouwens and Wilson used *λ*_ρ_ = 20 cm to model electrophysiological recordings of a neuron in the fly olfactory system (Gouwens and Wilson 2009). Computing the geometric mean with a diameter of 0.4 μm yields the electrotonic length *λ* = 280 μm, and with a diameter of 0.1 μm yields *λ* = 140 μm.

Meier and Borst used *λ*_ρ_ = 5 cm to model calcium imaging of a non-columnar neuron in the fly visual system (Meier and Borst 2019). This yields an electrotonic length of 140 μm for a diameter of 0.4 μm, and 70 μm for a diameter of 0.1 μm. Similar parameters were also used to model voltage imaging of a columnar visual neuron (Yang et al. 2016).

## A small Dm cell

The above computational procedures were applied to a Dm15 cell. The terminals of the two L2 cells were located on the left and right sides of the Dm15 arbor (Fig. 1e). To simulate the effects of L2 activity on the Dm15 cell, current was injected into the Dm15 cell at the locations of the synapses from each L2 cell (Fig. 2a, b), and the voltages were computed by solving the sparse linear set of equations.

**Figure 2.**
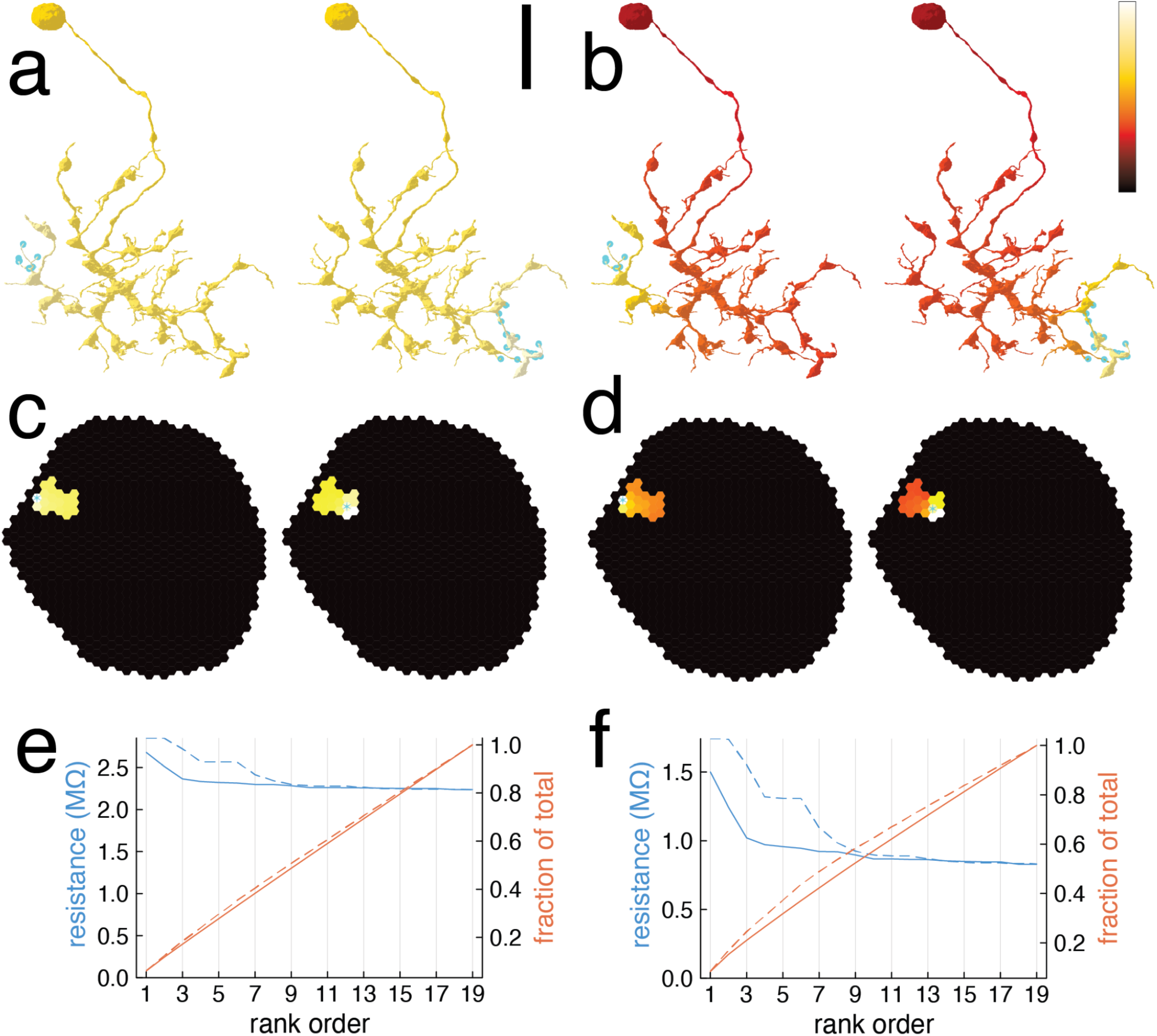
Effective resistance of a small Dm cell. Dm15 is a small-sized member of the Dm interneuron family. The attenuation of voltage with distance from the stimulated location is relatively minor, so the point neuron is a reasonable approximation. The simulation involves solving a linear system of equations for 2 million nodal voltages. **a**,**b**, Simulated Dm15 voltage (colormap) caused by activating input synapses (cyan) from L2 cells (Fig. 1e). The voltage is almost uniform for Gouwens-Wilson parameters (**a**), but attenuates noticeably for Meier-Borst parameters (**b**). The voltage is per unit of input current, and so has the units of resistance. **c**,**d**, Dm15 voltages of (**a**,**b**) mapped to visual locations in the hexagonal lattice of ommatidia in the compound eye. The color of each hexel represents the mean voltage at all the output synapses onto the L2 cell located at that hexel. The voltage attenuates with distance from the input L2 cell (cyan star) for Meier-Borst parameters (**d**), but the attenuation is weak for Gouwens-Wilson parameters (**c**). **e**,**f**, Dm15 voltages of (**c, d**) after rank ordering. The maximal effective resistance is near the stimulated location, and is higher for the right L2 cell (dashed) than for the left L2 cell (solid). The minimal effective resistance is far from the stimulated location, and depends negligibly on the location of stimulation. In both cases the cumulative sum of the resistances (red) is roughly linear. Scale bar: 10 µm (**a, b**). Color bar (**a, b**) runs from 0 to maximum resistance values in (**e, f**).

For Gouwens-Wilson parameters, the voltage varies little across the arbor (Fig. 2a), so the point neuron approximation is quite good. For Meier-Borst parameters, the voltage declines noticeably at locations distant from the stimulus (Fig. 2b), so the point neuron approximation is not as good.

The voltages in Figs. 2a and b are per unit of injected current, so that the depicted quantity is actually effective resistance rather than voltage. (For each L2 cell, the unit of injected current is divided equally across all its synapses onto the Dm15 cell.) Effective resistance focuses on intrinsic properties of the Dm15 cell, and ignores synaptic properties like the numbers and unitary strengths of L2 synapses received by the Dm15 cell.

The simulation yields the voltages at all locations in the cell. The effective resistance matrix of Eq. (1) is obtained by selecting the voltages corresponding to the locations of output synapses. As mentioned earlier, for visualization purposes, it is helpful to map the rows or columns of the effective resistance matrix onto the eye. Since L2 is a hexel type, every L2-Dm15 or Dm15-L2 synapse can be located on the hexagonal lattice of ommatidia or medulla columns. This is the resistance matrix as a function of space, and two of its columns are shown in Figs. 2c and 2d. All columns are shown in Supplementary Data 1.

For Meier-Borst parameters (Fig. 2d), a “hot spot” is apparent around the input location. While the voltage is lower at distal locations, there is not much variation outside the hot spot. If the voltages are graphed after rank ordering (Fig. 2f), they form an almost constant plateau outside the hot spot. One could say that the point neuron approximation holds, except for a boost near the stimulus location.

For Gouwens-Wilson parameters (Fig. 2c, e), the behavior is qualitatively similar, but there is little difference between the hot spot and the plateau.

## A medium Dm cell

A Dm6 cell is considerably larger than a Dm15 cell (Fig. 1f). The simulated Dm6 cell receives inputs from the same L2 cells (Fig. 1f) that were Dm15 inputs in Figs. 1e and 2. Here a hot spot is apparent near the stimulus location (Fig. 3a, b), for either parameter set. However, the behavior is more complex than for the Dm15 cell. If the hot spot is excluded, the response is approximately piecewise constant in left and right domains of the arbor. Both regions correspond to subtrees of the arbor, and are elongated along a (roughly) vertical axis. The response attenuates more along the horizontal axis, and less along the vertical axis. The domain structures are easier to see when the voltage responses are mapped to locations in the visual field (Fig. 3c,d).

**Figure 3.**
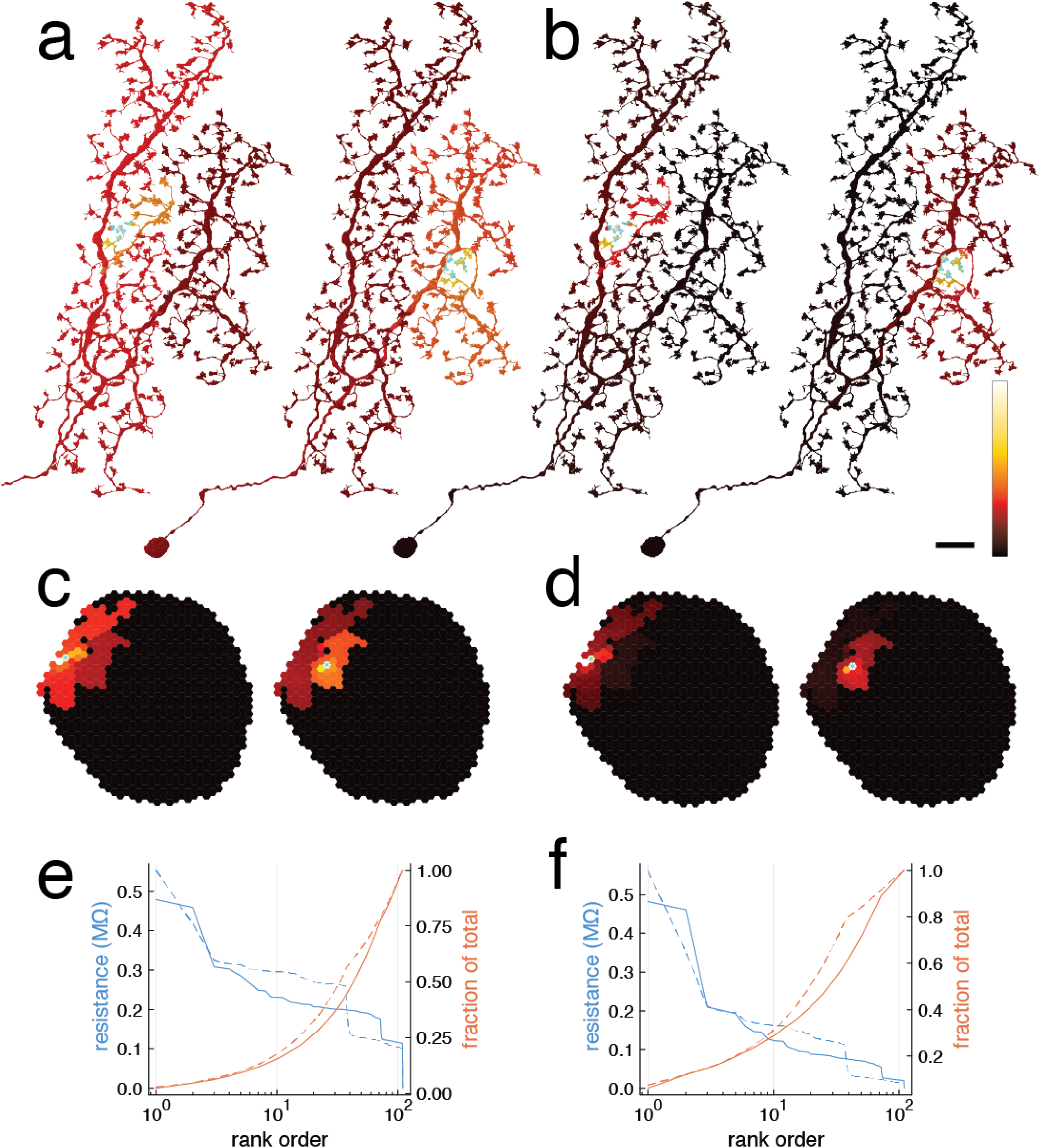
Effective resistance of a medium Dm cell. Dm6 is a medium-sized member of the Dm interneuron family. More attenuation of voltage with distance is seen, but the amount of attenuation depends on resistivity parameters. The simulation involves solving a linear system of equations for 24 million nodal voltages. **a, b**, Simulated Dm6 voltage (colormap) caused by activating input synapses (cyan) from L2 cells (Fig. 1f). For Gouwens-Wilson parameters (**a**), the voltage attenuates weakly along the vertical axis and strongly along the horizontal axis. For Meier-Borst parameters (**b**), both attenuations are stronger. For both parameter sets, a hot spot is seen right at the stimulated location. **c**,**d**, Dm6 voltages of (**a**,**b**) mapped to visual locations. The color of each hexel represents the mean voltage at all the output synapses onto the L2 cell located at that hexel. Apart from a hot spot at the stimulated location (cyan star), the voltage looks approximately piecewise constant in two spatial domains, a larger domain near the border of the eye, and a smaller domain in the interior of the eye. The response of the distal domain is weak for Gouwens-Wilson parameters (**c**) and almost imperceptible for Meier-Borst parameters (**d**). **e**,**f**, Dm6 voltages of (**c, d**) after rank ordering. The maximal effective resistance is near the stimulated location, and is higher for the right L2 cell (dashed) than for the left L2 cell (solid). The spatial domains in (**c**,**d**) correspond with the plateaus in the staircase-like shapes of the plots. Although the distal domain contains low resistance values, it can contribute as much as half the spatial sum of the resistances (compare blue and red dashed) for Gouwens-Wilson parameters (**c**), but the contribution is much less for Meier-Borst parameters (**d**). Scale bar: 10 µm (**a, b**). Color bar (**a, b**) runs from 0 to maximum resistance values in (**e, f**).

Just two columns of the effective resistance matrix are shown in Fig. 4b. A fuller view is provided in Supplementary Data 2, which exhaustively displays all columns of the effective resistance matrix. There is considerable variability across different input locations.

**Figure 4.**
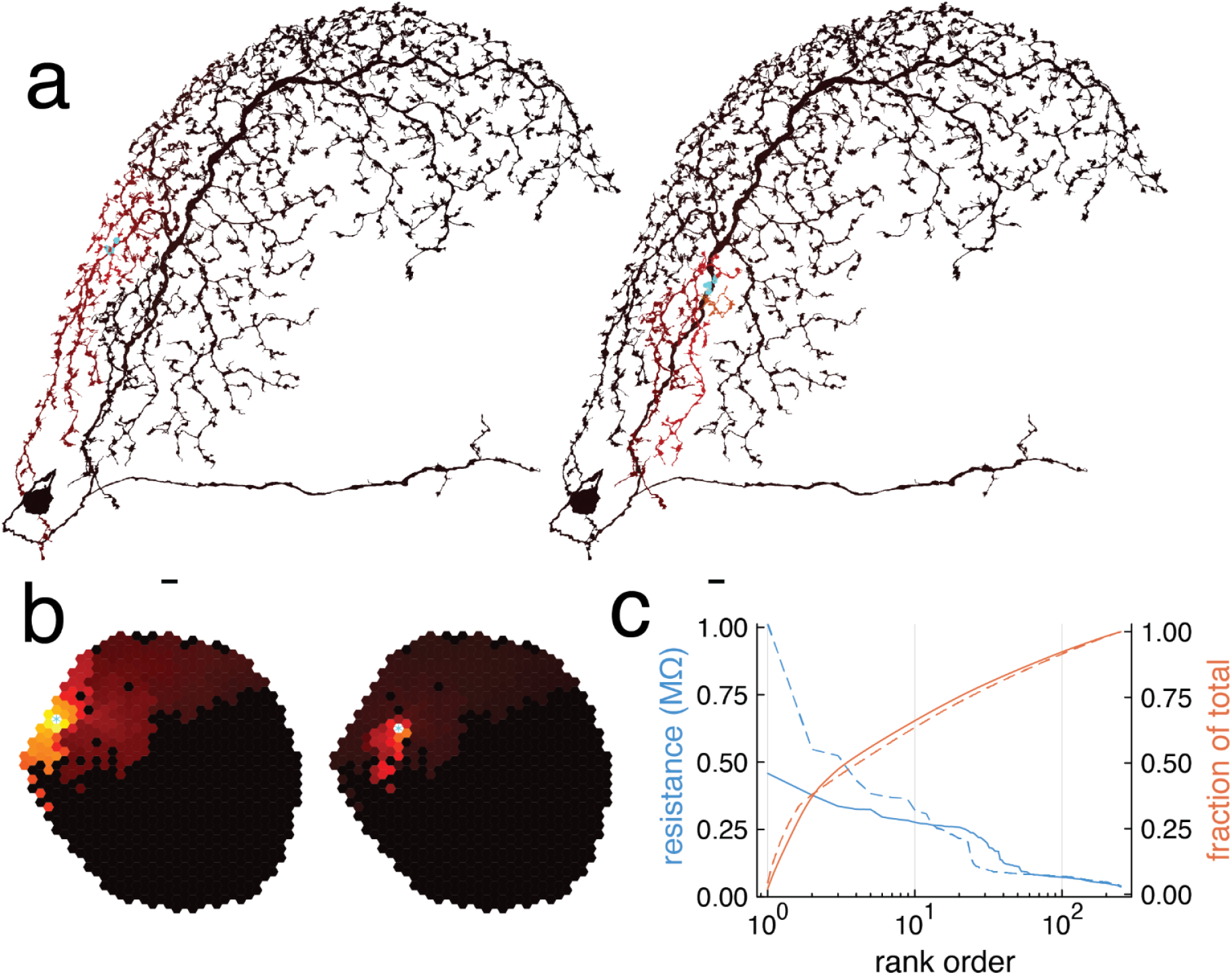
Effective resistance of a large Dm cell. Dm17 is a large member of the Dm interneuron family, covering a substantial fraction of the entire visual field. Strong attenuation of voltage with distance is observed; the point neuron approximation is clearly violated. The simulation involves solving a linear system of equations for 38 million nodal voltages. Gouwens-Wilson parameters are shown here, and Meier-Borst parameters in Extended Data Fig. 2. **a**, Simulated Dm17 voltage (colormap) caused by activating input synapses (cyan) from L2 cells (Fig. 1g). A voltage response is visible over only a small fraction of the arbor. **b**, Dm17 voltages of (**a**) mapped to visual locations. The color of each hexel represents the mean voltage at all the output synapses onto the L2 cell located at that hexel. The response to the left L2 input looks broader than the response to the right L2 input. Some domain structure like that of Fig. 3 might exist, but is more complex. **c**, Dm17 voltages of (**b**) after rank ordering. Attenuation is stronger for the right L2 input (dashed) than for the left L2 input (solid). A substantial fraction of the total summed resistance can be contributed by the weakly activated hexels (red).

For more quantitative information about attenuation, the voltages are graphed in Figs. 3e,f after rank ordering. Two plateaus are evident outside the hot spot, corresponding to the two domains. There is greater attenuation from hot spot to proximal domain to distal domain for Meier-Borst parameters (Fig. 3f) than for Gouwens-Wilson parameters (Fig. 3e). For Meier-Borst parameters, the voltage in the distal domain is so attenuated that it is barely visible in Fig. 3d.

## A large Dm cell

A Dm17 cell is even larger than a Dm6 cell, and covers a significant fraction of the visual field (Fig. 1g). Stimulation of an L2 cell produces a complex response (Fig. 4a). There is a hot spot near the location of the stimulated L2 cell. Neighboring locations also exhibit some voltage response. But most of the arbor appears to have little or no response in Fig. 4a. This strong attenuation is observed for Gouwens-Wilson parameters (Fig. 4a), and is even stronger for Meier-Borst parameters (Extended Data Fig. 2). Either way, the point neuron approximation seems completely inapplicable to the Dm17 cell.

If the effective resistance is mapped onto visual locations (Fig. 4b), a faint voltage response becomes visible at the distal locations in the Dm17 arbor that seemed completely inactive in Fig. 4a. Two columns of the effective resistance matrix are shown in Fig. 4b, and all columns are provided in Supplementary Data 3.

Rank ordering the voltages reveals that the distal responses are weak indeed (Fig. 4c). It should be noted that the most distal voltages do not depend on stimulus location (Fig. 4c). The distal voltages only look different in Fig. 4b because the colormap maximum is normalized to the hot spot voltage.

## Point neuron model as rank one approximation

If a neuron were electrotonically compact, then all elements of its effective resistance matrix would be the same. In other words, the matrix would be proportional to the outer product of the vector of all ones with itself. Generalizing slightly, a point neuron can be defined as one for which the effective resistance matrix is an outer product of two vectors (Fig. 5a). One vector can be interpreted as the input connections of a point neuron, and the other vector as the output connections (Fig. 5b). Then the accuracy of the point neuron approximation can be quantified as the accuracy of a rank one approximation to the effective resistance matrix.

**Figure 5.**
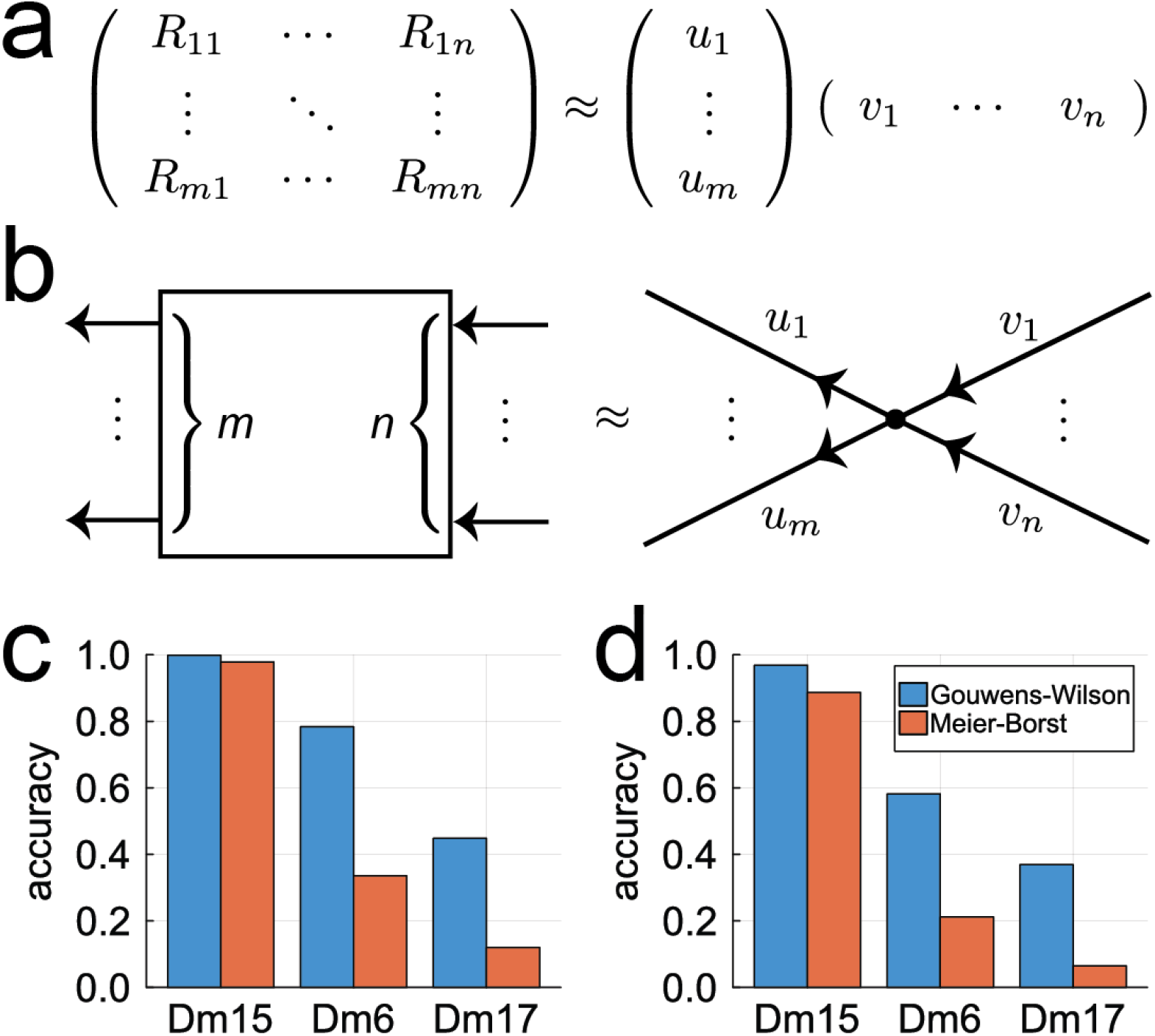
Accuracy of the point neuron approximation. Dm15 is well-approximated as a point neuron (≈90% accuracy or better), while Dm17 is not. For Dm6 the approximation might be tolerable, depending on parameters. **a**, The effective resistance matrix can be approximated as a rank one matrix, which is an outer product of two vectors. **b**, A neuron computes *m* outputs from *n* inputs. The rank one approximation can be interpreted as a device (point neuron) that computes a weighted sum of *n* inputs, and then broadcasts this scalar value to *m* outputs with a rescaling. **c**, A rank one approximation can be computed by truncating the singular value decomposition of the effective resistance matrix. Accuracy of the point neuron model is quantified by the ratio of the squared Frobenius norms of the rank one matrix and the effective resistance matrix. **d**, Alternatively, a rank one approximation can be obtained by replacing each row of the effective resistance matrix with its minimum value. This approximation is a lower bound for the effective resistance. Accuracy is quantified by the ratio of the summed elements of the rank one matrix and of the effective resistance matrix.

For example, a point neuron model can be constructed by truncating the singular value decomposition to have rank one. This is an excellent approximation for the Dm15 cell (98% accuracy or better), and is poor for the Dm17 cell (Fig. 5c). For the Dm6 cell, accuracy approaches 80% for Gouwens-Wilson parameters, but is much worse for Meier-Borst parameters.

Alternatively, a point neuron model can be constructed by replacing each row of the effective resistance matrix with its minimum value (Fig. 5d). This is also a rank one approximation, for which the residual error (difference between left and right sides of Fig. 5a) is nonnegative. Since a nonnegative matrix is decomposed into a sum of nonnegative matrices, accuracy can be evaluated by simply summing matrix elements, and taking ratios (Fig. 5d). This alternative measure of accuracy gives similar results as singular value decomposition (Fig. 5d). The same accuracy measure will be used again for the more sophisticated nonnegative decomposition that follows.

## Hierarchical decomposition of the resistance matrix

The rank one approximation is poor for Dm6 and Dm17. However, it turns out to be possible to construct a low rank approximation that can be interpreted as a hierarchy of virtual point neurons. This is done by decomposing the effective resistance matrix into the sum of nested block diagonal matrices.

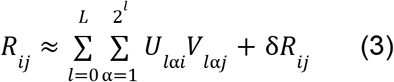

The block diagonal matrix at the *l*th level of the hierarchy contains 2^*l*^ blocks (Fig. 6a). The blocks at the *l*th level are a refinement of the blocks at the (*l*–1)st level. The computation of the decomposition is described in the Methods. Here only the result is described, and its properties can be understood without knowing how the decomposition was derived.

**Figure 6.**
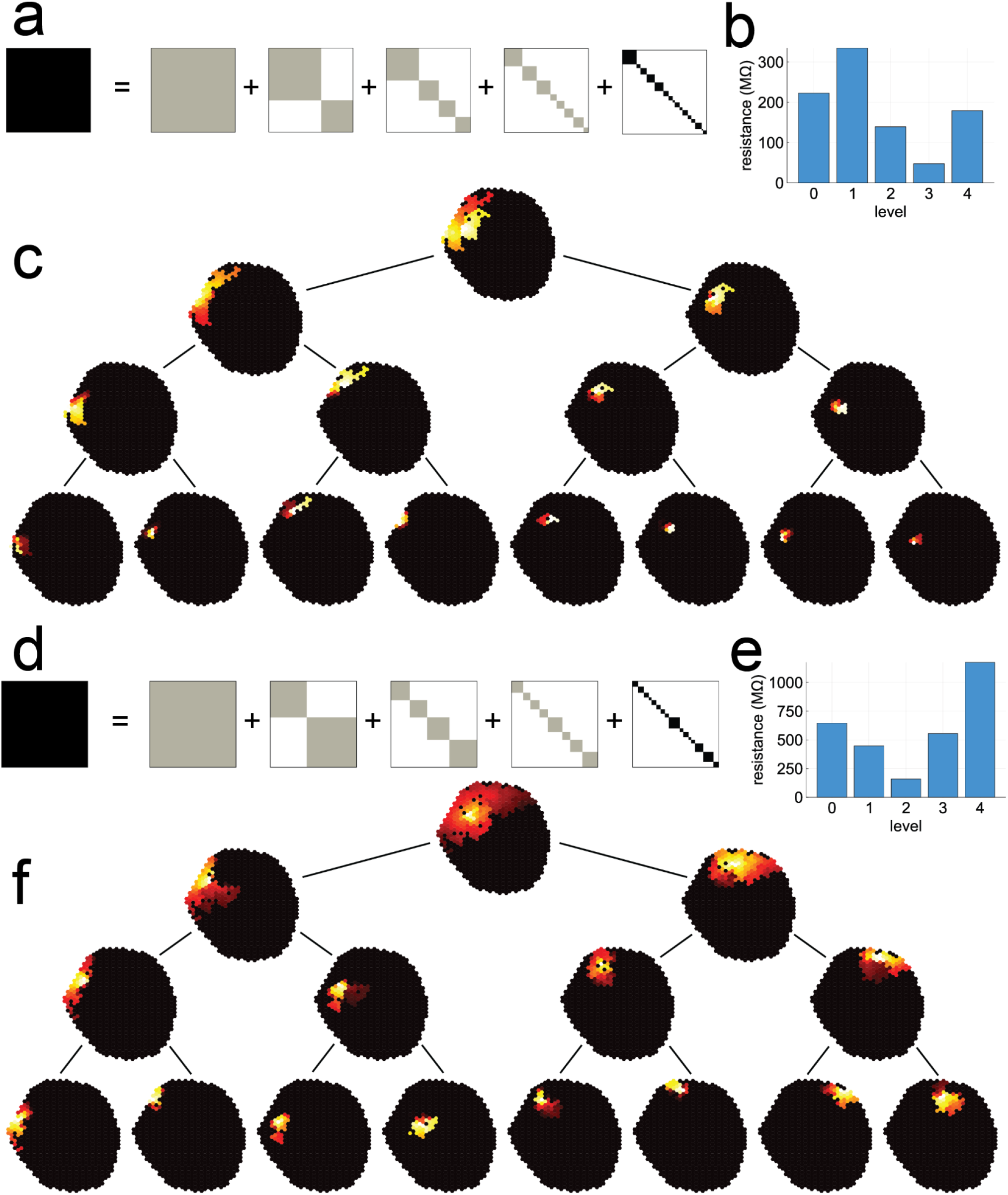
Decomposition into hierarchy of virtual point neurons. **a**, The resistance matrix for Dm6 with Meier-Borst parameters is decomposed into a sum of block diagonal matrices. Each gray square represents a block that is rank one, and can be interpreted as a virtual point neuron. Each set of blocks is a refinement of the previous set of blocks. **b**, How the Dm6 resistance matrix is distributed over the levels of the hierarchy. **c**, Input connections of the virtual neurons in the hierarchical decomposition of Dm6. Output connections look very similar. The virtual neurons at each level correspond to gray blocks in one of the matrices depicted in (**a**). **d**, The resistance matrix for Dm17 with Gouwens-Wilson parameters is decomposed into a sum of block diagonal matrices. **e**, How the Dm17 resistance matrix is distributed over the levels of the hierarchy. **f**, Input connections of the virtual neurons in the hierarchical decomposition of Dm17. The virtual neurons at each level correspond to gray blocks in one of the matrices depicted in (**d**).

This decomposition has two motivations. First, multiplying by the matrix *R* in Eq. (1) is time-consuming, because the matrix is dense. It requires *N*_out_ × *N*_in_ multiply-adds. A point neuron model is much faster to simulate, requiring just *N*_out_ + *N*_in_ multiply-adds. If the hierarchical decomposition of Eq. (4) is used, multiplying by *R* requires a number of operations that is *L*(*N*_out_ + *N*_in_) plus the number of elements in the residual matrix δ*R*. The latter number is small because δ*R* is a block matrix, and its blocks are a refinement of those in level *L* of the decomposition. Therefore the decomposition of Eq. (4) could speed up simulations of the synapse net.

The second motivation for the decomposition is interpretability. In levels 0 to *L*, each block is a rank one matrix, and therefore can be interpreted as a point neuron. Therefore levels 0 to *L* contain a hierarchy of virtual point neurons. The virtual neurons at the *l*th level have size that is roughly half that of the virtual neurons at the (*l*–1)st level, so the decomposition includes a hierarchy of length scales.

The residual matrix δ*R* also has a block diagonal form, but the 2^*L*+1^ blocks are not low rank. They can be interpreted as 2^*L*+1^ virtual multi-output neurons that are smaller than the original neuron. Alternatively, they can be interpreted as *N*_out_ virtual point neurons, one for each output synapse.

A decomposition with *L*=3 is shown for the Dm6 cell with Meier-Borst parameters. Summing the matrix elements at each level of the hierarchy shows that the effective resistance is distributed over levels, and not concentrated at a single level (Fig. 6b). The matrix factors *U* can be interpreted as the input connections for a set of virtual point neurons. They are visualized by mapping them to the eye (Fig. 6c). The matrix factors *V* can be interpreted as the output connections of the point neurons, and look similar (data not shown). The level 0 factor covers the entire arbor. Levels 1 through 3 cover domains of the arbor. The residual virtual neurons are shown in Supplementary Data 4. The decomposition includes a hierarchy of scales from the entire arbor down to small neighborhoods.

A decomposition with *L*=3 is shown for the Dm17 cell with Gouwens-Wilson parameters. The block diagonal matrices are shown in Fig. 6d. Again, the effective resistance is distributed over levels of the hierarchy (Fig. 6e). The matrix factors *U* are visualized for each virtual point neuron by mapping them to the eye (Fig. 6f). The residual virtual neurons are shown in Supplementary Data 5.

## Uncertainties and limitations

Even if the linear approximation of Eq. (1) is assumed accurate, there are parametric uncertainties regarding the specific membrane and cytoplasmic resistivities. Cells were simulated for two parameter sets drawn from the literature, which were meant to define a range of biologically plausible parameters (Gouwens and Wilson 2009; Yang et al. 2016; Meier and Borst 2019). The simulations suggest that the point neuron approximation is poor for Dm6 and Dm17. Therefore Dm6 and Dm17 might turn out to behave as true multi-output devices. An important caveat is that this expectation is based on the assumption that Dm cells are nonspiking.

## The possibility of spiking or hybrid electrical behaviors

Many columnar neurons in the fly visual system appear to be nonspiking (Behnia et al. 2014; Yang et al. 2016). Both columnar (Yang et al. 2016) and non-columnar (Meier and Borst 2019) neurons in the fly visual system have been successfully modeled as passive electrical devices. Dm interneurons might likewise turn out to be nonspiking. However, some *Drosophila* visual neurons are known to exhibit spiking, or some hybrid of action and graded potentials (Mu et al. 2012).

Action potentials in neurons are due to the presence of voltage-gated sodium channels, and Para is the relevant channel in *Drosophila* neurons. The biological evidence for or against Para channels in Dm cells is inconclusive at present (Methods). Dm cells might turn out to contain little or no Para, which would boost confidence in the passive electrical simulations presented here. At the other extreme, Para channels could turn out to be numerous and distributed throughout Dm cells. Then Para could support the propagation of an action potential throughout the arbor, and the passive formalism introduced here would be of limited utility.

In an intermediate scenario, Para could turn out to be concentrated in some spike initiation zone (SIZ) restricted to a small portion of the arbor (Ravenscroft et al. 2020). For amacrine cells, which lack axons, Para might be concentrated in the primary neurite. Some empirical evidence for this idea has been reported for local interneurons of the antennal lobe (Schenk and Gaudry 2023). Then spikes would initiate in the SIZ, and propagate to the rest of the neuron with passive attenuation. Outside the SIZ, attenuated spikes would be superimposed on graded potentials. One can potentially model such hybrid behavior using the effective resistance between input synapses and the hypothetical SIZ, but this is outside the scope of the present work.

### Calcium and voltage imaging

Given the preceding uncertainties, an important physiological experiment is to look for spiking in Dm interneurons. This could be done by patch clamping the cell body, or by voltage imaging of the cell body or neurites. A related experiment would probe the extent to which voltage changes are global versus local in the Dm interneurons. For example, an imaging experiment could reveal the voltage or calcium signals as a function of spatial location, in response to stimulation of a single ommatidium or a few neighboring ommatidia. To some extent, this is equivalent to measuring voltage or calcium at a single location, and varying the location of a spot stimulus in the visual field.

Visual responses of Dm9 (Heath et al. 2020; Schnaitmann et al. 2024), Dm8 (Pagni et al. 2021; Li et al. 2021), and Dm12 (Gür et al. 2024) have been observed by calcium imaging, but these are cells with relatively small arbors. If similar experiments were performed for larger cells like Dm6 and Dm17, they would reveal whether local stimulation evokes global responses that are strong everywhere, as one would expect if the cells were spiking, or local responses that decay rapidly, as one would expect if the cells were nonspiking. Spiking local interneurons of the antennal lobe exhibit global responses to local activation of a single glomerulus (Hong and Wilson 2015).

### Subthreshold nonlinearities

Even if the Dm cells turn out to be nonspiking, many subthreshold nonlinearities could be important. Synapses were modeled as current sources, which add linearly. In fact, the effects of synapses combine nonlinearly because postsynaptic currents are generated by changes in postsynaptic conductance. The current source model is a reasonable approximation for excitatory synapses, which have reversal potentials that are well-separated from the operating voltage range of the neuron. But inhibitory synapses often have reversal potentials that are at or near the resting potential of the neuron. Such shunting inhibition has the potential to be highly nonlinear (Silver 2010), though there is some contention about whether such nonlinearity is strong in real neurons (Holt and Koch 1997). Modeling shunting inhibition would be an important extension to the present formalism.

Many intrinsic conductances exhibit voltage dependence, even if they are not involved in the action potential. These could also lead to subthreshold nonlinearities.

### Realism in modeling

The detailed shapes of cells as reconstructed from electron microscopic images were used to construct passive electrical models. While connectomic reconstructions give unprecedented detail concerning the 3D shapes of neurons, they are not perfectly accurate. Conventional sample preparation techniques for electron microscopy are known to cause swelling of cells and shrinkage of extracellular space (Pallotto et al. 2015), and may fail to preserve “pearls-on-a-string” morphology of axons (Griswold et al. 2025). Other reconstruction artifacts may emerge during electron microscopic image acquisition and processing.

Nevertheless, the present work has illustrated that biophysical models of single cells are no longer restricted to the traditional descriptions of cells as assemblies of cylindrical segments (Rall et al. 1992). No claim is made that the approach as implemented here is more accurate than the traditional models. However, the present work can serve as a starting point for incorporating even more realism. For example, it is possible to reconstruct mitochondria from the same EM images, as has been done in other connectomic datasets (Turner et al. 2022). It would be more accurate to exclude mitochondrial volume from the electrical model of the intracellular space, and many of the thicker portions of the Dm arbor contain mitochondria.

It will become possible to constrain the electrical properties of the model based on the ion channels identified in Dm neurons by transcriptomics and proteomics. Transcriptomic studies currently cover some Dm types (Davis et al. 2020; Kurmangaliyev et al. 2020; Özel et al. 2021), but not yet Dm15, Dm6, and Dm17. In the long run, proteomics could eventually achieve the spatial resolution required to constrain the distribution of ion channels across the neuron.

In the present work, the specific resistivities were assumed to be the same throughout the neuron, but they might actually vary with location. Furthermore, the nodes of the model are so small that individual transmembrane resistances should be set stochastically by scattering individual ion channels across the neuronal membrane. Some preliminary simulations suggest that the results are qualitatively similar (data not shown). Further improvements in realism would require the modeling of calcium concentration as well as voltage at every node, and the addition of voltage-gated conductances.

### Dynamics

The steady state model presented here gives some insights into the attenuation of voltage across a Dm arbor. Generalizing to a full dynamical model might seem trivial from the theoretical viewpoint, as a passive electrical system is linear. However, practical implementation is nontrivial because the linear system is extremely stiff. It is well-known that traditional compartmental models of neurons are stiff, and the models introduced here are even more stiff because the number of variables has exploded. Dynamical simulations are outside the scope of the present work, and will be studied elsewhere. The hierarchical decomposition introduced in Figure 6 seems related to domain decomposition techniques in partial differential equations, which will be important for integrating the differential equations in a dynamical simulation. Addition of reaction-diffusion dynamics to voltage dynamics is another challenge at the frontier of current research, which aims to extend biophysical models of neurons to the nanoscale (Chen et al. 2022). The present work is a version of this challenge that is simplified enough to be employed in large-scale brain simulations, yet should also be rich enough to capture some of the biophysical realities of *Drosophila* neurons.

## Discussion

A true multi-output neuron deviates from the traditional conception that a neuron broadcasts a single scalar-valued signal to all output synapses. The possibility of multi-output neurons in the fly visual system was investigated through passive electrical modeling. Neuronal shapes were taken from electron microscopic reconstructions. Resistivity parameters were borrowed from previous studies that fit passive models of fly neurons to physiological measurements (Gouwens and Wilson 2009; Yang et al. 2016; Meier and Borst 2019).

Dm cells of small, medium, and large sizes were simulated. The effective resistance matrix of Eq. (1) was computed for each cell. The small Dm cell conformed well to the point neuron approximation (Figs. 2, 5), but the medium (Fig. 3) and large (Fig. 4) Dm cells deviated considerably (Fig. 5). For the medium and large cells, the effective resistance matrix was decomposed into a sum of block diagonal matrices. The blocks were organized into a hierarchy of layers. Each block was interpreted as a virtual point neuron, except the blocks in the lowest layer, which could be interpreted as either multi-output or point neurons.

### Receptive field vs. pooling field

The receptive field of a visual neuron was classically defined as “the region of the retina which must be illuminated in order to obtain a response” (Hartline 1938). In principle, it is possible for output synapses of a single neuron to have different receptive fields. This is not the case for my simulations of a small neuron (Dm15), for which the point neuron approximation is highly accurate (Fig. 5). All output synapses are expected to share the same receptive field, which spans all locations covered by the arbor (Fig. 2).

For the medium (Dm6) cell, predicting receptive fields from simulations is more subtle because the effective resistance, while nonzero, can be very small at many locations covered by the arbor. Illuminating such locations seems unlikely to drive detectable activity at the output synapse. Therefore, thresholding the effective resistance at some value could be a way of predicting receptive fields. If the threshold value is (somewhat arbitrarily) set to about 20% of the maximum, a Dm6 receptive field is predicted to contain ≈100 hexels for Gouwens-Wilson parameters (Fig. 3e), but only ≈10 hexels for Meier-Borst parameters (Fig. 3f). (The data shown in Fig. 3 is actually for columns of the effective resistance matrix or “projective fields”, but the data for rows or “receptive fields” is similar.)

While the receptive field of an output synapse is predicted to be smaller than the Dm6 arbor for Meier-Borst parameters, locations outside the receptive field could still potentially influence activity of the output synapse. Fig. 2f shows that the weaker elements of the effective resistance matrix still contribute a substantial fraction because they are so numerous. Therefore one might define a “pooling field” for the Dm6 cell that spans its entire arbor.

For the large (Dm17) cell, setting the threshold value at 20% of maximum results in a predicted receptive field that is much smaller than the arbor for either parameter set (Fig. 4c, Extended Data Fig. 2c).

To summarize, Dm6 and Dm17 may simultaneously exhibit small receptive fields and large pooling fields. The latter are expected to be weak and diffuse, but could give rise to significant effects when integrated over the large arbor areas. Therefore pooling fields could be physiologically detectable through manipulations of background luminance and contrast.

### Hierarchical computation

The dichotomy between receptive field and pooling field is simplistic. The hierarchical decomposition of Fig. 6 paints a more complex picture. There may be multiple levels of pooling fields corresponding to groups of virtual point neurons in a hierarchy. The pooling may be broadly distributed over multiple length scales (Fig. 6b,e), so that a single neuron may implement multi-scale pooling.

Hierarchical decomposition can also be compared with the simpler concept of “extreme compartmentalization.” In CT1, the largest amacrine cell of the fly visual system, electrical signals from one hexel propagate very little to other hexels (Meier and Borst 2019). In a recent simulation of the fly visual system, the CT1 cell was modeled by chopping it into many virtual point neurons, one for each hexel (Lappalainen et al. 2024). In this way, the multi-output nature of the CT1 cell was easily eliminated due to extreme compartmentalization. Visual neurons in other species have also been conceptualized in a similar manner. For example, the neurites of the starburst amacrine cell in the mammalian retina have been modeled as independent functional units (Euler, Detwiler, and Denk 2002; Hausselt et al. 2007). The hierarchical decomposition introduced here should be more broadly relevant for multi-output neurons, because it is applicable even when neurons are not extremely compartmentalized.

Previous conceptions of hierarchical computation by mammalian neurons have emphasized nonlinear and/or active dendrites (Poirazi, Brannon, and Mel 2003; Stuart, Spruston, and Häusser 2016). The present work shows that even linear and passive dendrites can exhibit rich computational behaviors when neurons are multi-output devices.

### Convolution by a single neuron

In the fly visual system, many cell types can be regarded as analogous to feature maps in a convolutional net. The neurons of such cell types perform the same computation at different locations in the visual field, because connectivity is approximately translation invariant. Convolutional kernels can be extracted from the wiring diagram of synaptic connectivity (Lappalainen et al. 2024; Seung 2024b).

While a convolution is computed by a population of neurons in a convolutional net, it is also theoretically possible for a single neuron to compute a convolution, at least in one dimension. The steady state voltage of an infinite cable is the convolution of the injected current with an exponential kernel. This construction does not obviously generalize to 2D, because a neuron has to cover 2D space with neurites that are effectively 1D. Nevertheless, it might be possible for a single neuron to approximate a 2D convolution.

For Dm6 and Dm17, the effective resistance matrix is clearly not convolutional, as violations of translation invariance are prominent and interactions are nonlocal (Figs. 3, 4, Supplementary Data 3-6). The residual matrix remaining after hierarchical decomposition is quite local (Supplementary Data 7-8), and comes closer to having convolutional structure.

### Applicability to brain simulation

Eq. (1) defined an effective resistance matrix for the synapses of a single neuron. This is a simplistic model with many limitations (Uncertainties and Limitations), if viewed from the field of single neuron biophysics. But from the viewpoint of brain simulation, Eq. (1) is much more complex than the point neurons in traditional neural network models (Lappalainen et al. 2024; Shiu et al. 2024).

To simulate a brain as a synapse network based on a connectome, one could first precompute and store the effective resistance matrix for every reconstructed neuron. Then simulating a synapse network would require O(*N*_out_ × *N*_in_) operations for every neuron due to the matrix-vector multiplication in Eq. (1), a dramatic increase in computational complexity relative to a traditional neural network, which requires O(*N*_out_ + *N*_in_) operations per point neuron. Replacing the effective resistance matrix in Eq. (1) by its hierarchical decomposition would reduce the computational complexity of simulation, due to the low rank and block structure. So the present work makes it possible to simulate synapse networks while incurring surprisingly little additional cost relative to neural networks.

The present work could be extended to dynamics by adding capacitors to the resistive grid, which results in a differential equation with the same sparse matrix of conductances. Numerical simulation is more challenging than finding the steady state solution, because the linear dynamical system is very stiff. Simply adding a τ*dV*_*i*_/*dt* term to the left hand side of Eq. (1) is more computationally tractable, and could be a reasonable approximation for some purposes even though it is mathematically incorrect.

## Methods

### Spiking, non-spiking, or hybrid

Given the lack of electrophysiological recordings in the literature, it is impossible to know at present whether Dm interneurons are spiking, non-spiking, or hybrid.

Transcriptomic information is available for some fly visual neuron types, including Dm types. Transcription of *para*, the gene that codes for voltage-gated sodium channels (Loughney, Kreber, and Ganetzky 1989; O’Dowd, Germeraad, and Aldrich 1989), might seem to be a predictor of spiking. But certain nonspiking local interneurons in the olfactory system are known to transcribe *para* without translating it (Schenk and Gaudry 2023). Therefore proteomic rather than transcriptomic information is needed.

One might attempt to predict whether a neuron is spiking on the basis of morphological criteria, such as the presence of an axon. Indeed, in some visual and olfactory neurons, it has been shown that Para protein is enriched in a “distal axonal segment” that is analogous to the axon initial segment of vertebrate neurons (Ravenscroft et al. 2020). However, it is already known that the presence of an axon does not reliably indicate whether a neuron is spiking. For example, the “broad” class of local interneuron in the antennal lobe lacks axons (Philipp Schlegel et al. 2021), yet most such cells are thought to be spiking (Chou et al. 2010; Seki et al. 2010). Conversely, certain types of columnar neurons in the optic lobe have axons, yet appear to be nonspiking based on electrophysiological recordings (Mu et al. 2012; Behnia et al. 2014).

### Passive electrical model

The effective resistance matrix *R* is the inverse of the conductance matrix, which is the sum of a graph Laplacian and a nonnegative diagonal matrix. The effective resistance is nonnegative and symmetric, as required by physical law. To ward off confusion, it should be noted that the term “effective resistance” in graph theory often denotes a related quantity, *d*_*ij*_ = 2*R*_*ij*_ − *R*_*ii*_ − *R*_*jj*_, which satisfies the triangle inequality and the other properties of a distance metric (Spielman 2019).

The 3D reconstruction of a neuron is converted into a 3D resistive grid as follows. Each voxel corresponds to a node in the grid. For each pair of nearest neighbor voxels, an intracellular conductance *l*_*x*_ /ρ_*i*_ or *l*_*y*_ /ρ_*i*_ or *l*_*z*_ /ρ_*i*_ is placed between the corresponding nodes, depending on whether the voxels are separated along the *x, y*, or *z* axis. The 6-neighborhood is used, and voxel dimensions are (*l*_*x*_, *l*_*y*_, *l*_*z*_) = (32, 32, 40) nm after downsampling. For any face of a voxel with no neighboring voxel in the neuron, a transmembrane conductance *l*_*y*_ *l*_*z*_ /ρ_*m*_ or *l*_*x*_ *l*_*z*_ /ρ_*m*_ or *l*_*x*_ *l*_*y*_ /ρ_*m*_ is placed from the corresponding node to ground. Applying Kirchoff’s and Ohm’s Laws yields

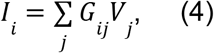

where the conductance matrix *G* is the sum of a Laplacian matrix and a diagonal matrix. The Laplacian matrix contains the intracellular conductances in a sparse set of negative off-diagonal elements, and the positive diagonal elements are such that row and column sums vanish. The diagonal matrix contains the transmembrane conductances. The conductance matrix is a symmetric, diagonally-dominant M-matrix or SDDM matrix.

In principle, Eq. (4) can be solved by inverting the conductance matrix, but this would be computationally intractable for the systems (2 to 38 million nodes) simulated here. Furthermore, the inverse of the conductance matrix is full rather than sparse, so merely storing the result could be difficult. Instead, Eq. (4) is solved only for current injection into the nodes corresponding to input synapses, and only the voltages of the nodes corresponding to output synapses are stored. Repeated solution of Eq. (4) for multiple current injections is efficiently done after Cholesky decomposition of the conductance matrix, which is standard in many numerical linear algebra software packages. The larger Dm cells were solved on a computer with 256 GB of RAM.

### Resistivity parameters

The resistivities *ϱ*_*i*_ = 250 Ω cm and *ϱ*_m_ = 20000 Ω cm^2^ are round numbers based on electrophysiological recordings of “Cell 3” in a fly olfactory system (Gouwens and Wilson 2009). The resistivities enter the model only through their ratio 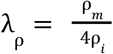, which is *λ*_ρ_ = 20 cm for these Gouwens-Wilson parameters.

Meier and Borst estimated *ϱ*_*i*_ = 400 Ω cm and *ϱ*_m_ = 8000 Ω cm^2^ from calcium imaging of the fly visual neuron CT1 (Meier and Borst 2019). Their parameters are also consistent with fits of passive models to voltage imaging of the fly visual neuron Mi1 (Yang et al. 2016), and lead to a ratio *λ* _ρ_ = 5 cm.

The ratio *λ* _ρ_ influences the length constant of the decay through 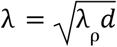, which is the geometric mean of the neurite caliber (diameter) *d* and *λ*_ρ_. Changes in resistivity and dendrite caliber should influence the length constant rather weakly, due to the presence of the square root.

### Neurotransmitter and receptor identity

Both the Janelia optic lobe reconstruction and FlyWire provide predictions of neurotransmitter identity that are based on the same computer vision methods applied to different EM datasets (Eckstein et al. 2024). The Janelia reconstruction predicts that Dm15, Dm6, and Dm17 are all glutamatergic (Aljoscha Nern et al. 2024). FlyWire predicts that Dm15 is glutamatergic, Dm6 is GABAergic, and provides no prediction for Dm17. Both Janelia and FlyWire predict that L2 is cholinergic.

L2 is predicted cholinergic by published transcriptomic surveys, which do not cover the three Dm types (Davis et al. 2020; Özel et al. 2021; Yoo et al. 2023).

Acetylcholine is excitatory when the postsynaptic receptor is nicotinic, which is generally the case in the fly brain (Rosenthal and Yuan 2021). GABA is generally inhibitory. Glutamate is often inhibitory, but can sometimes be excitatory depending on the identity of the postsynaptic receptor. Glutamate is inhibitory when the postsynaptic receptor is GluClα (W. W. Liu and Wilson 2013). According to a molecular study, L2 expresses the GluClα receptor. Therefore Dm15, Dm6, and Dm17 are highly likely to inhibit L2.

The interpretation of the Dm cells as playing a role in spatial normalization of L2 activity depends on the assumption that they are inhibitory (Seung 2024a). The validity of the Dm simulations in the present work, however, does not depend on the assumption that Dm cells are inhibitory. The validity of modeling synapses as current sources does depend on L2 being excitatory, because this approximation is better when the reversal potential of the synapse is far from the resting potential.

### Hierarchical decomposition

The hierarchical decompositions of Dm6 and Dm17 were computed as follows. I computed the effective resistance matrix between all synapse locations, whether they were input or output synapses. The resulting matrix was (*N*_out_ + *N*_in_)×(*N*_out_ + *N*_in_), and was preferred because its symmetry guaranteed a simplification, that the same spatial domains defining block structures would apply to both input and output synapses. The following semiautomated procedure was used to decompose the matrix.

Distances between synapse locations were computed by the Pearson correlation or Jaccard distance between corresponding rows/columns of the matrix. Hierarchical clustering with single or complete linkage was used to split the synapse locations into two groups. Then SVD was used to factorize the rectangular matrix from one group of locations to the other group of locations. If the second singular value was tiny and the groups were roughly balanced, then this split was accepted. If not, then hierarchical clustering was used to split the synapse locations into more than two clusters. These clusters were manually merged in various ways in order to find two roughly balanced groups that passed the SVD test. The same procedure was recursively applied to divide each of the two groups into two subgroups, and so on.

This procedure led to the hierarchical decompositions shown in Fig. 6, which are almost exact. The difference between the original matrix and the decomposition is less than 10^-6^ for all elements. After the square symmetric matrix was decomposed, the submatrix corresponding to the input and output synapse locations was retained.

## Acknowledgements

I thank the FlyWire Consortium for the connectome and cell types on which this analysis is based. I am grateful for advice from Michael Hausser on dendritic biophysics, Dan Spielman on Laplacian systems, Misha Tsodyks on network theory, Yerbol Kurmangaliyev on fly transcriptomics, and Damon Clark, Tim Currier, and James Jeanne on fly neurophysiology.

## Extended Data Figures

**Extended Data Figure 1.**
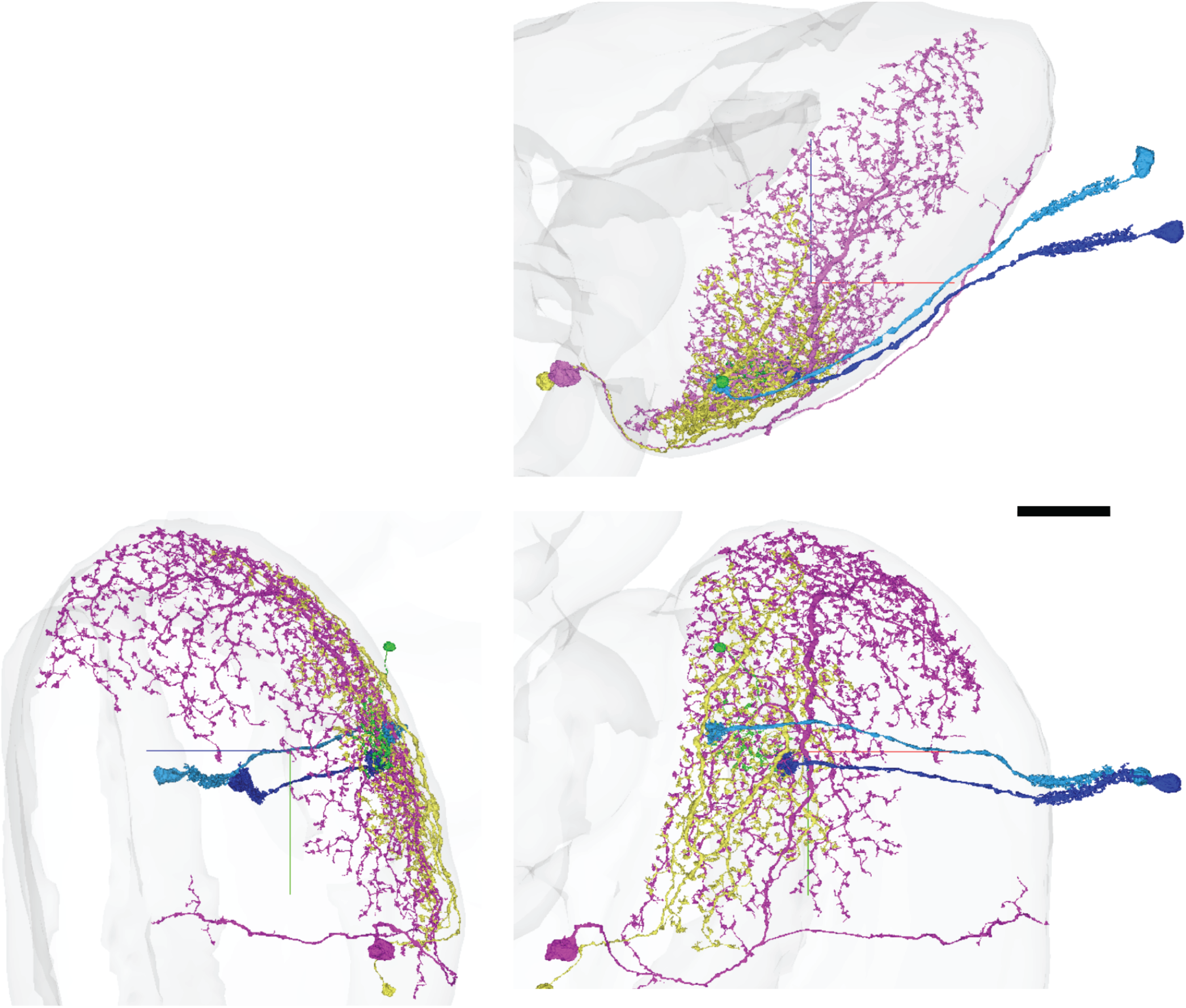
Orthogonal views of Dm and L2 cells. Dm15 (green), Dm6 (yellow, and Dm17 (purple) cells are each reciprocally connected with L2 cells (cyan, blue).

**Extended Data Figure 2.**
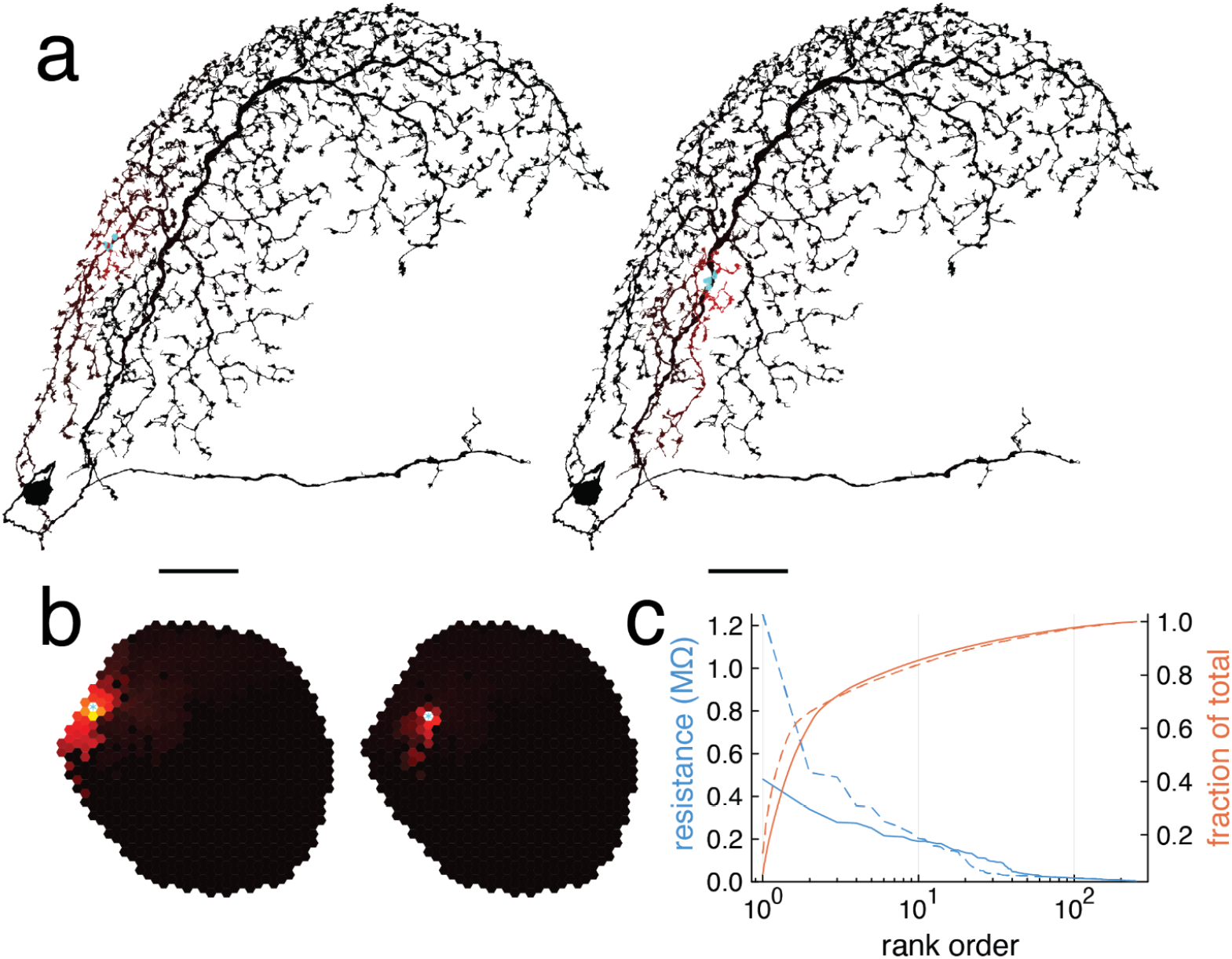
Simulation of a large Dm cell. Simulations of the same Dm17 and L2 cells as in Fig 4 but with Meier-Borst parameters. **a**, Simulated Dm17 voltage (colormap) caused by activating input synapses (cyan) from L2 cells (Fig. 1g). A voltage response is visible over only a small fraction of the arbor. **b**, Dm17 voltages of (**a**) mapped to visual locations. The color of each hexel represents the mean voltage at all the output synapses onto the L2 cell located at that hexel. The response to the left L2 input looks broader than the response to the right L2 input. **c**, Dm17 voltages of (**b**) after rank ordering. Attenuation is stronger for the right L2 input (dashed) than for the left L2 input (solid).

## Supplementary Data

### Supplementary Data 1

All columns of the effective resistance matrix of the Dm15 cell for Gouwens-Wilson parameters (Fig. 2, left). Cyan star indicates the location of the stimulated L2-Dm15 synapses. The colormap maximum is set to the maximum value for each heatmap.

### Supplementary Data 2

All columns of the effective resistance matrix of the Dm15 cell for Meier-Borst parameters (Fig. 2, right).

### Supplementary Data 3

All columns of the effective resistance matrix of the Dm6 cell for Gouwens-Wilson parameters (Fig. 3, left).

### Supplementary Data 4

All columns of the effective resistance matrix of the Dm6 cell for Meier-Borst parameters (Fig. 3, right).

### Supplementary Data 5

All columns of the effective resistance matrix of the Dm17 cell for Gouwens-Wilson parameters (Fig. 4).

### Supplementary Data 6

All columns of the effective resistance matrix of the Dm17 cell for Meier-Borst parameters (Extended Data Fig. 2).

### Supplementary Data 7

Residual effective resistance matrix of the Dm6 cell remaining after hierarchical decomposition for Meier-Borst parameters (Fig. 6, top).

### Supplementary Data 8

Residual effective resistance matrix of the Dm17 cell remaining after hierarchical decomposition for Gouwens-Wilson parameters (Fig. 6, bottom).

## Neuroglancer links

Dm15 720575940624368722

Dm6 720575940627980968

Dm17 720575940639823413

L2 720575940626299710

L2 720575940617175710

Fig. 1a, b. Input and output synapses of Dm6

https://ngl.cave-explorer.org/#!middleauth+https://global.daf-apis.com/nglstate/api/v1/4991061341503488

Fig. 1e, f, g. Extended Data Fig. 1. Dm15, Dm6, and Dm17 frontal view

https://ngl.cave-explorer.org/#!middleauth+https://global.daf-apis.com/nglstate/api/v1/5459592210284544

Extended Data Fig. 1. back view

https://ngl.cave-explorer.org/#!middleauth+https://global.daf-apis.com/nglstate/api/v1/5318854721929216

Extended Data Fig. 1. lateral view

https://ngl.cave-explorer.org/#!middleauth+https://global.daf-apis.com/nglstate/api/v1/5262322349113344

## Mathbox. Synapse networks

Sebastian Seung February 2, 2025

Eq. (1) was introduced to model electrical signaling within a single neuron. It can be generalized so that the indices *i* and *j* run over all synapses in a neural circuit or entire brain. This multi-cell effective resistance matrix is sparse, as *R*_*ij*_ vanishes if there is no intracellular path by which the presynaptic voltage *V*_*i*_ can be affected by the postsynaptic current *I*_*j*_.

### Network of synapses

For a closed set of equations, there should also be a model of synaptic transmission between neurons. The postsynaptic current generated by synapse *j* is assumed to be a function of the presynaptic voltage of synapse *j*,

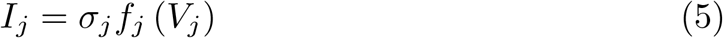

The nonlinear activation function *f*_*j*_ is assumed nonnegative. The parameter *σ*_*j*_ is +1 for an excitatory synapse, and −1 for an inhibitory synapse. Combining Eqs. (1) and (5) yields a steady state equation for a synapse network,

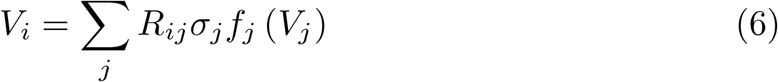

Neurons are implicit in the above equations. They can be made explicit by writing the effective resistance matrix as 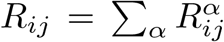, where the sum runs over all neurons *α*. The elements of 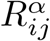 are positive only if synapses *i* and *j* are output and input synapses, respectively, of a particular cell *α*. Neuron *α* receives a connection from neuron *β* if there exists some *i, j*, and *k* such that 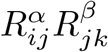 is nonzero, or equivalently if synapse *j* is an input synapse of neuron *α* and an output synapse of neuron *β*. Therefore neural connectivity is implicit in the effective resistance matrix *R*_*ij*_.

Let 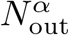 and 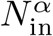 denote the number of output and input synapses, respectively, of neuron *α*. The total number of synapses in the network is 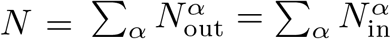. The latter two sums are equal assuming that every synapse has one input and one output cell. As defined above, the matrices *R*^*α*^ are *N ×N*, like *R*. However, *R*^*α*^ can also be regarded as an 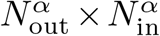 matrix by restricting its indices to its output and input synapses; all other elements are vanishing. It is this 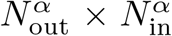 matrix that is defined in Eq. (1) and computed by the resistive grid simulations in the main text.

### Point neuron approximation

In the point neuron approximation, *R*_*ij*_ ≈ Σ_*α*_*R*^*α*^ *c*_*iα*_*d*_*αj*_ Here *c*_*iα*_ = 1 if synapse *i* is an output of cell *α*, and vanishes otherwise. Similarly, *d*_*αj*_ = 1 if synapse *j* is an input to cell *α*, and vanishes otherwise. These are both assignment matrices, meaning that every synapse is assigned to one presynaptic cell and one postsynaptic cell. Then Eq. (6) simplifies to *V*_*i*_ = Σ_*α*_ *c*_*iα*_*V*_*α*_ where

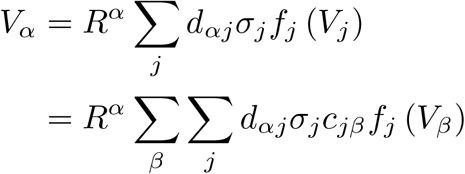

is the shared voltage of all output synapses of cell *α*. The second line follows from the identity *f*_*j*_ (Σ_*β*_ *c*_*jβ*_*V*_*β*_) = Σ_*β*_ *c*_*jβ*_*f*_*j*_ (*V*_*β*_), which is valid since *c*_*jβ*_ is an assignment matrix. If all *α* ← β synapses share the same *σ* and activation function, then

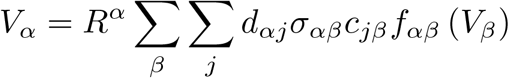

This can be written in the same form as the model of Lappalainen et al. (2024),

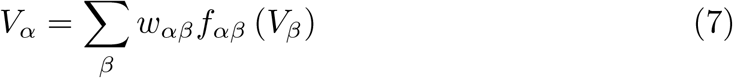

where *w*_*αβ*_ = *R*_*α*_*σ*_*αβ*_ Σ_*j*_ *d*_*αj*_ *c*_*jβ*_ = *R*_*α*_*σ*_*αβ*_*n*_*αβ*_ and *n*_*αβ*_ is the number of *α* ← *β* synapses.

### Virtual neurons

More generally, suppose that the effective resistance is written as

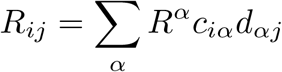

where *c* and *d* are not constrained to be assignment matrices. Here *α* is allowed to be a virtual neuron, in which case *c*_*iα*_ and *d*_*αj*_ can be arbitrary vectors.

Then Eq. (6) becomes

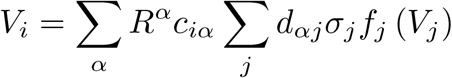

This can be rewritten as

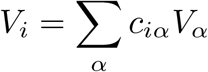

with the definition

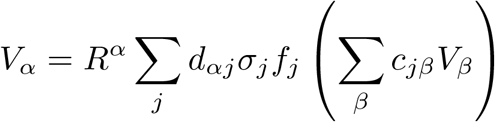

Then *V*_*α*_ is the voltage of (real or virtual) neuron *α*, with incoming connections *d*_*αj*_ and outgoing connections *c*_*iα*_.

