## Supplementary figures and images for "Multi-output computation by single neuron biophysics in a visual system"

### Data S1

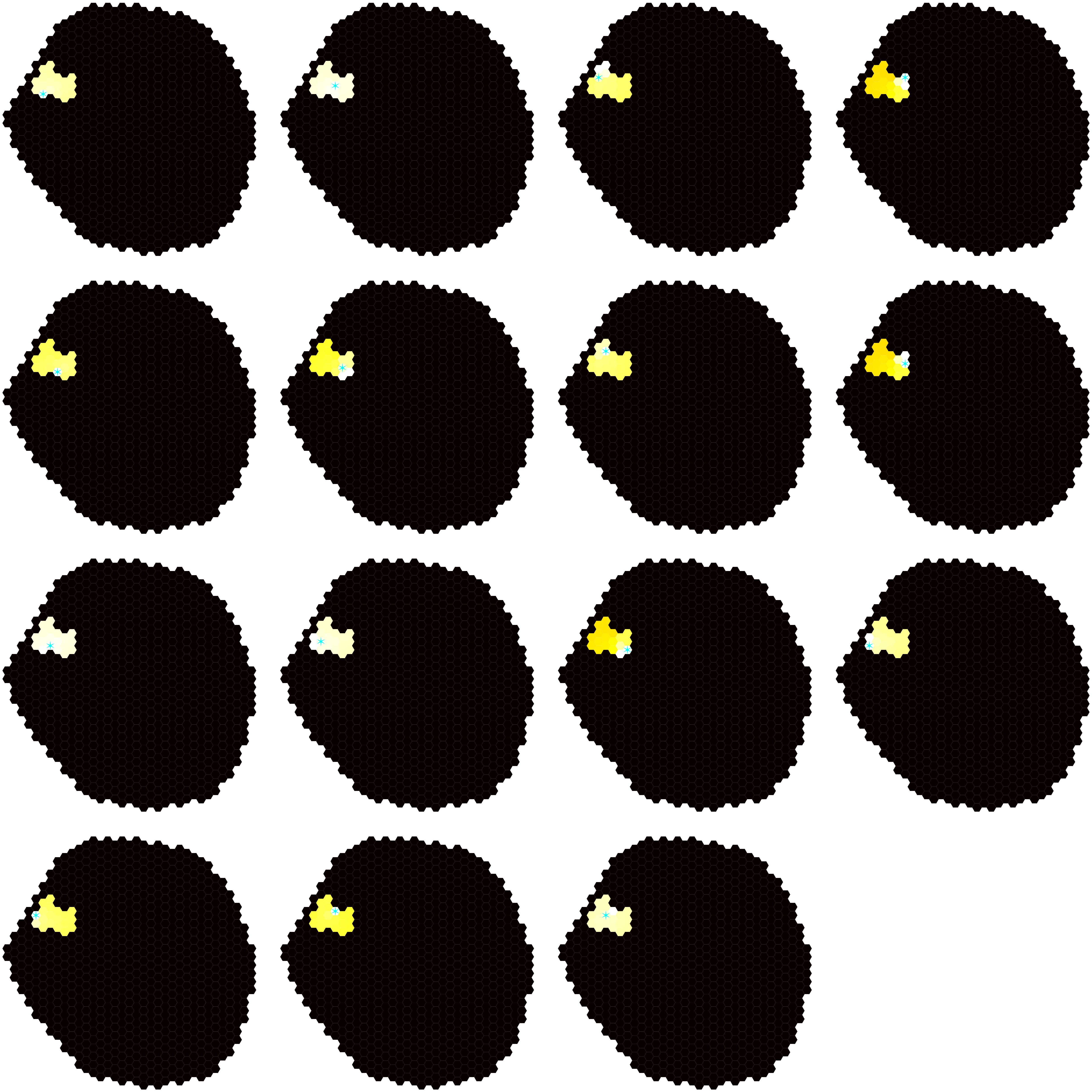

### Data S2

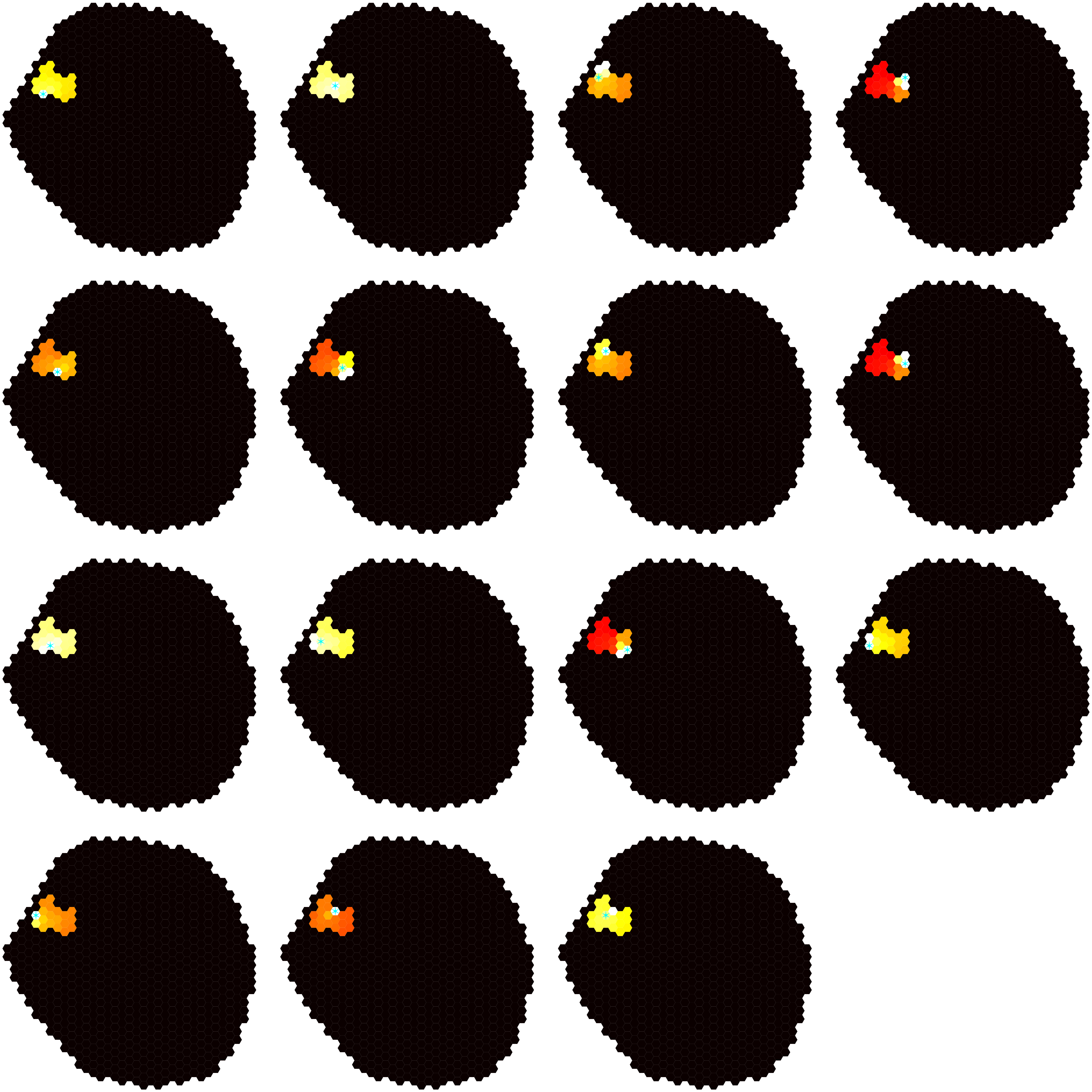

### Data S3

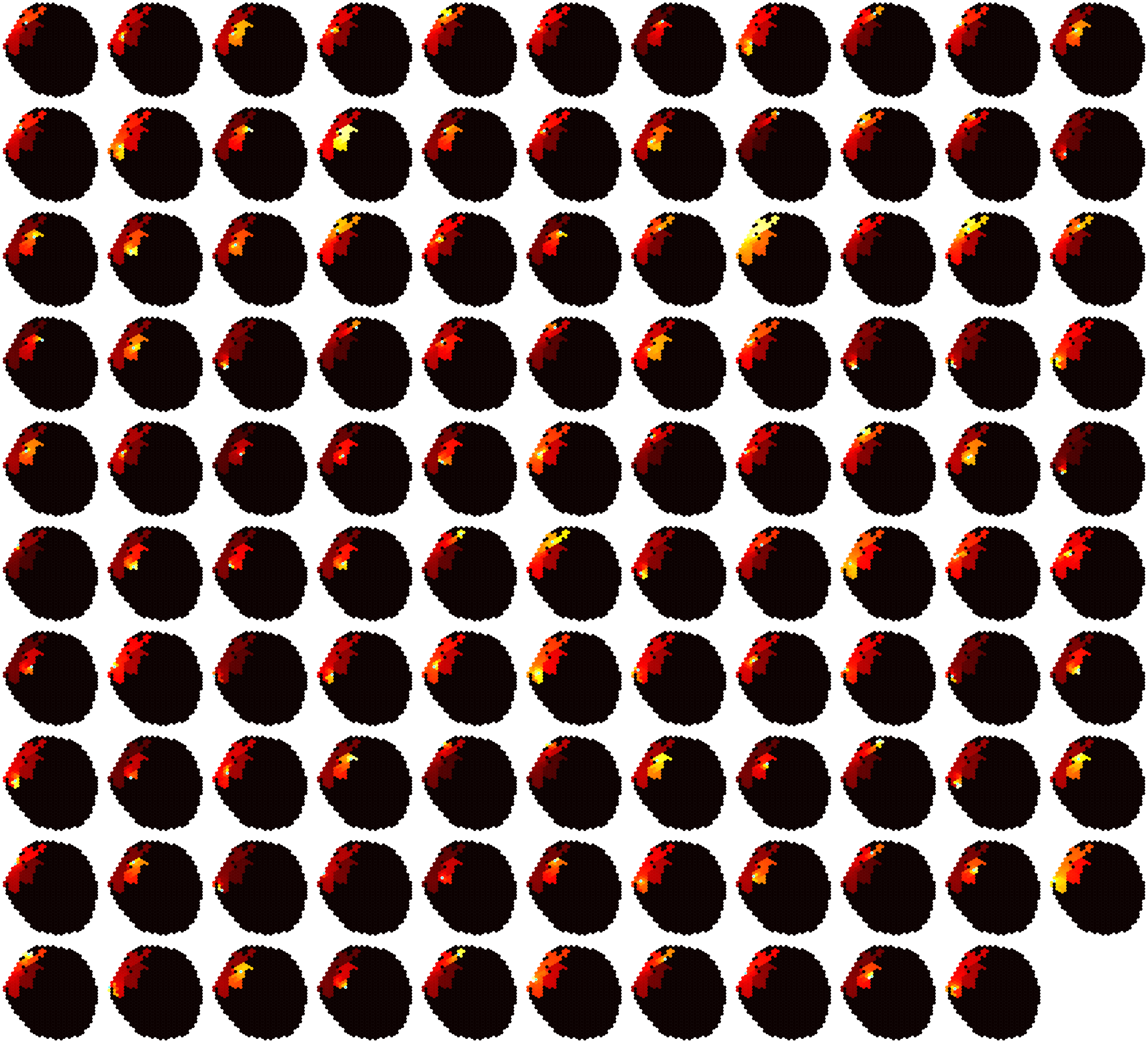

### Data S4

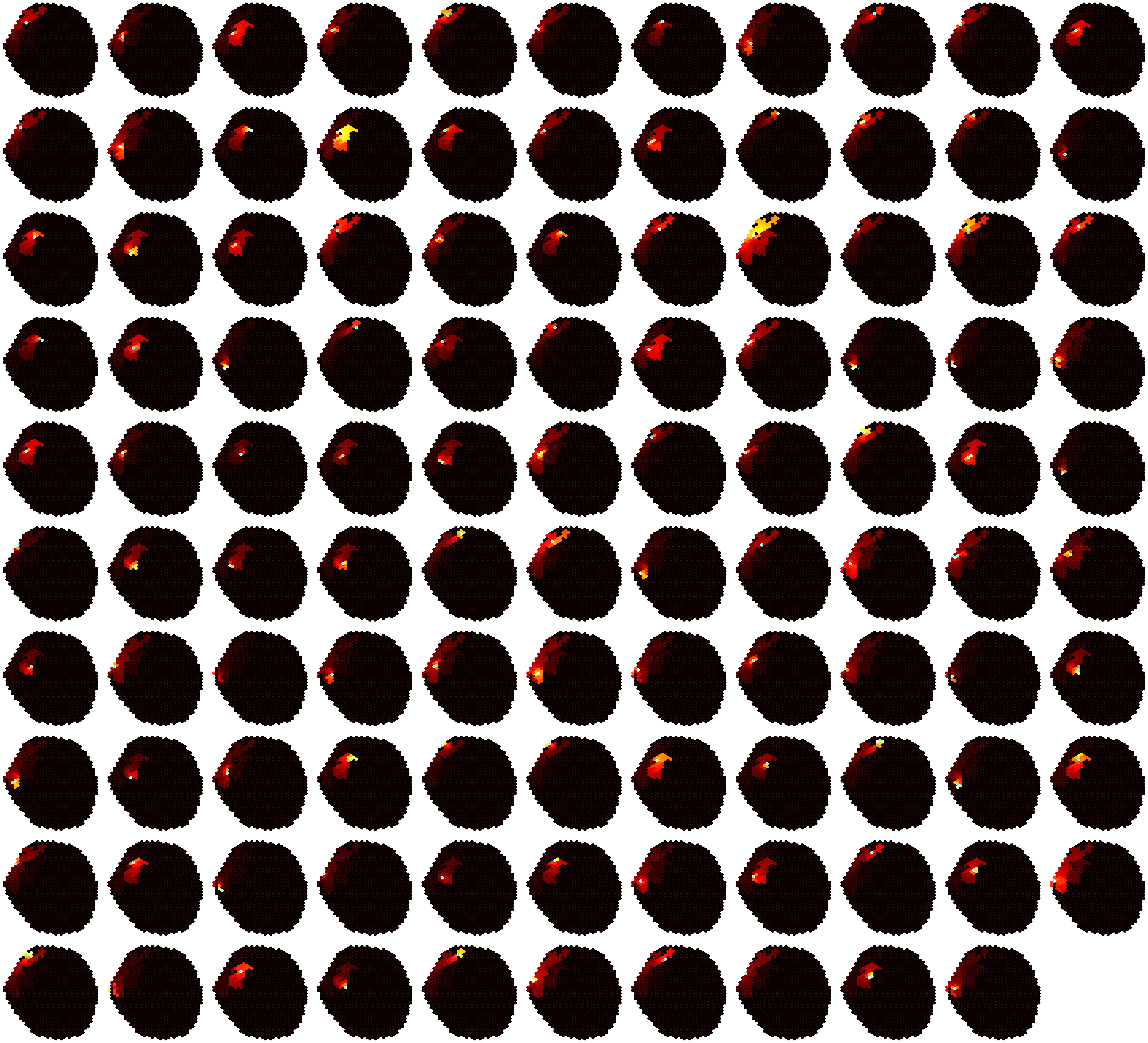

### Data S5

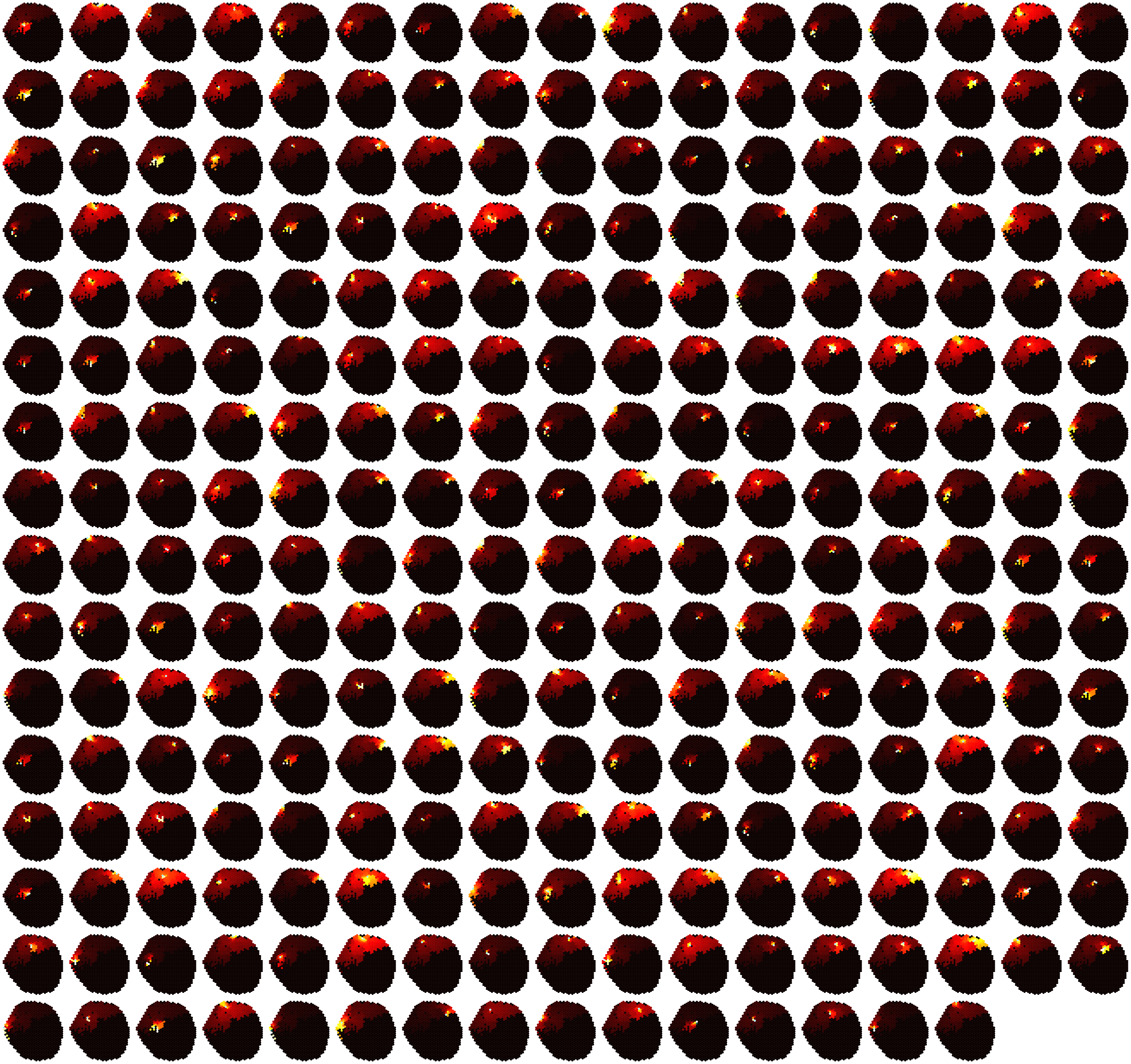

### Data S6

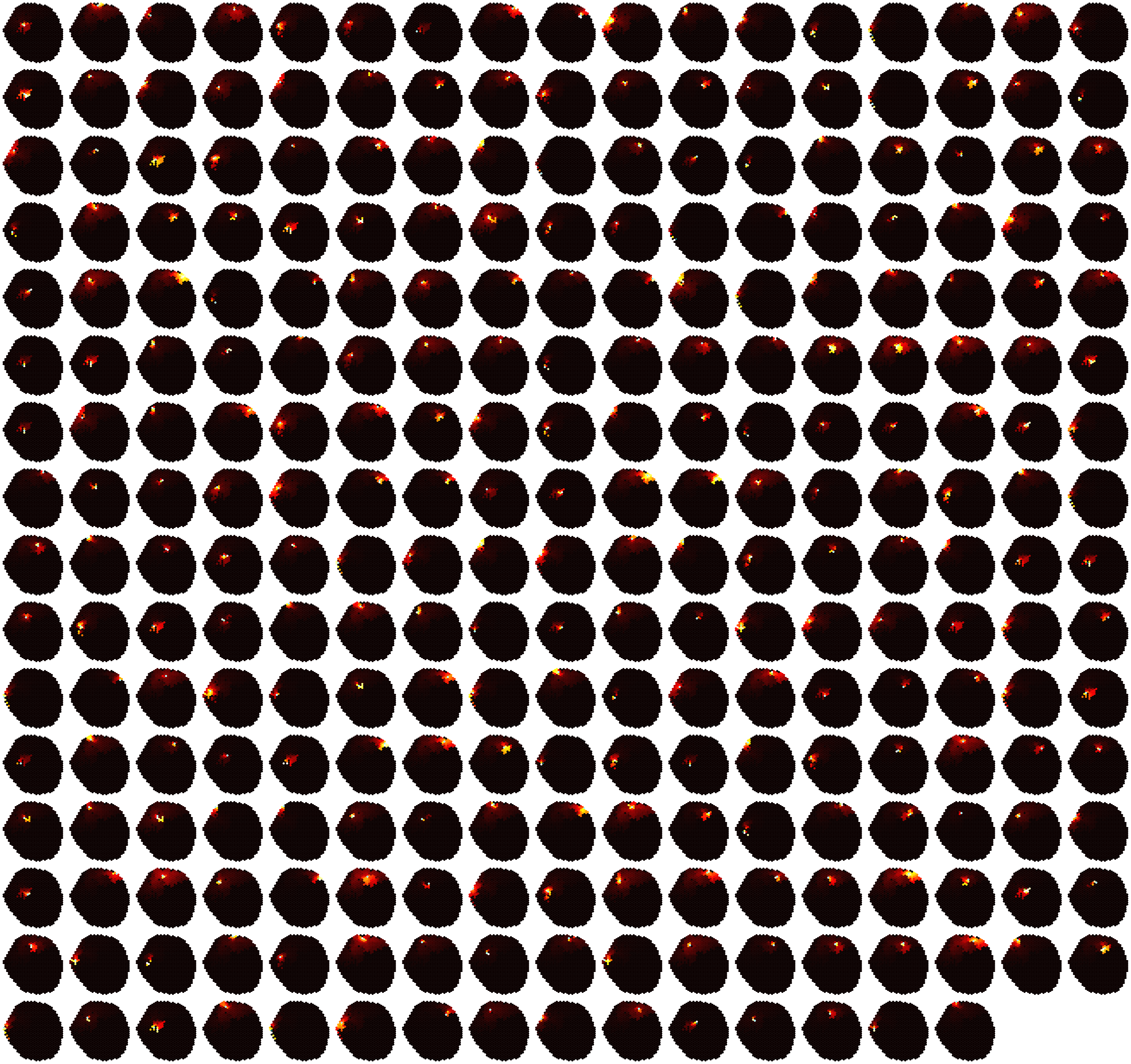

### Dm6-level5-1.pdf

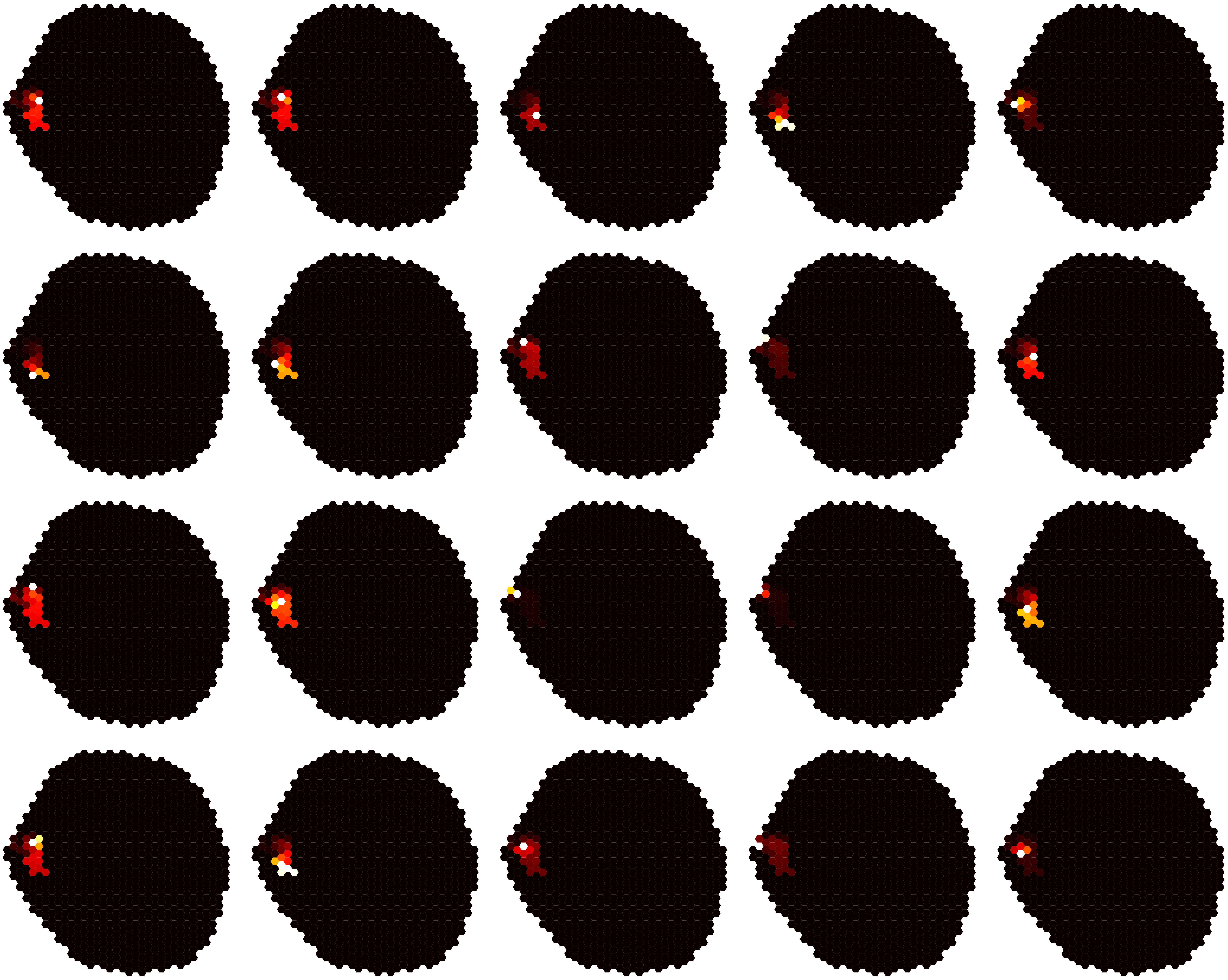

### Dm6-level5-2.pdf

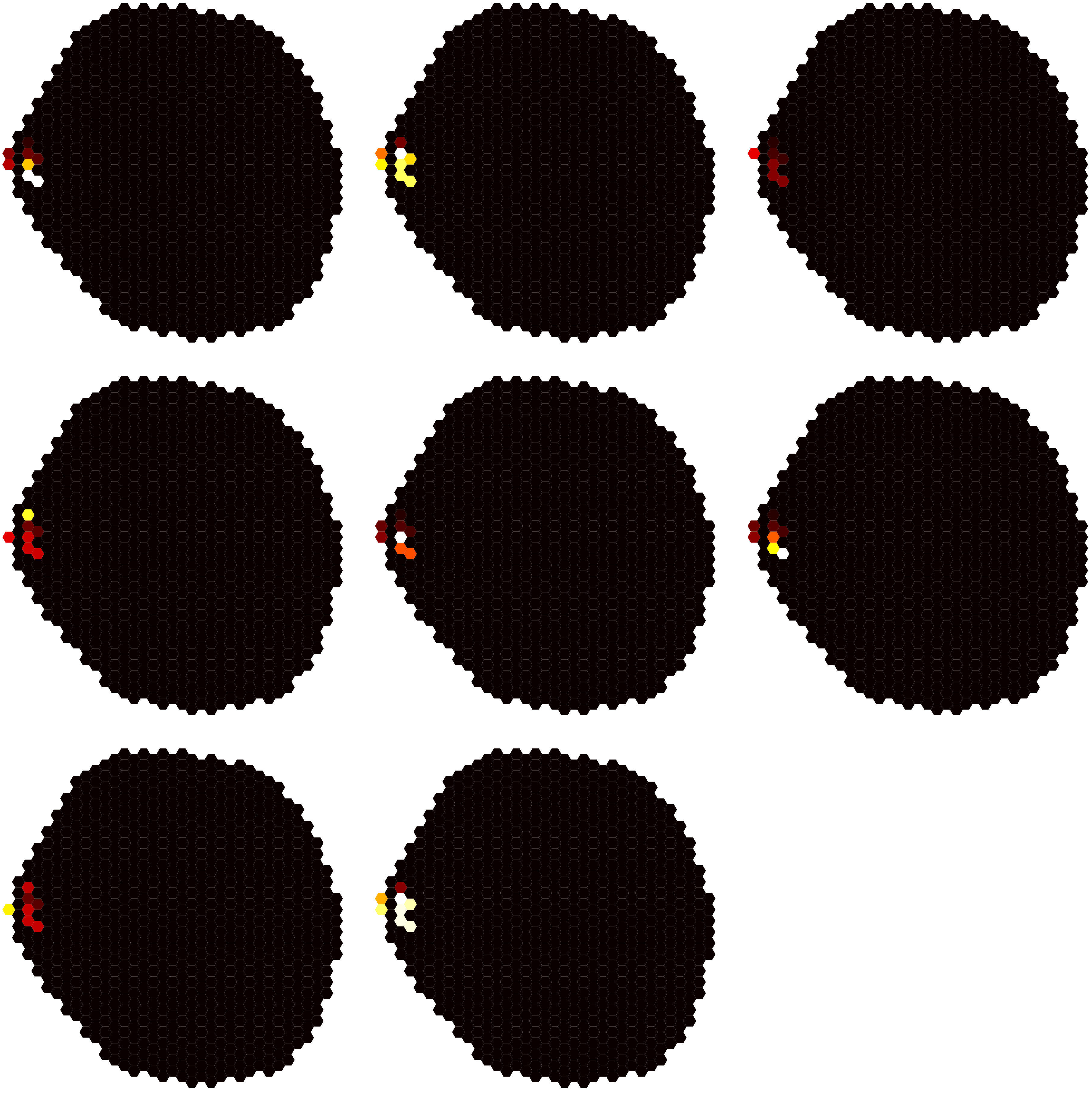

### Dm6-level5-3.pdf

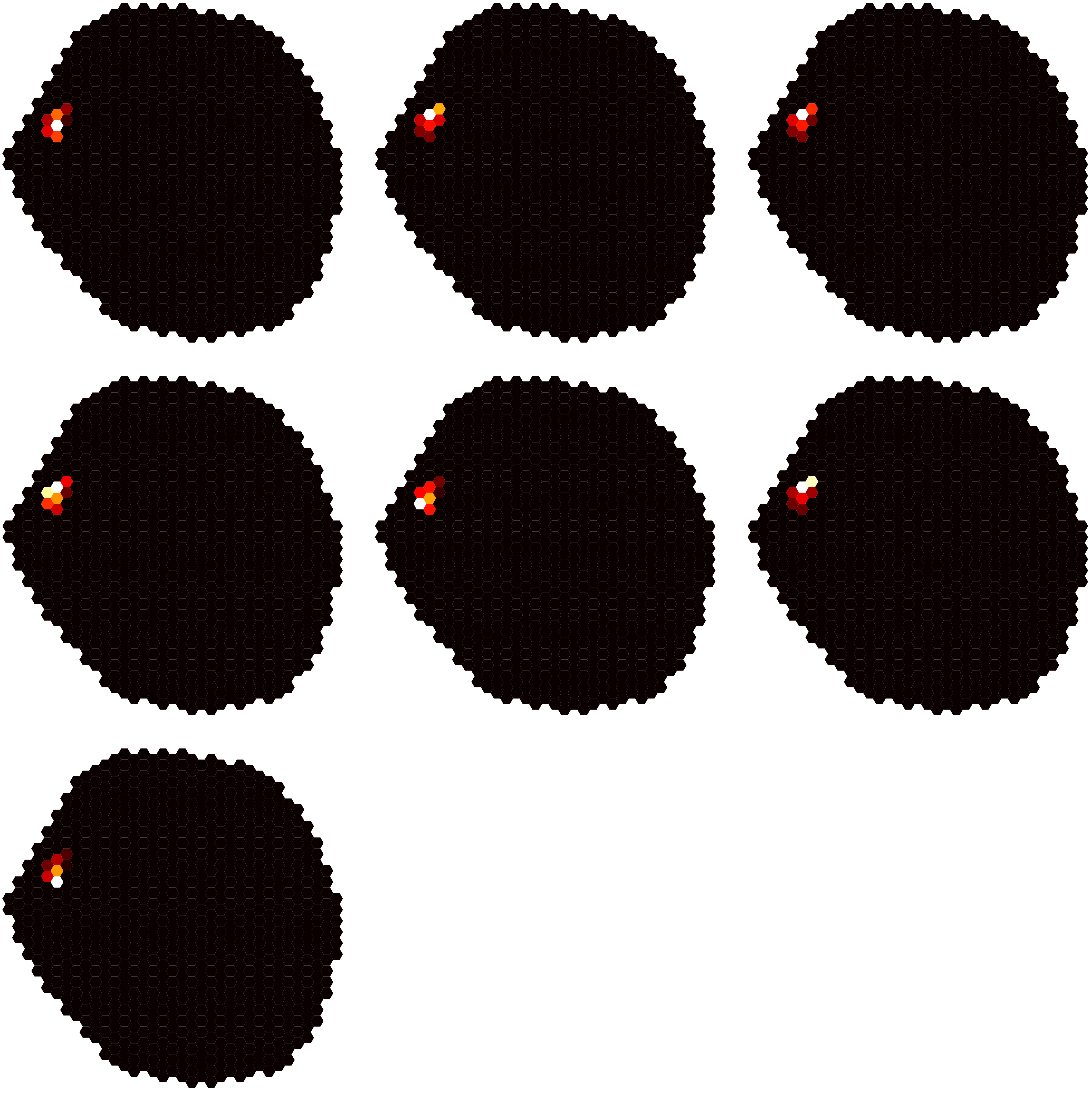

### Dm6-level5-4.pdf

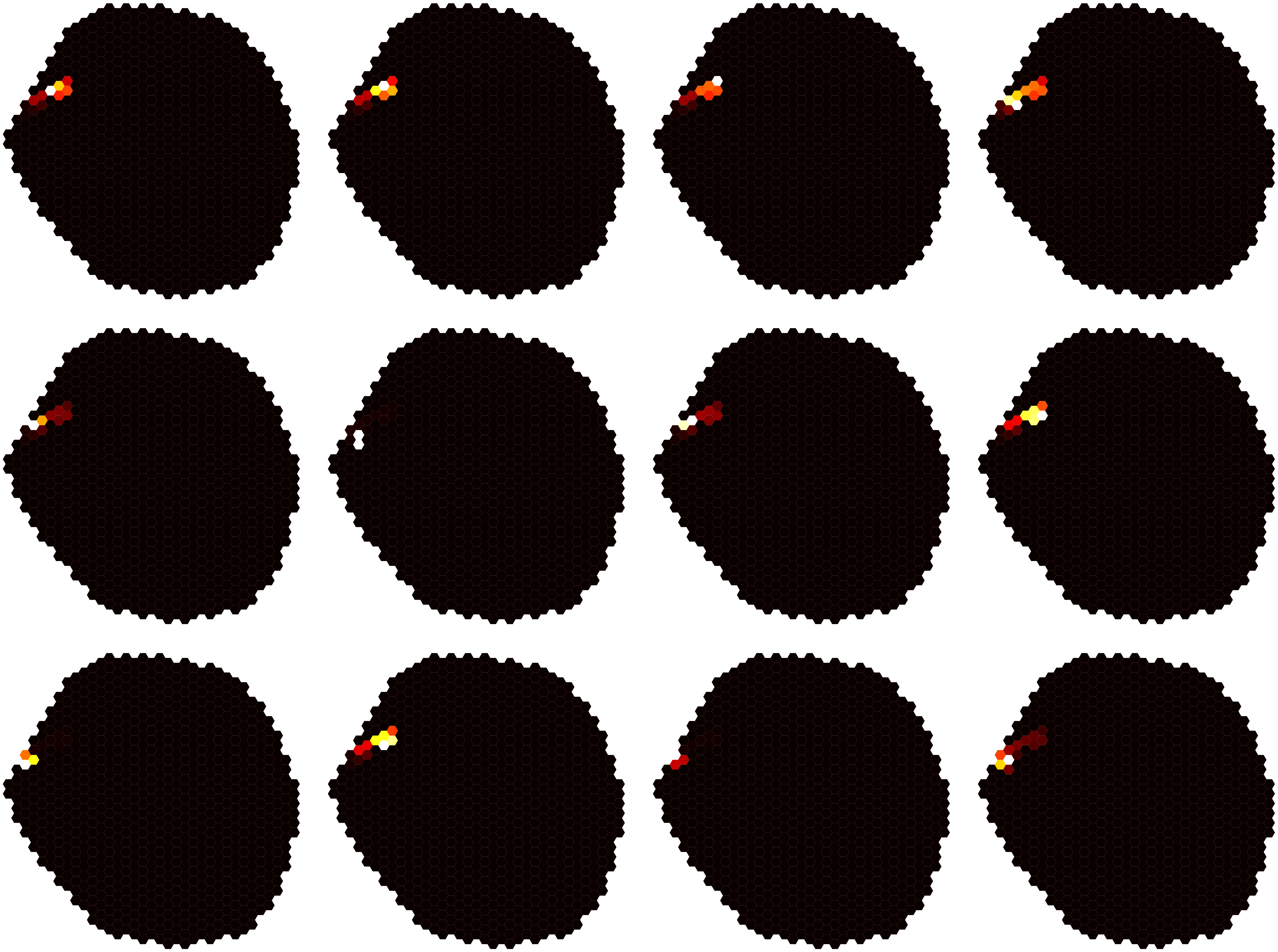

### Dm6-level5-5.pdf

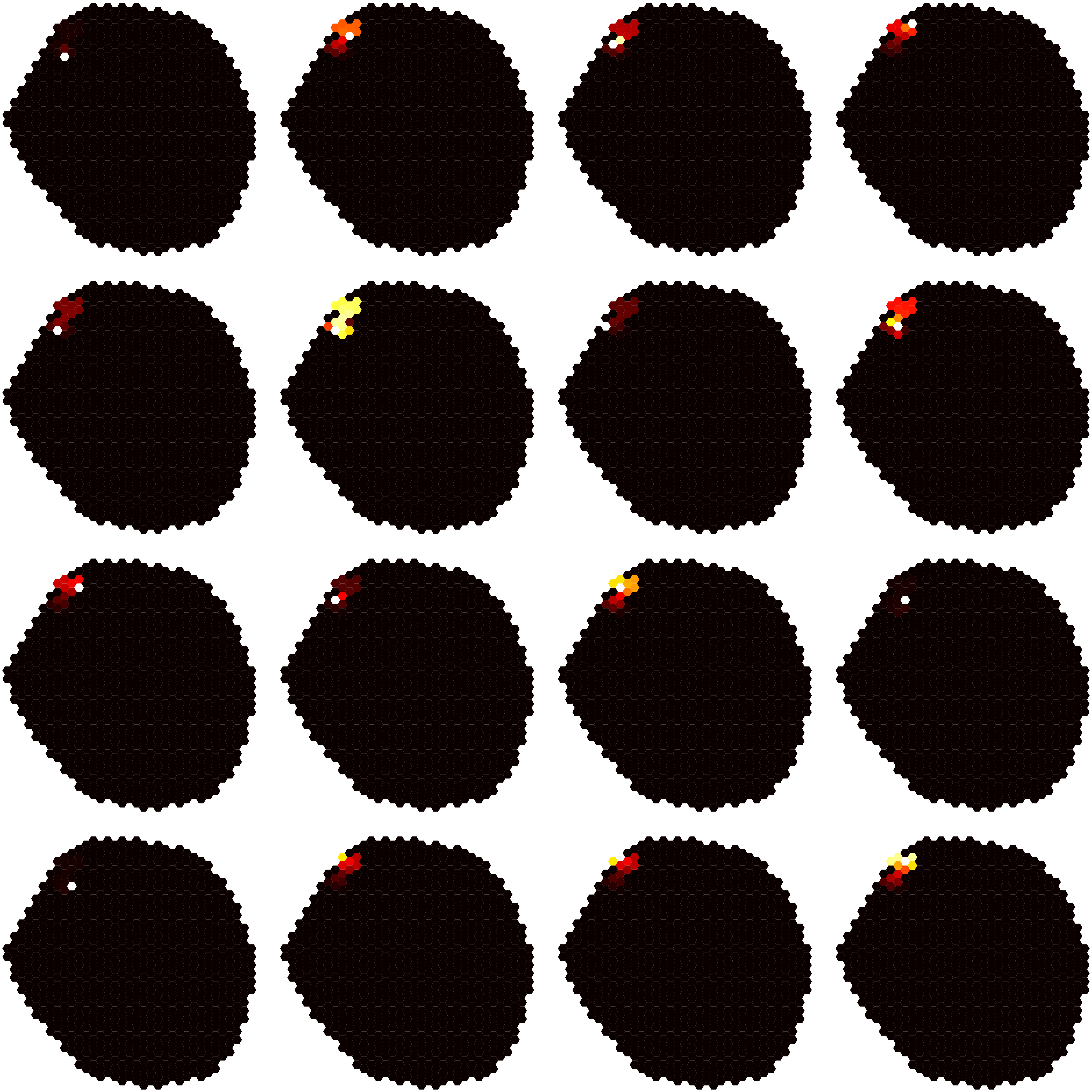

### Dm6-level5-6.pdf

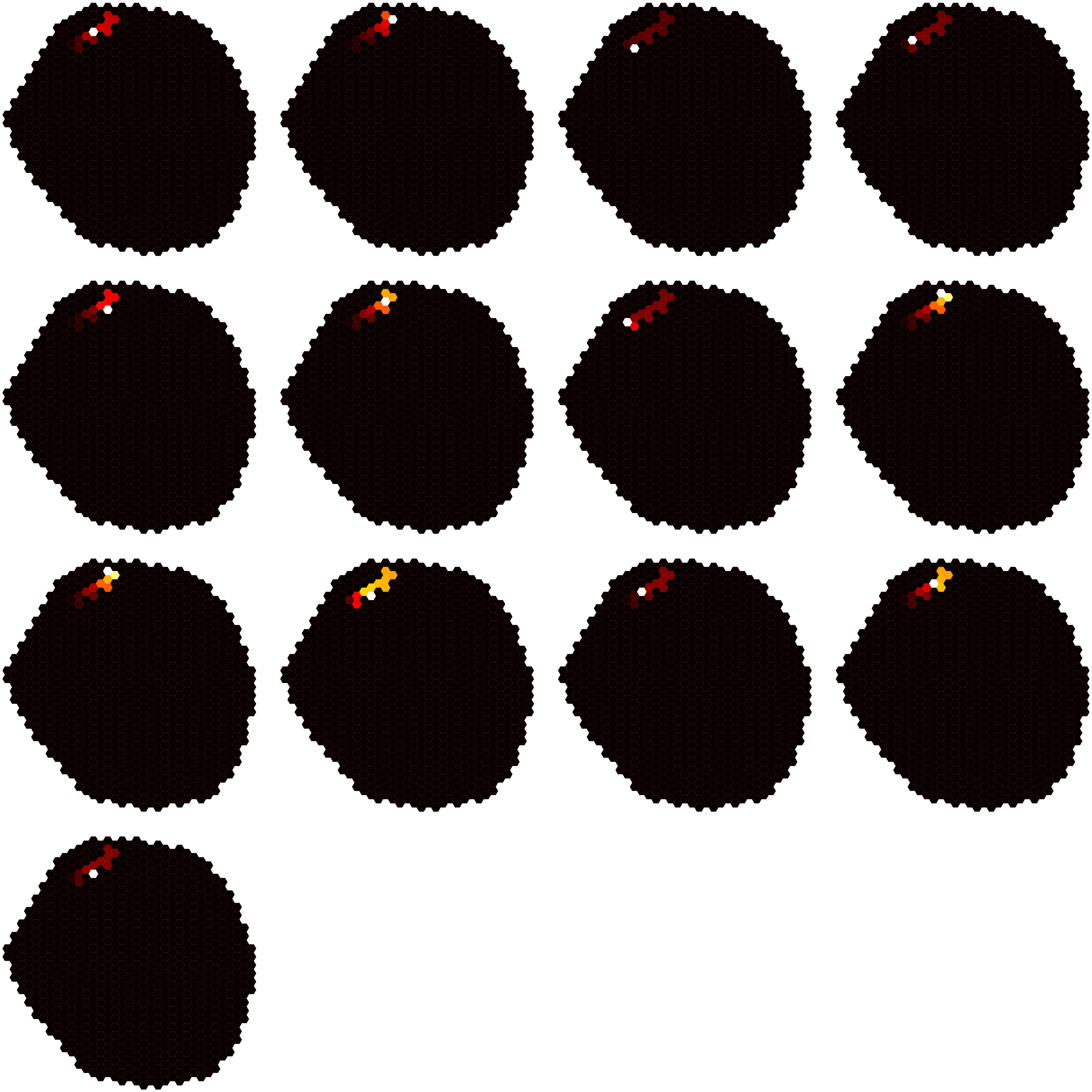

### Dm6-level5-7.pdf

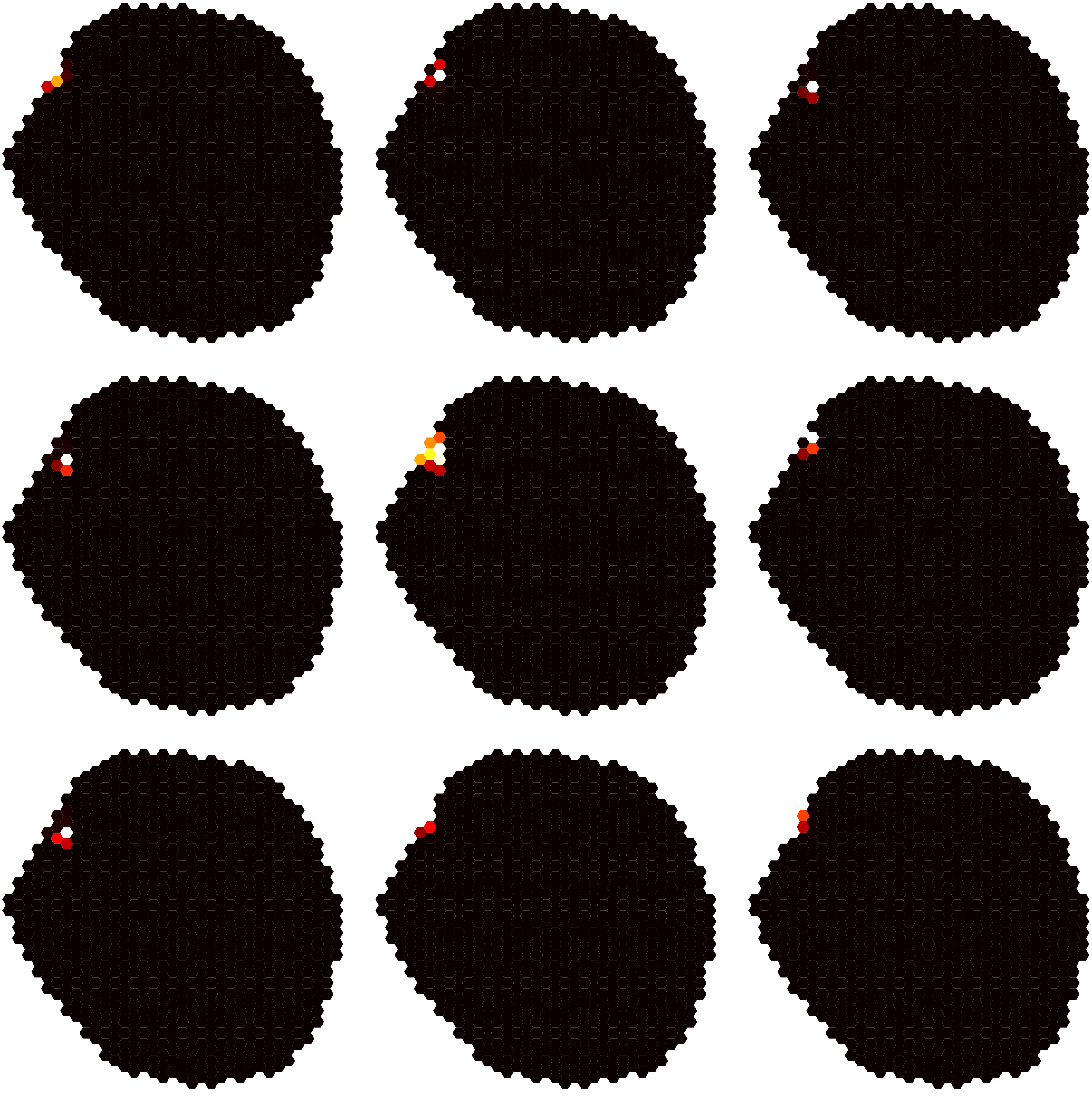

### Dm6-level5-8.pdf

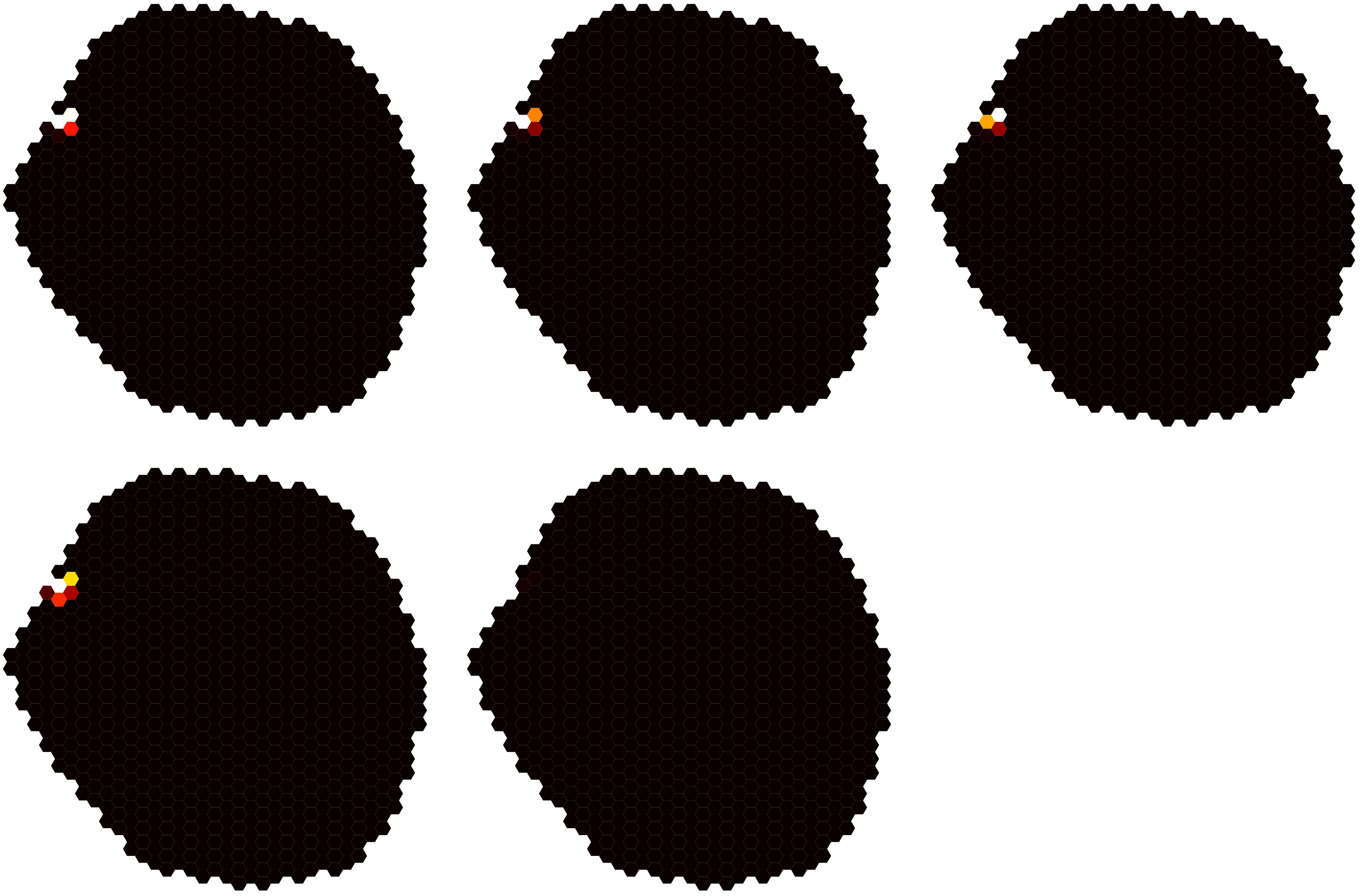

### Dm6-level5-9.pdf

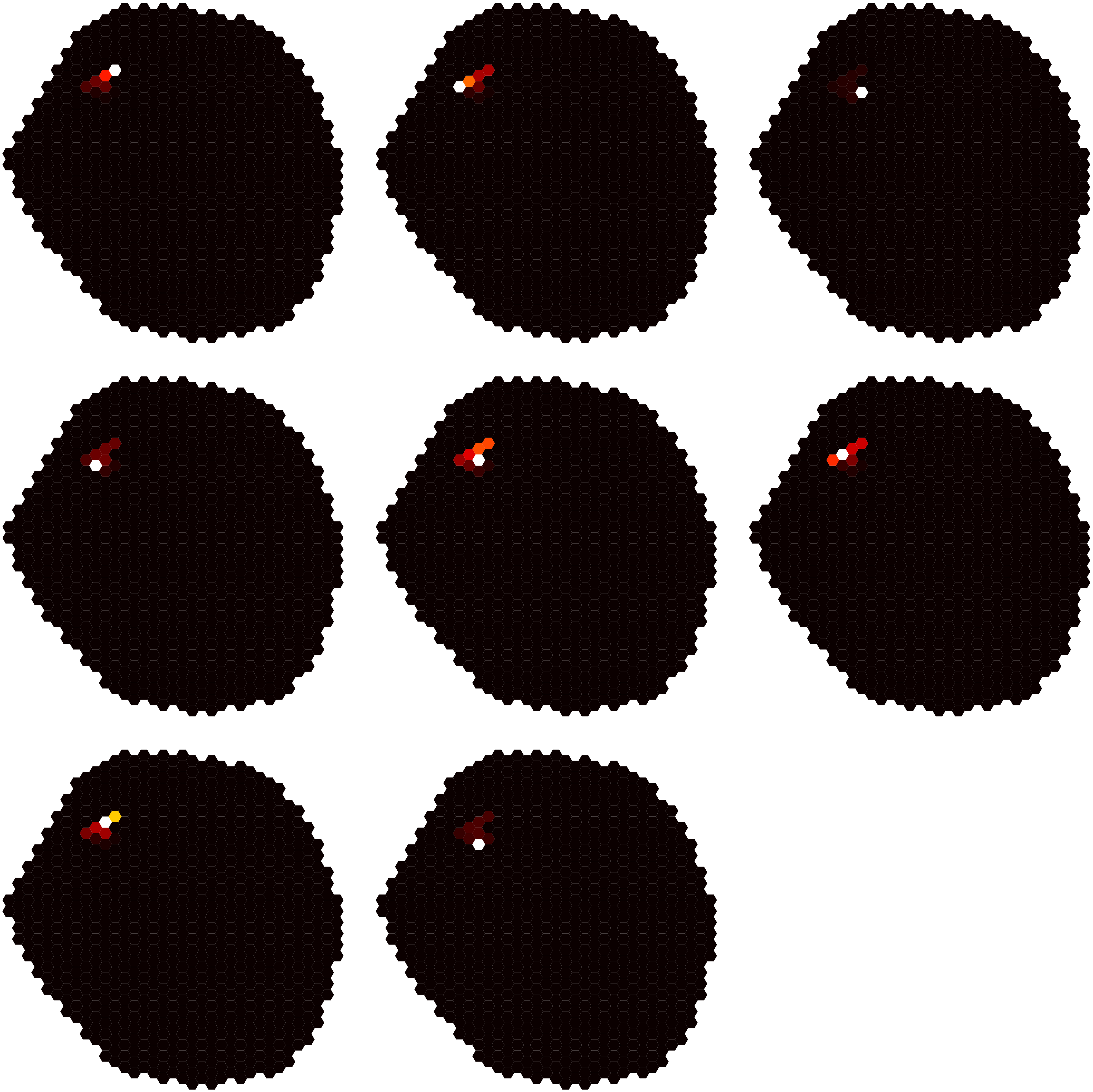

### Dm6-level5-10.pdf

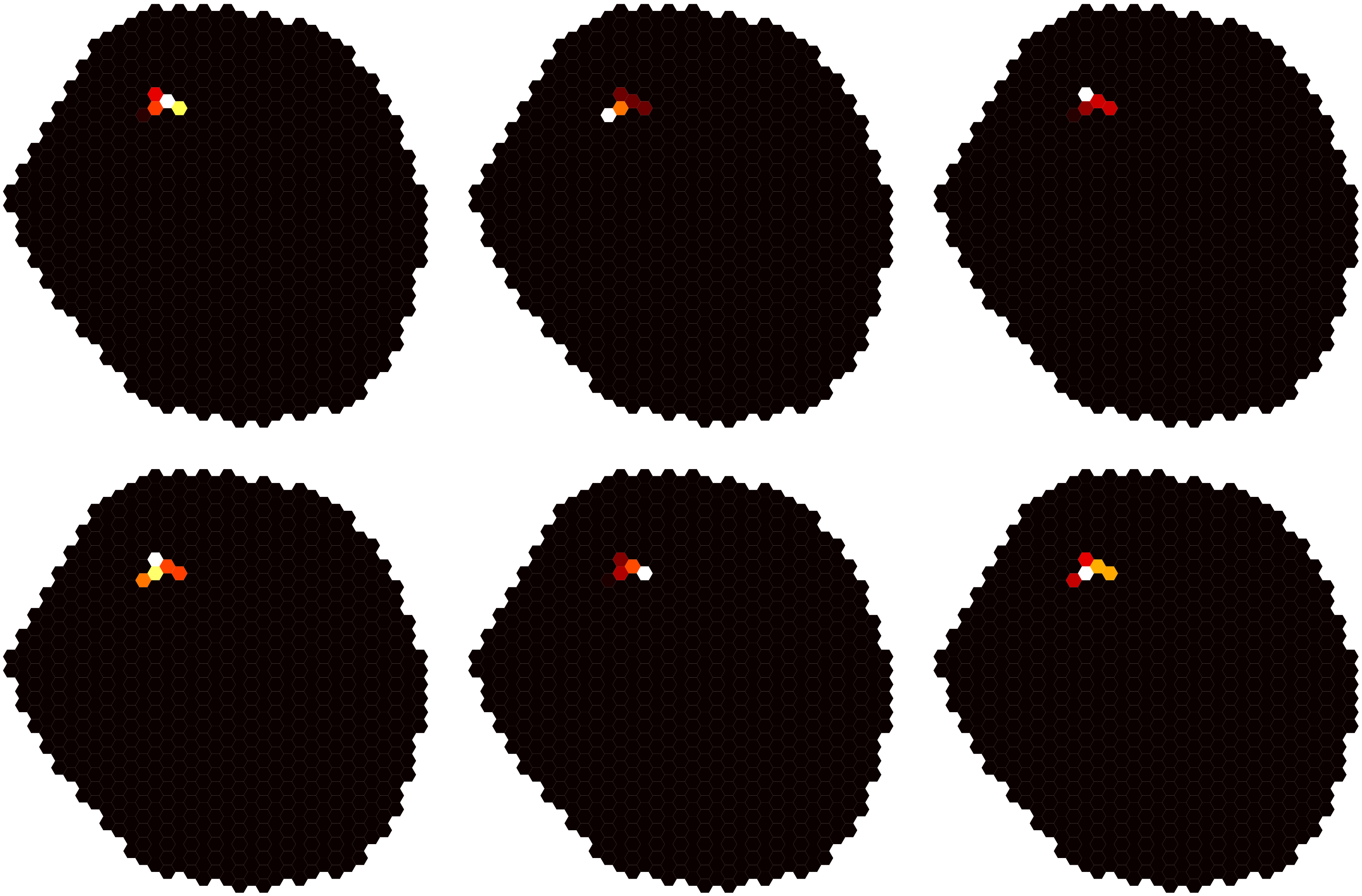

### Dm6-level5-11.pdf

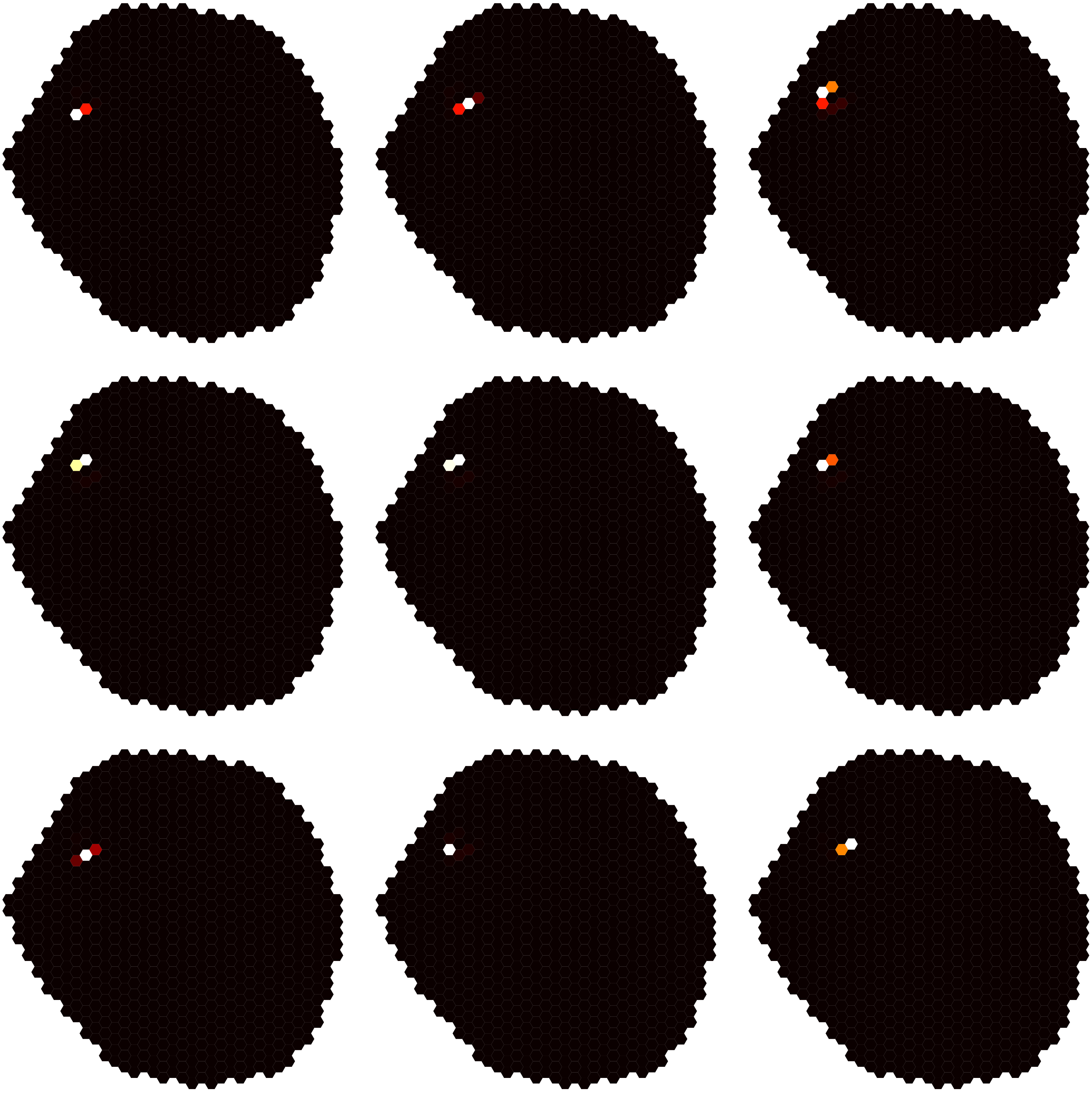

### Dm6-level5-12.pdf

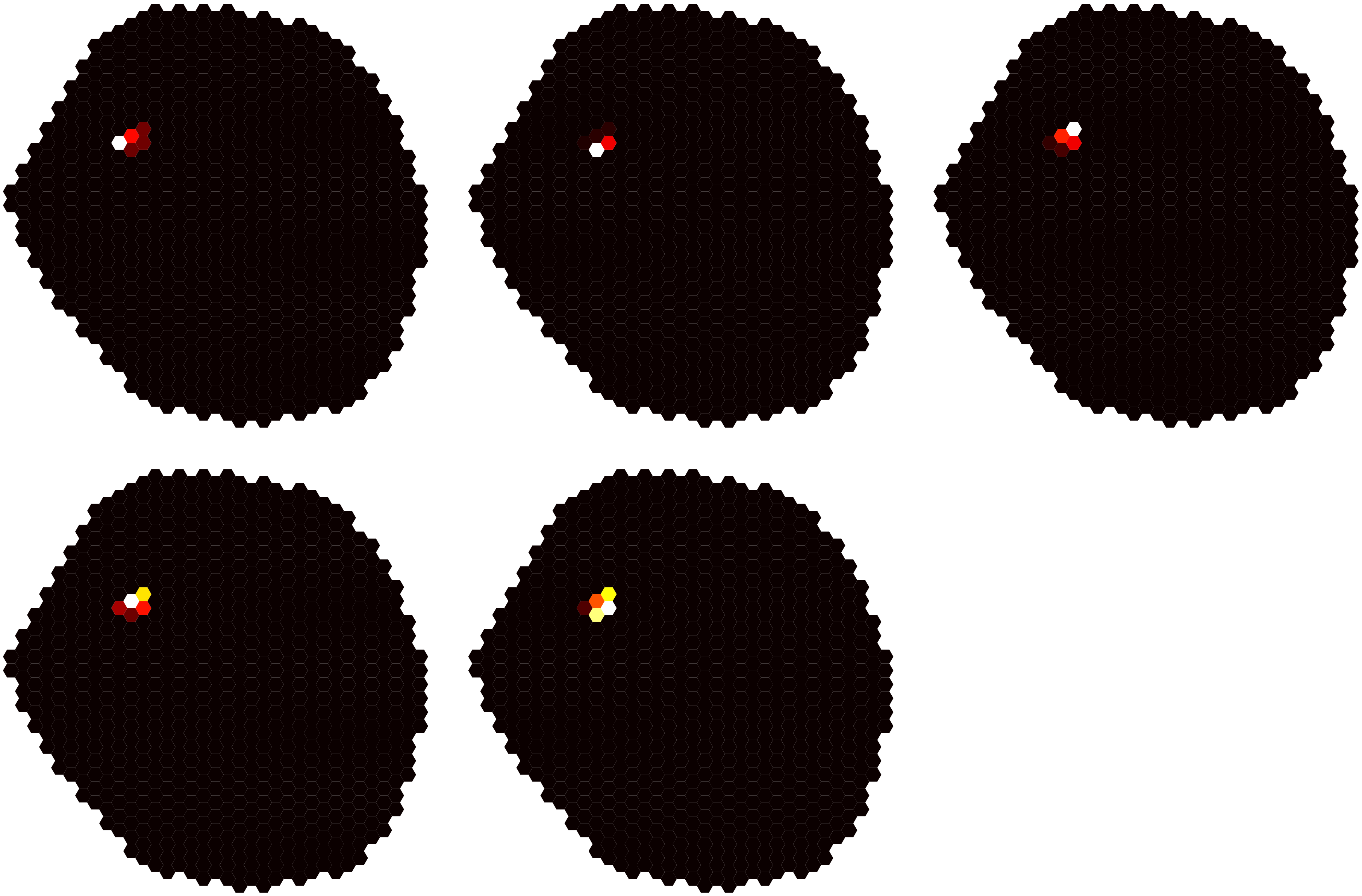

### Dm6-level5-13.pdf

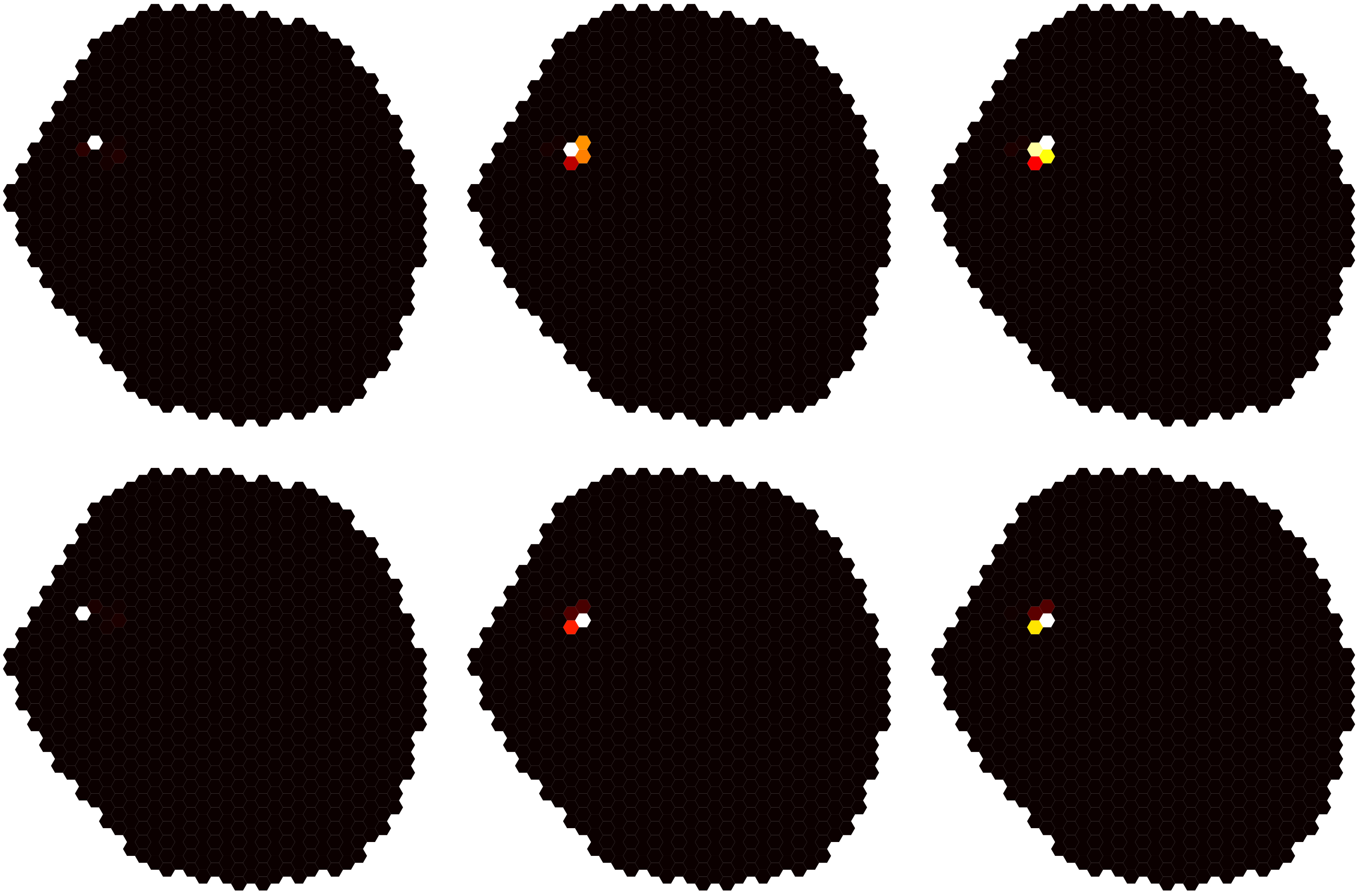

### Dm6-level5-14.pdf

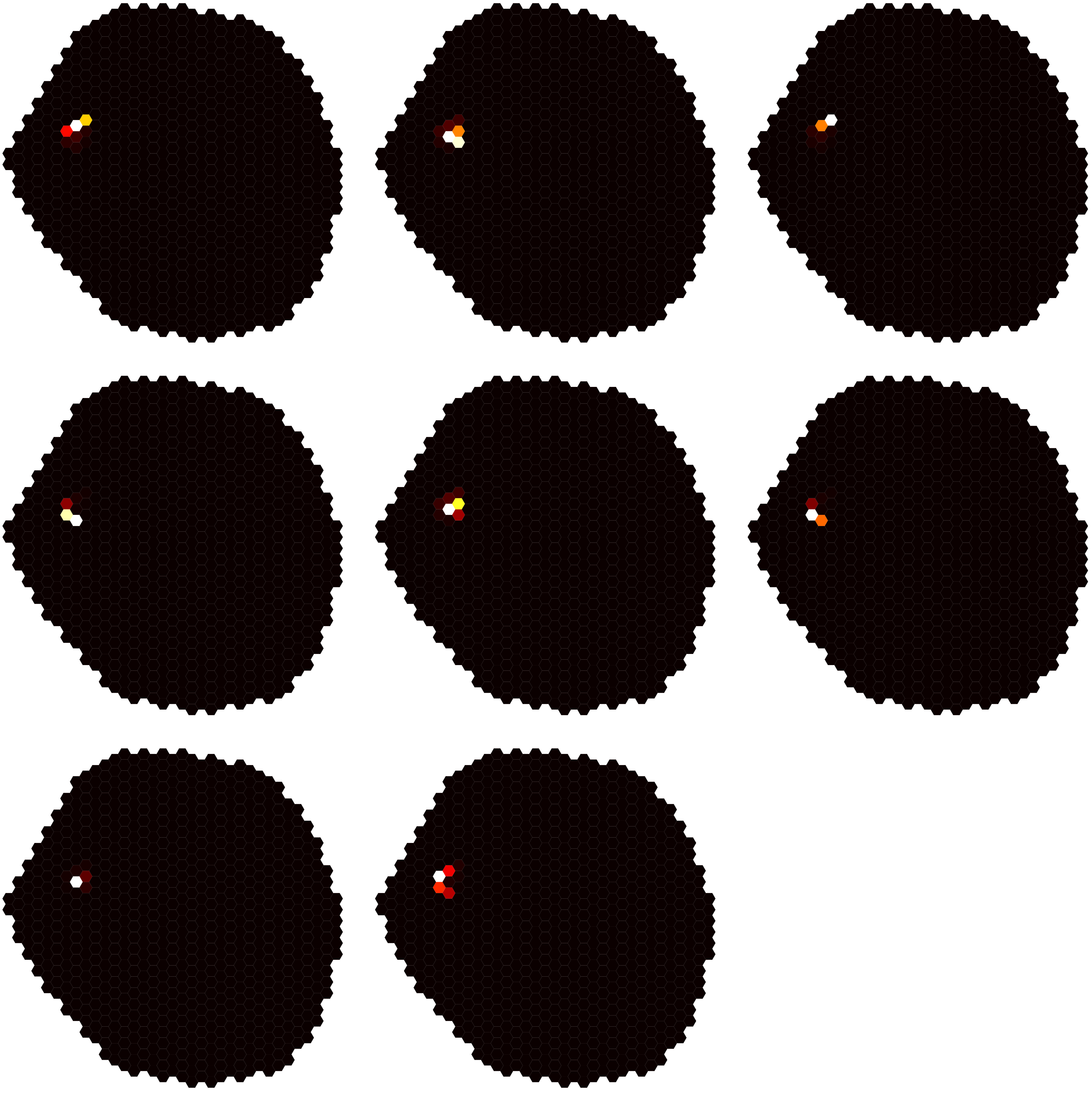

### Dm6-level5-15.pdf

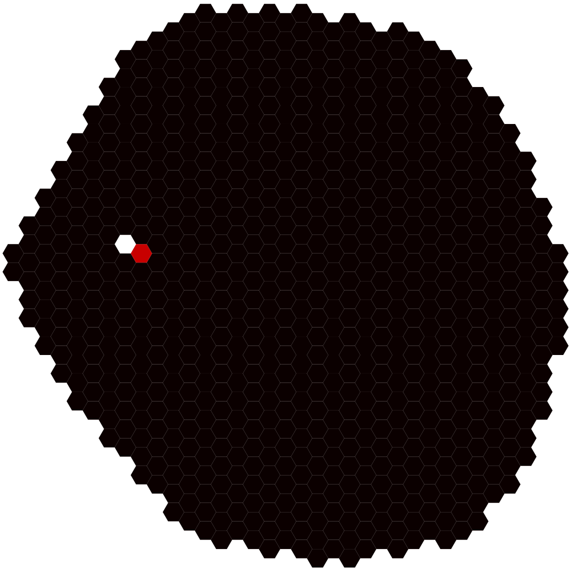

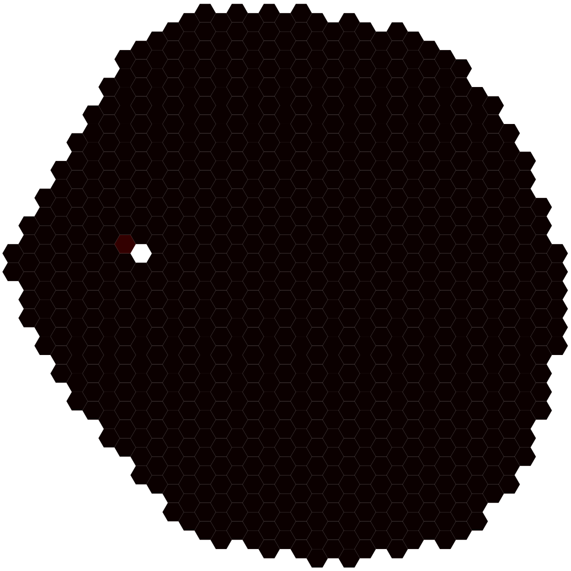

### Dm6-level5-16.pdf

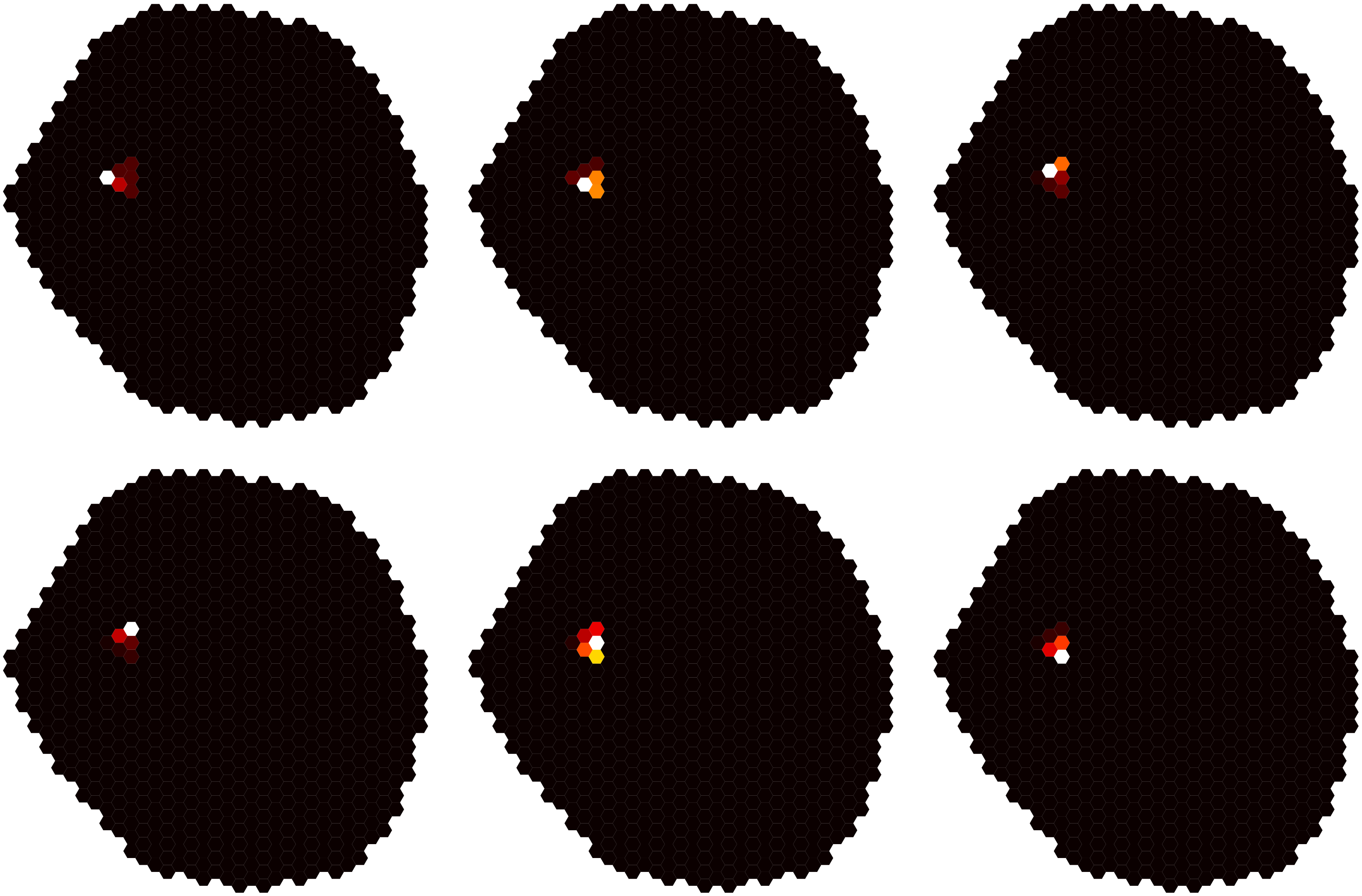

### Dm17-level5-1.pdf

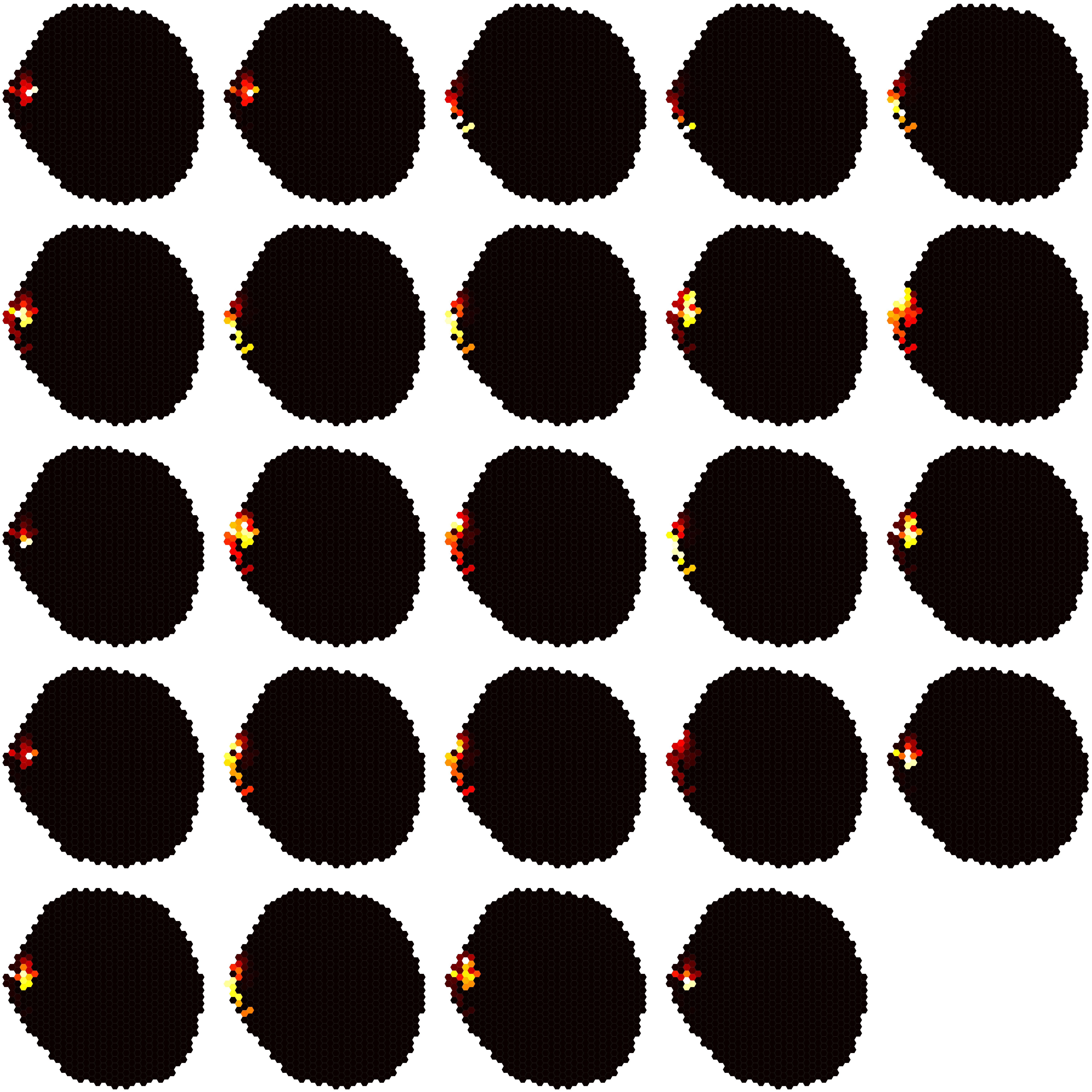

### Dm17-level5-2.pdf

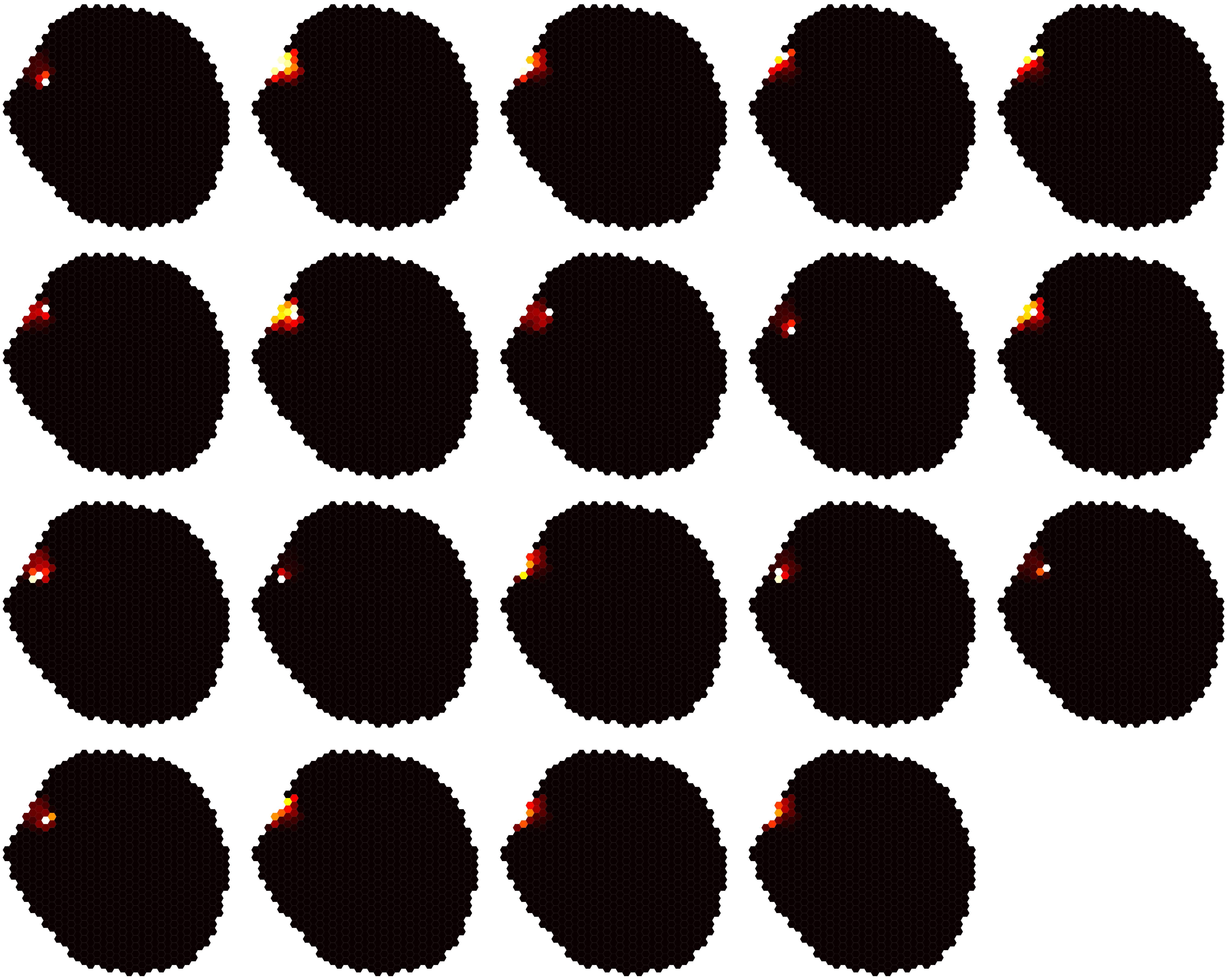

### Dm17-level5-3.pdf

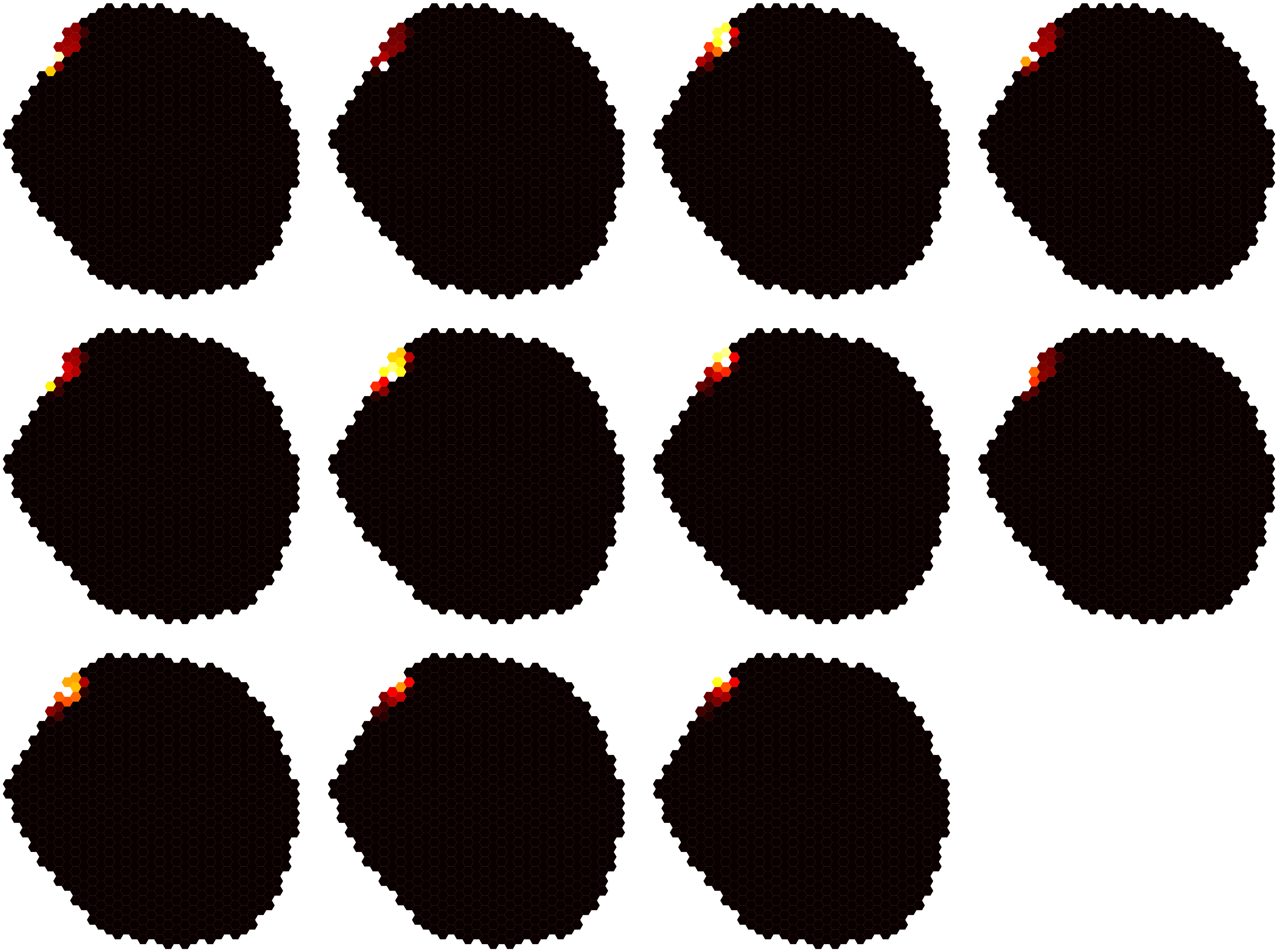

### Dm17-level5-4.pdf

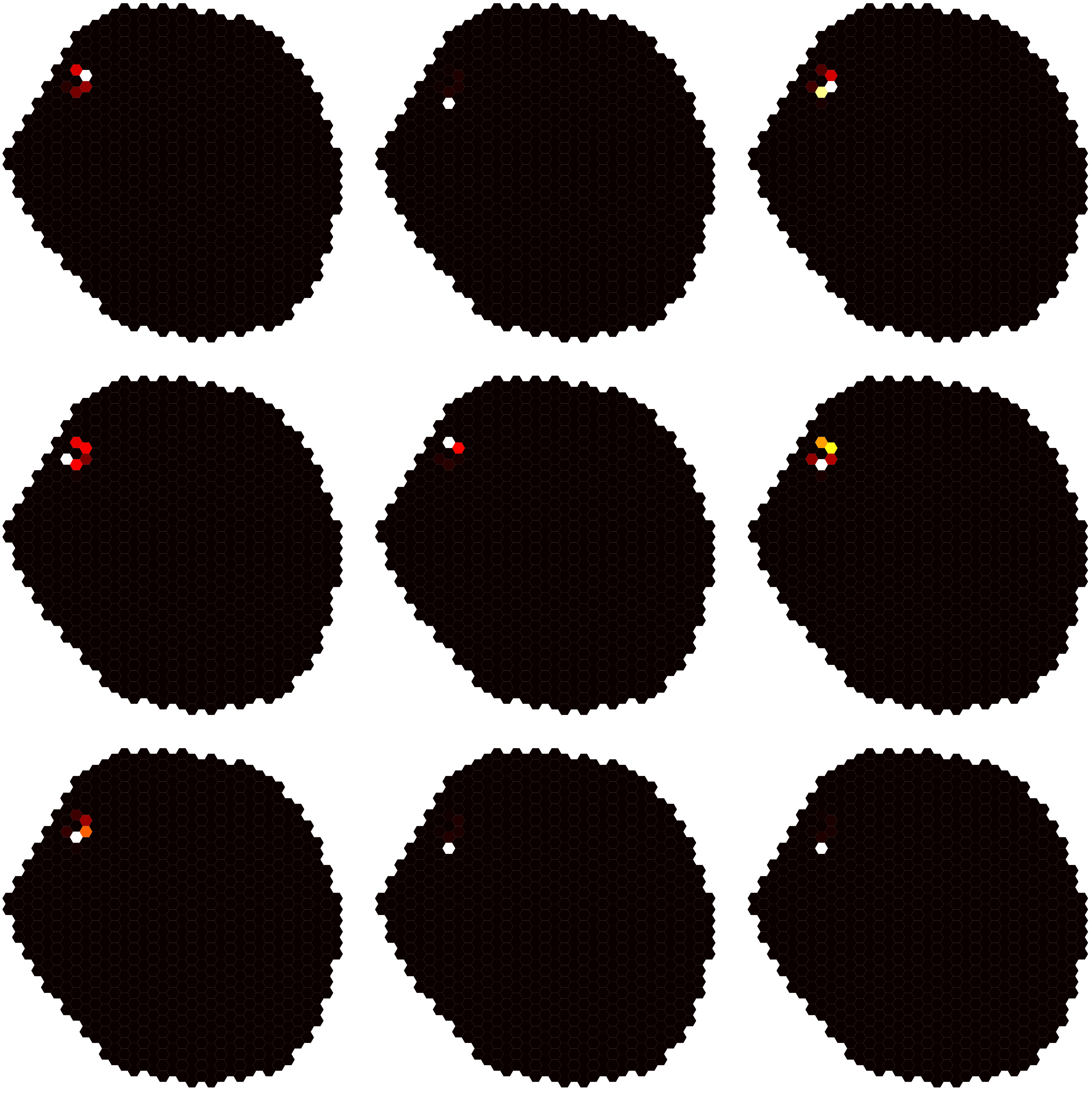

### Dm17-level5-5.pdf

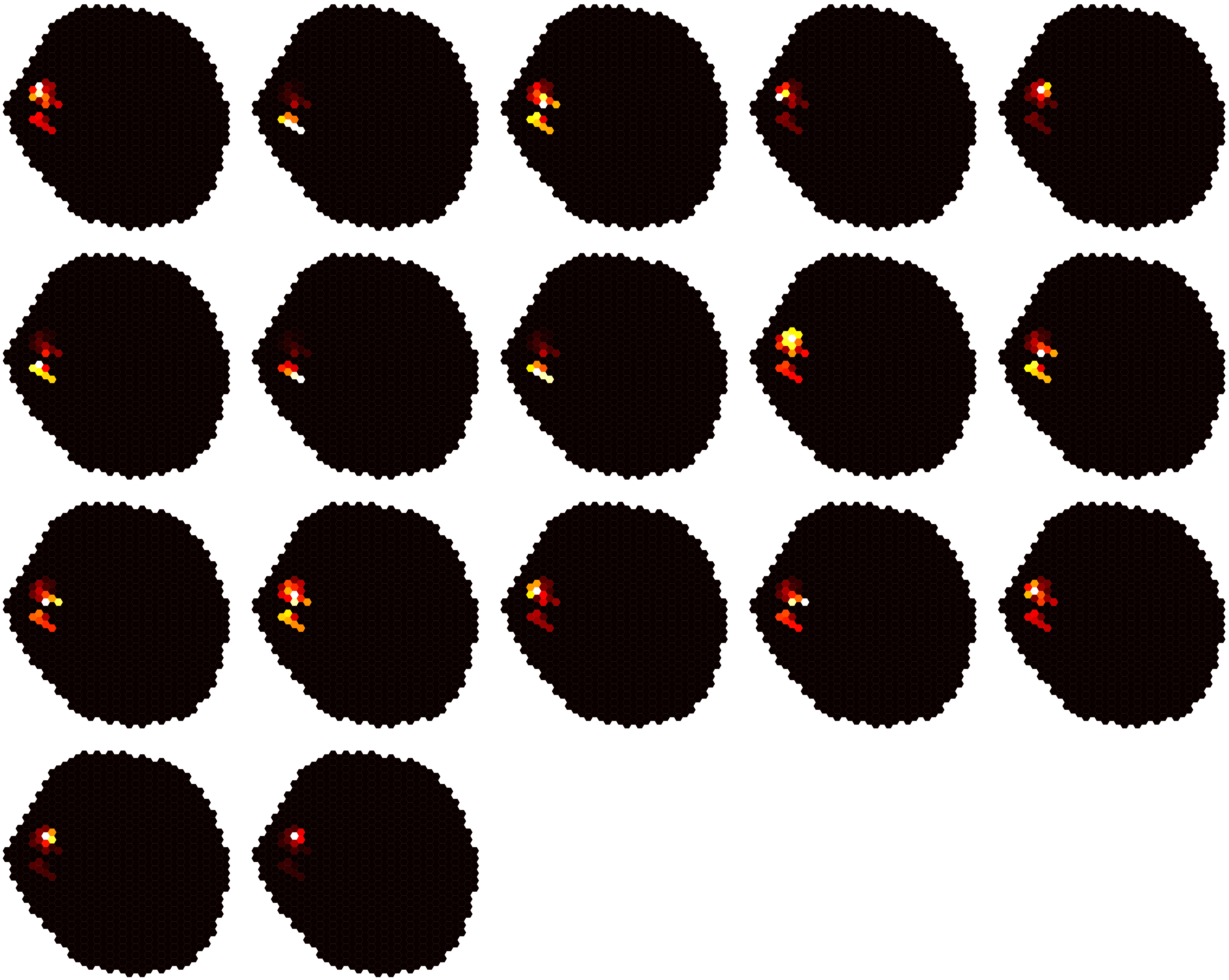

### Dm17-level5-6.pdf

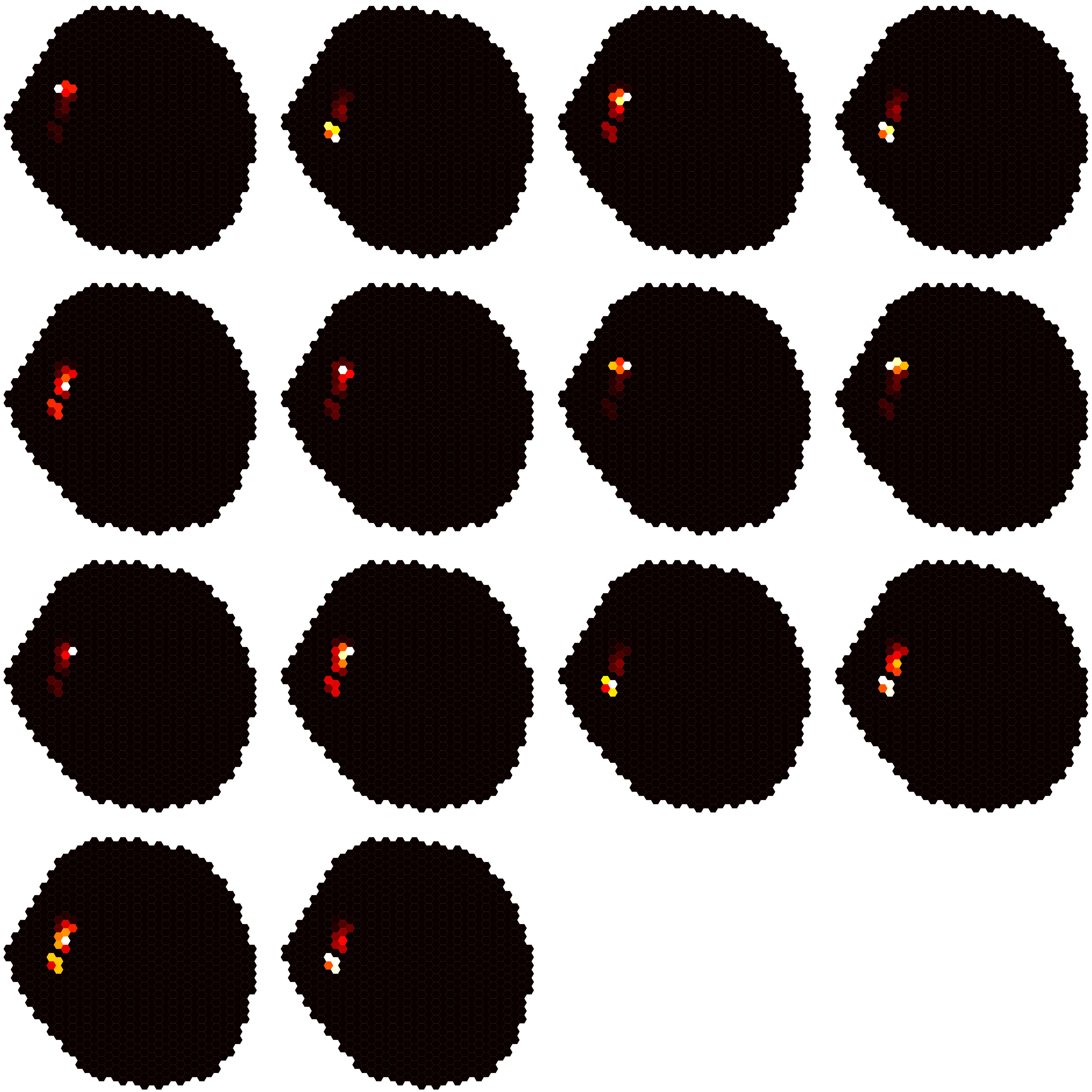

### Dm17-level5-7.pdf

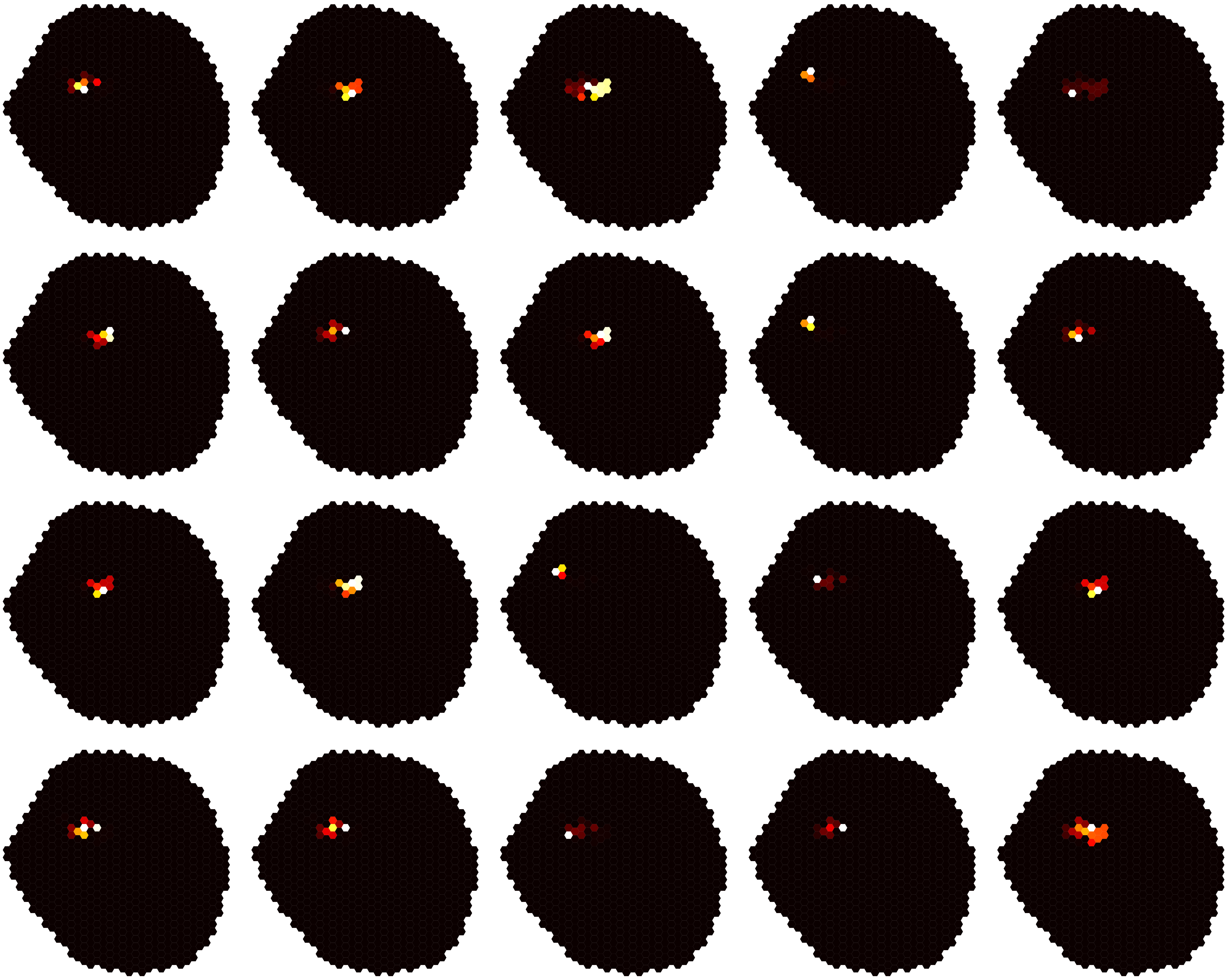

### Dm17-level5-8.pdf

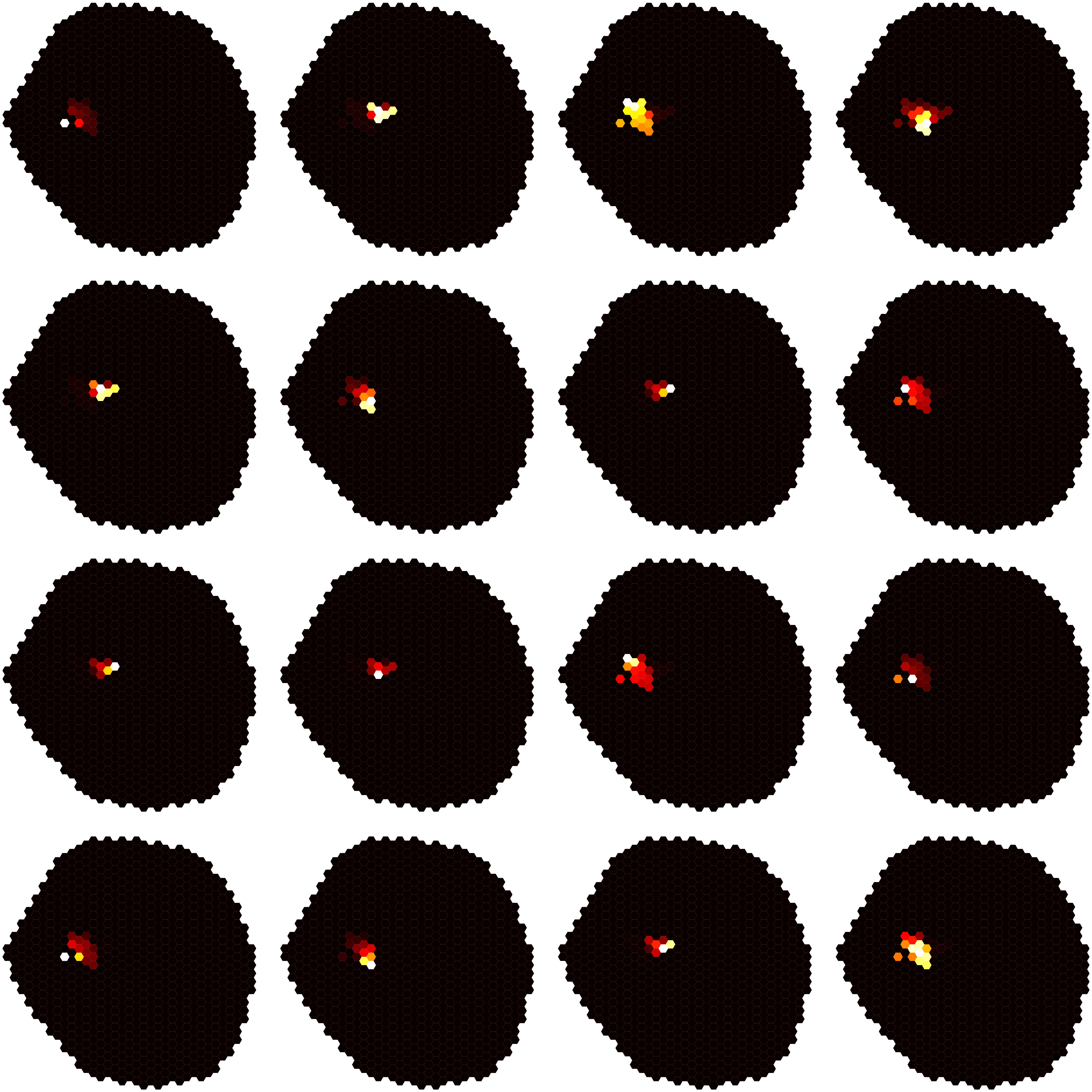

### Dm17-level5-9.pdf

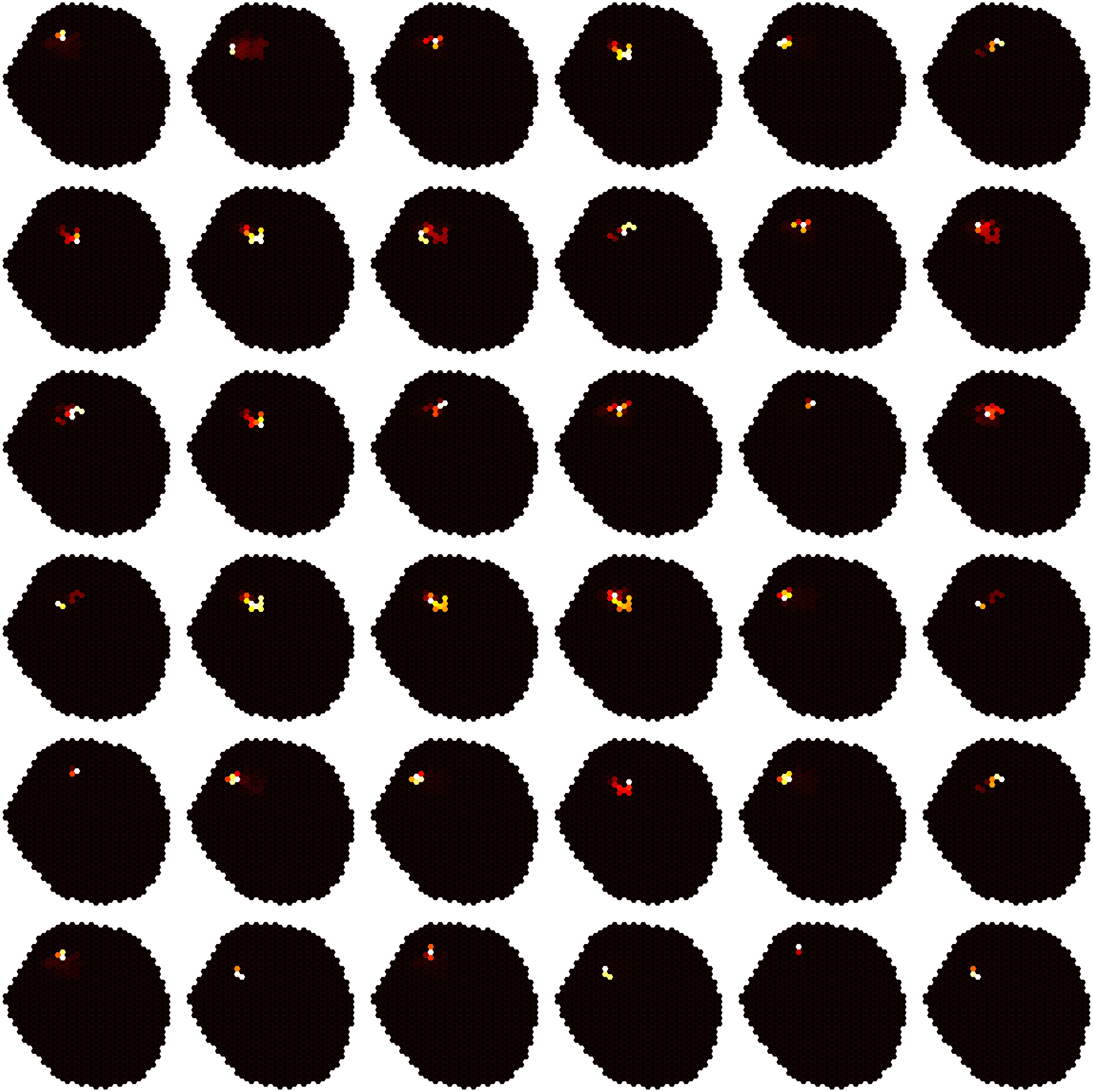

### Dm17-level5-10.pdf

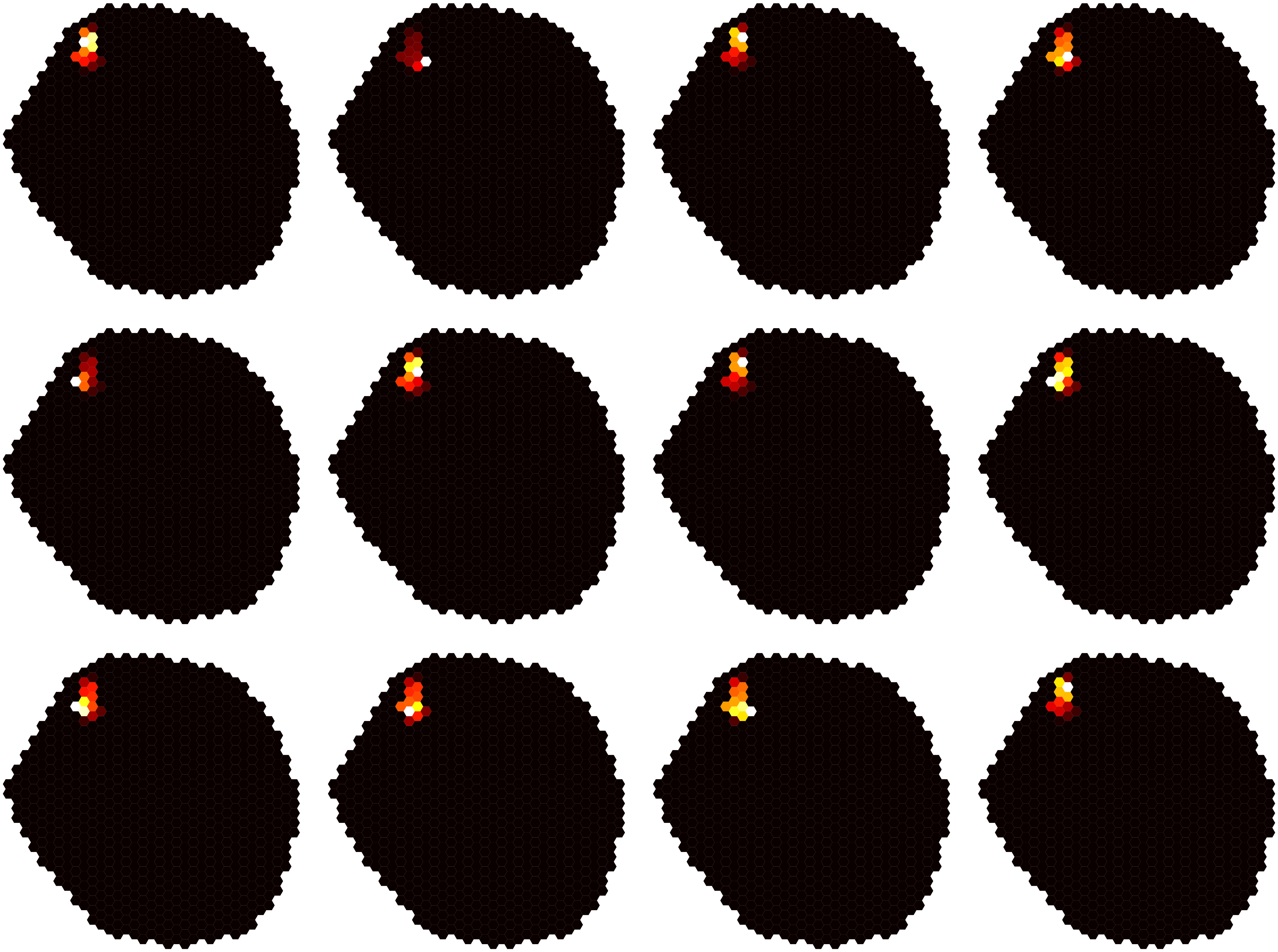

### Dm17-level5-11.pdf

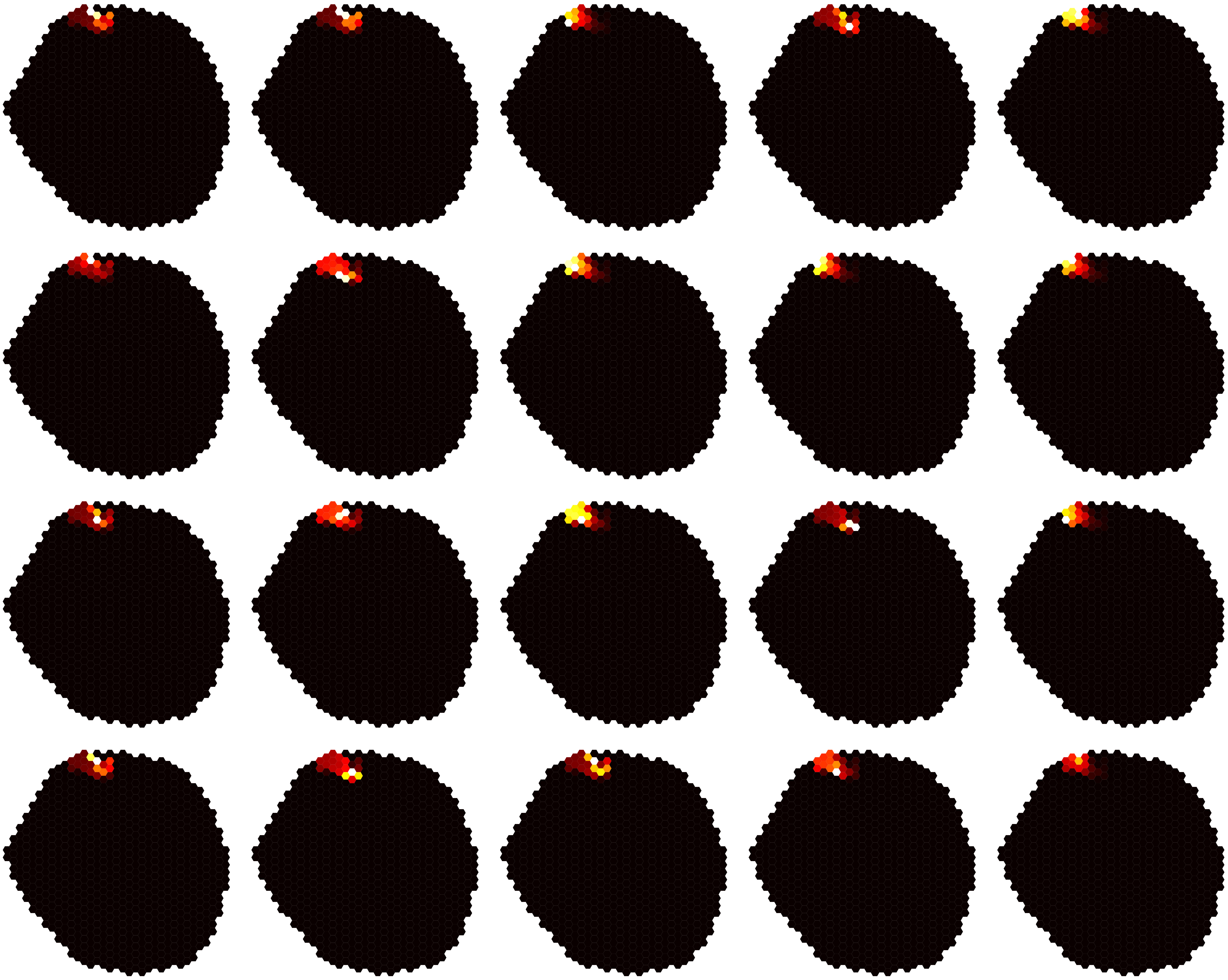

### Dm17-level5-12.pdf

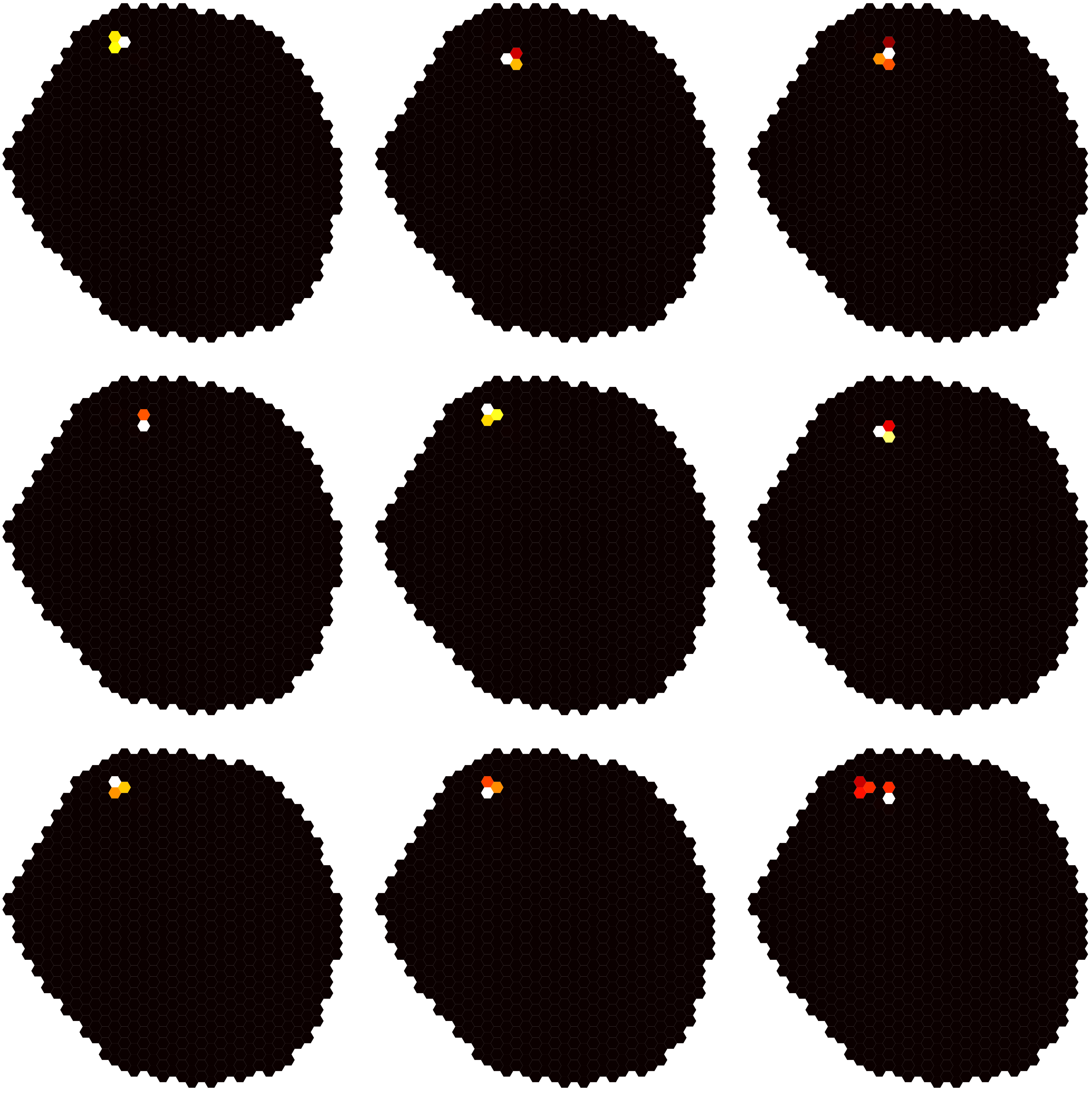

### Dm17-level5-13.pdf

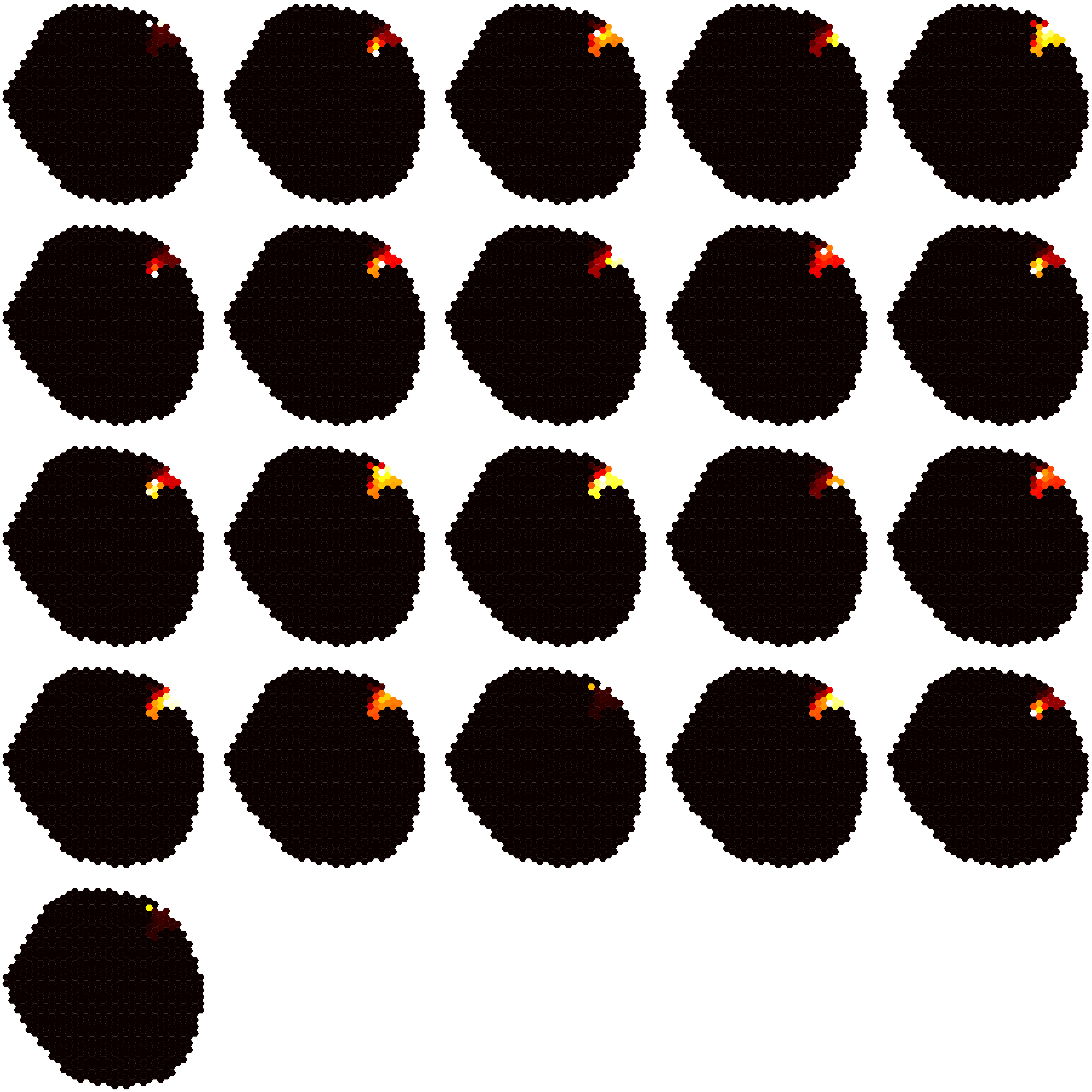

### Dm17-level5-14.pdf

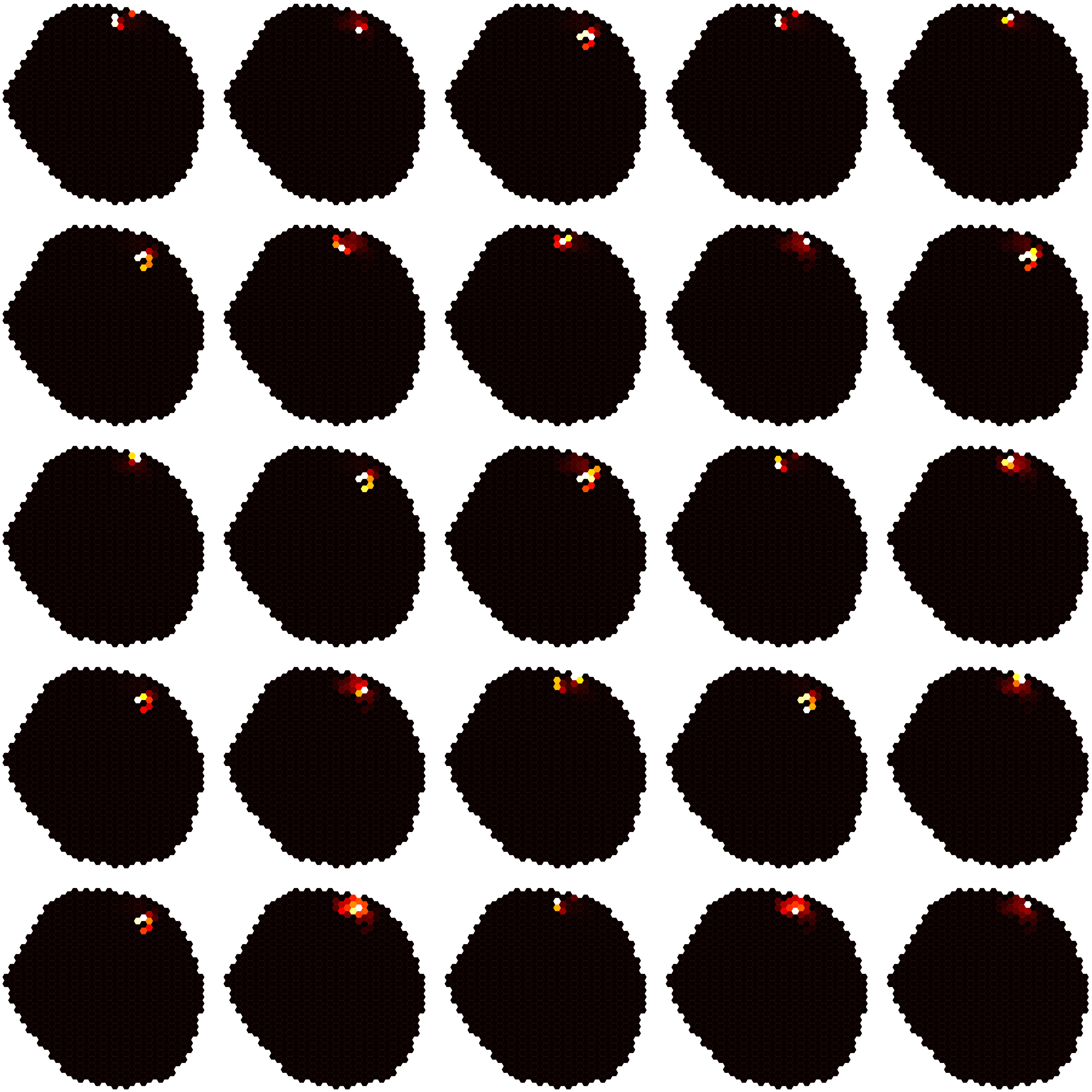

### Dm17-level5-15.pdf

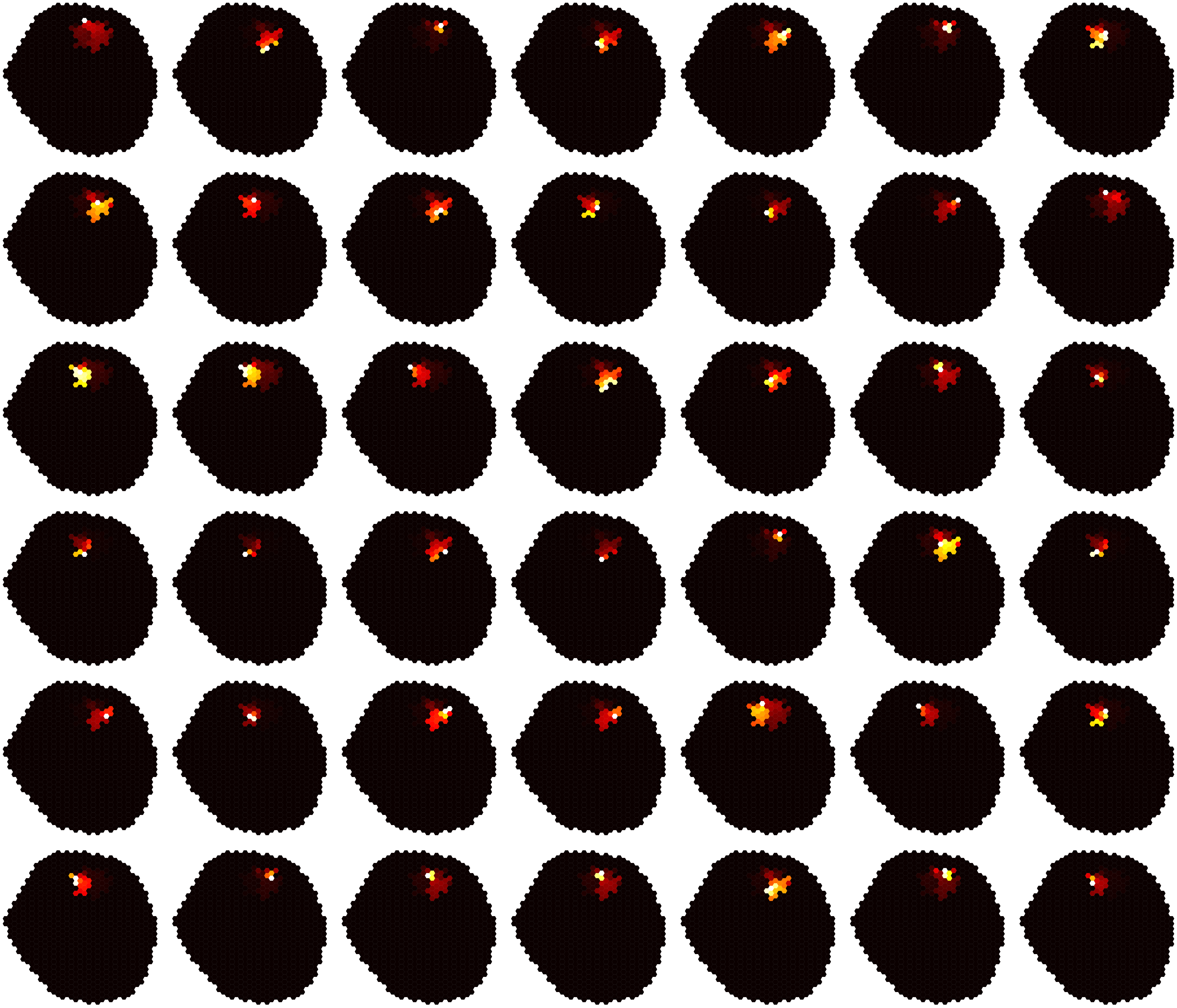

### Dm17-level5-16.pdf

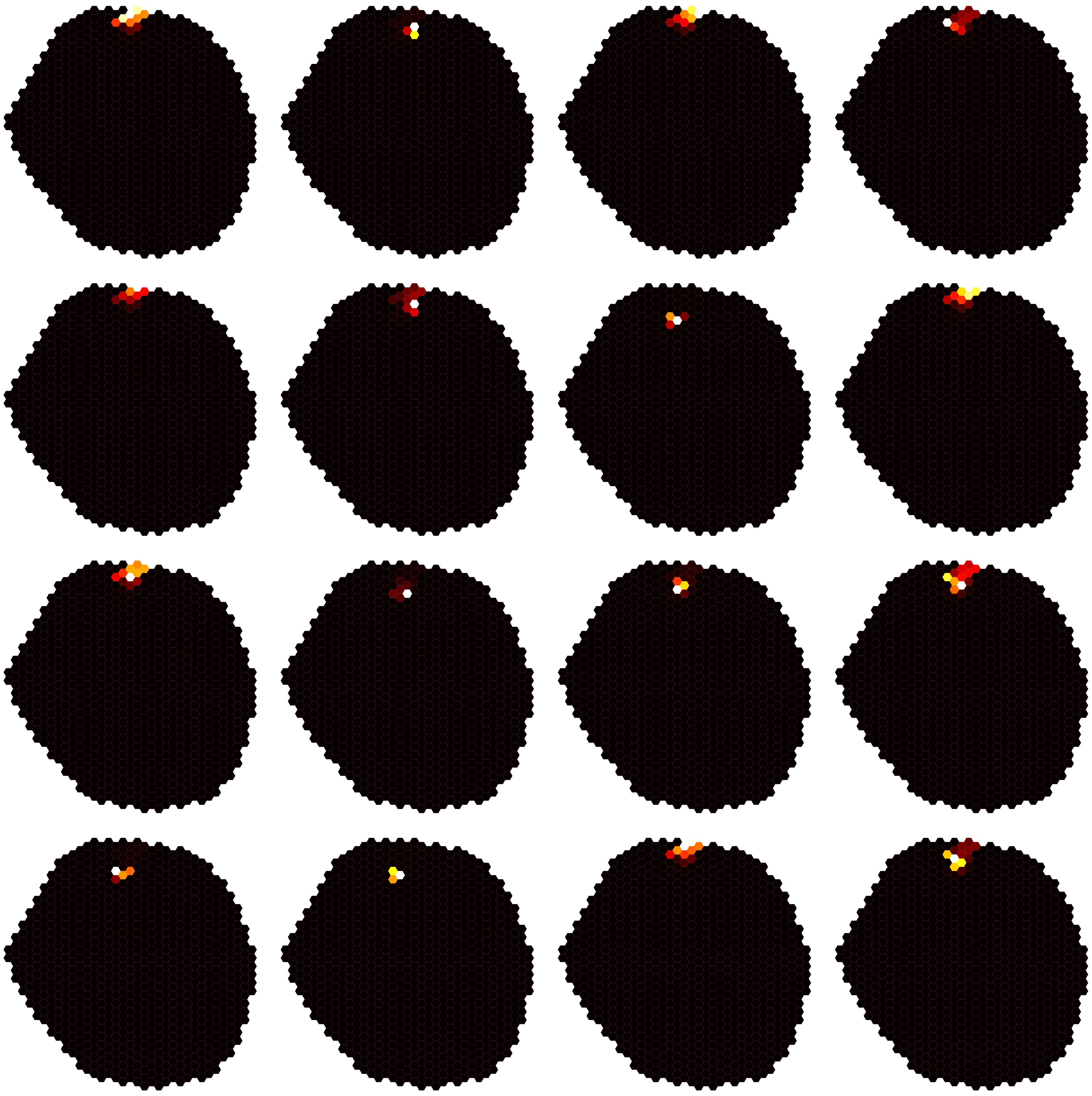
